# Computer experimentation on *E. coli* ammonium transport and assimilation reveals mechanisms for energy coupling, balanced futile cycling, and robust growth

**DOI:** 10.64898/2026.05.09.723968

**Authors:** Kazuhiro Maeda, Hiroyuki Kurata, Arnaud Javelle, Hans V. Westerhoff, Fred C. Boogerd

## Abstract

Nitrogen is essential for all life forms, and microorganisms prefer ammonium as a nitrogen source. Due to the low affinity of glutamine synthetase (GS) for ammonium, *E. coli* must maintain high intracellular ammonium (NH_4_^+^) concentrations to sustain its rapid growth. Under ammonium limitation, *E. coli* imports ammonium through the transporter AmtB and incorporates it into glutamine by using GS. On the basis of structural and mutagenesis information, mechanisms have been proposed for the transport of ammonia (NH_3_) and protons by AmtB through spatially (partly) separate routes. These mechanisms do not explain the required *coupling* between proton and ammonia transports. How does the membrane potential push the ammonia inward so as to attain high concentrations near GS? We here compare six candidate kinetic models of *E. coli* ammonium transport and assimilation in terms of how they reproduce experimental data from the literature: three variants of the ‘electro-binding model’ in which the membrane potential affects AmtB–NH_4_^+^ binding, and three variants of the ‘electro-flipping model’ in which it influences the conformational flip of the transporter. The computer simulations decide that the electro-binding models are 28 times more plausible than the electro-flipping models and suggest that the transmembrane electric potential affects AmtB–NH_4_^+^ binding from the cytoplasmic side. The addition of kinetic and thermodynamic features to existing structural information plus our requirement of an explanation of the coupling, suggest a new spatiotemporal mechanism of coupling of ammonia and proton flows in AmtB. Further simulations show that GS and AmtB regulation is coordinated via both the uridylyltransferase/uridylyl-removing enzyme (UTase) and 2-oxoglutarate binding, allowing the cell to minimize futile cycling while maintaining rapid growth. The free energy cost of transport-related futile cycling exceeded that of the GS reaction itself. Moreover, AmtB enabled robust growth under varying ammonium concentrations and pH levels, albeit at a cost of futile cycling that became substantial at low ammonium. These findings highlight the crucial roles of GS and AmtB in *E. coli*’s adaptations and provide new insights into the trade-off mechanism between nutrient acquisition and energy efficiency.

## Introduction

Nitrogen is an essential element for all life forms and available in the environment as part of various inorganic and organic compounds. Most microorganisms prefer ammonium as their nitrogen source, which consists of two molecular species, NH_4_^+^ (ammonium) and NH_3_ (ammonia). In the present study, we use NH_x_ as a general term when the specification of the molecular species is unwanted or unnecessary. The protonation of NH_3_ to NH_4_^+^ occurs instantaneously. In the neutral pH range, NH_4_^+^ is the predominant species (pKa = 8.95 at T = 37 °C). The uptake of NH_x_ is not a problem as long as the compound is amply present in the direct environment of the microbial cells, because, like water, the unprotonated species NH_3_ can permeate the microbial phospholipid membranes readily. However, if the environmental availability of NH_x_ becomes limited, unmediated NH_3_ diffusion through the membrane is no longer sufficient to supply the central nitrogen metabolic machinery with NH_x_^1.^

When confronted with ammonium limitation, *E. coli* produces the transporter AmtB ^1^. The consensus in the literature is that AmtB has high-specificity and high-affinity binding sites for NH_4_^+^ (not NH_3_) on the periplasmic side of the transporter. However, the question of whether ammonium is transported as NH_4_^+^ or NH_3_ to the cytoplasm has been the subject of vivid debate over the last 40 years ^2-4^ (reviewed in refs ^5-11^). In a hypothesis paper ^12^, co-authored by two of the authors of the present study, it was argued that the transport of ammonium must be an active process to ensure sufficient accumulation of intracellular NH_4_^+^; NH_4_^+^ transport by AmtB may be called ‘active transport’ of NH_x_ as it consumes proton-motive force, i.e., one proton per nitrogen atom transported. It is not so much the uptake rate that matters, but the intracellular ammonium concentration, because GS has a relatively low affinity for ammonium, characterized by a K_m_ of 100 μM NH_4_^+^ ^13^. Consequently, the transported substance should be the cation NH_4_^+^ (or NH_3_ strictly coupled to H^+^, but not NH_3_ alone) ^12^. Then the transmembrane electric potential (Δ*Ψ), which is neg*ative inside by more than 120 mV, would pull the NH_4_^+^ (but not the neutral NH_3_) through the AmtB into the cytoplasm up to a more than 100-fold accumulation. If NH_3_ as such were transported by AmtB, the intracellular concentration of NH_4_^+^ would be as low as or even lower than its extracellular concentration. In an earlier study, we developed a detailed kinetic model of the *E. coli* ammonium transport and assimilation network ^14^. By model-based comparison of the active and passive AmtB transport mechanisms, we revealed that it is highly plausible that AmtB should be an ‘active’ (electrogenic) transporter that transports NH_4_^+^ (or NH_3_ + H^+^) against its concentration gradient. The evidence in favor of passive or active AmtB-mediated transport has been discussed in the review by van Heeswijk, Westerhoff and Boogerd ^1^. Two more recent experimental papers by Javelle et al. ^15-16^ also concluded that AmtB in *E. coli* is an electrogenic transporter.

AmtB-mediated ammonium uptake entails a great deal of expense ^12,14^: The accumulation of ammonium inside the cells leads to outward NH_3_ diffusion through the cytoplasmic membrane, followed by the re-uptake of NH_4_^+^. This cycle, known as the *futile cycle*, might be repeated several times before NH_4_^+^ is finally assimilated into glutamine, requiring extra free-energy investment with each cycle. To mitigate this potential free-energy expense, *E. coli* uses the regulatory protein GlnK under ammonium-limited conditions. The *glnK* gene, discovered by the late Dr. Wally C. van Heeswijk ^17^, forms an operon with the *amtB* gene in multiple prokaryotes ^18^. The cytoplasmic GlnK protein forms a complex with membrane-bound AmtB, inhibiting its transporter function. This complex formation is regulated by specific modifications of GlnK ^19-21^, allowing the adjustment of AmtB activity to meet the cellular need for ammonium and minimizing the associated ammonium futile cycling ^22^.

Despite these previous insights, several important questions remain unanswered. First, it is unclear how precisely the transmembrane electric potential affects AmtB-mediated NH_4_^+^ transport—does it primarily influence AmtB–NH_4_^+^ binding or the conformational transitions of AmtB? Second, what is the mechanism of the coupling between the NH_3_ and the proton transport; is the proton really spatially separable from the NH_3_, as proposed by Williamson et al. ^16^, without the coupling being compromised? Third, how are GS and AmtB coordinated to balance efficient ammonium uptake and the Gibbs energy loss due to futile cycling? Fourth, are there conditions, even at low ammonium concentrations, under which AmtB is not required, i.e., the ΔAmtB mutant can grow just as well as the wild type? After all, without AmtB, there would be no free energy loss due to ammonium futile cycling. Finally, are ammonium transport and assimilation robust to environmental and physiological fluctuations such as changes in pH? In wet laboratory experiments, changing one parameter inevitably affects other parameters, e.g., through alterations of gene expression, making it difficult to measure specific effects. Computer simulations can help answer questions of this type because they enable one to change one parameter or one process at a time. Our experimentally validated kinetic model of the *E. coli* ammonium transport and assimilation network ^14^ should be suitable for this purpose.

In the present study, we first compare two classes of distinct mechanistic hypotheses for AmtB-mediated NH_4_^+^ transport and find that the model classes assuming that the transmembrane electric potential influences AmtB– NH_4_^+^ binding from the cytoplasmic side are more plausible. This leads us to a new mechanism for AmtB’s coupling of proton to NH_3_ transport. The most plausible models further reveal that the coordinated regulation of GS and AmtB by regulatory proteins such as GlnK, GlnB, UTase, and ATase minimizes futile cycling while maximizing growth under various environmental and physiological conditions.

## Results

### Model description

In this study, we employed the kinetic model ^14^ for the ammonium transport and assimilation network in *E. coli* that ranked best vis-à-vis the description of three pertinent and precise experimental studies ^21-23^. The network structure is illustrated in **Fig. 1**, where the CADLIVE notation ^24-28^ is used. Briefly, the model contains the AmtB-mediated NH_4_^+^ transport (at rate *v*_*amtb*_), passive diffusion of NH_3_ across the cytoplasmic membrane (at rate *v*_*diff*_), and regulation of AmtB activity by GlnK. This model also contains glutamate dehydrogenase (GDH), glutamate synthase (GOGAT), and glutamine synthetase (GS) as well as proteins such as GlnB, UTase, and ATase that regulate GS. The specific growth rate (μ) is modeled as a function of intracellular glutamate and glutamine concentrations [Eqs. (30)-(31) in Methods]. The total concentrations of AmtB, GlnK, and GS are adjusted using experimental data on transporter activity ^22^ (see Section 1 of Supplementary Notes). This model does not account for carbon metabolism and assumes that carbon is always available in excess relative to NH_x_. 2-Oxoglutarate concentration is modeled as a function of internal NH_4_^+^ concentration (see Section 2 of Supplementary Notes). We modified the rate equations of the original model ^14^ for the GOGAT reaction and the AmtB-mediated NH_4_^+^ transport. This allows us to expand the range of culture conditions and to investigate the mechanism of AmtB-mediated NH_4_^+^ transport. The final model consists of 13 balance equations, 23 rate equations, and 118 model parameters (**Tables S1-S4** in Supplementary Tables).

**Fig. 1.**
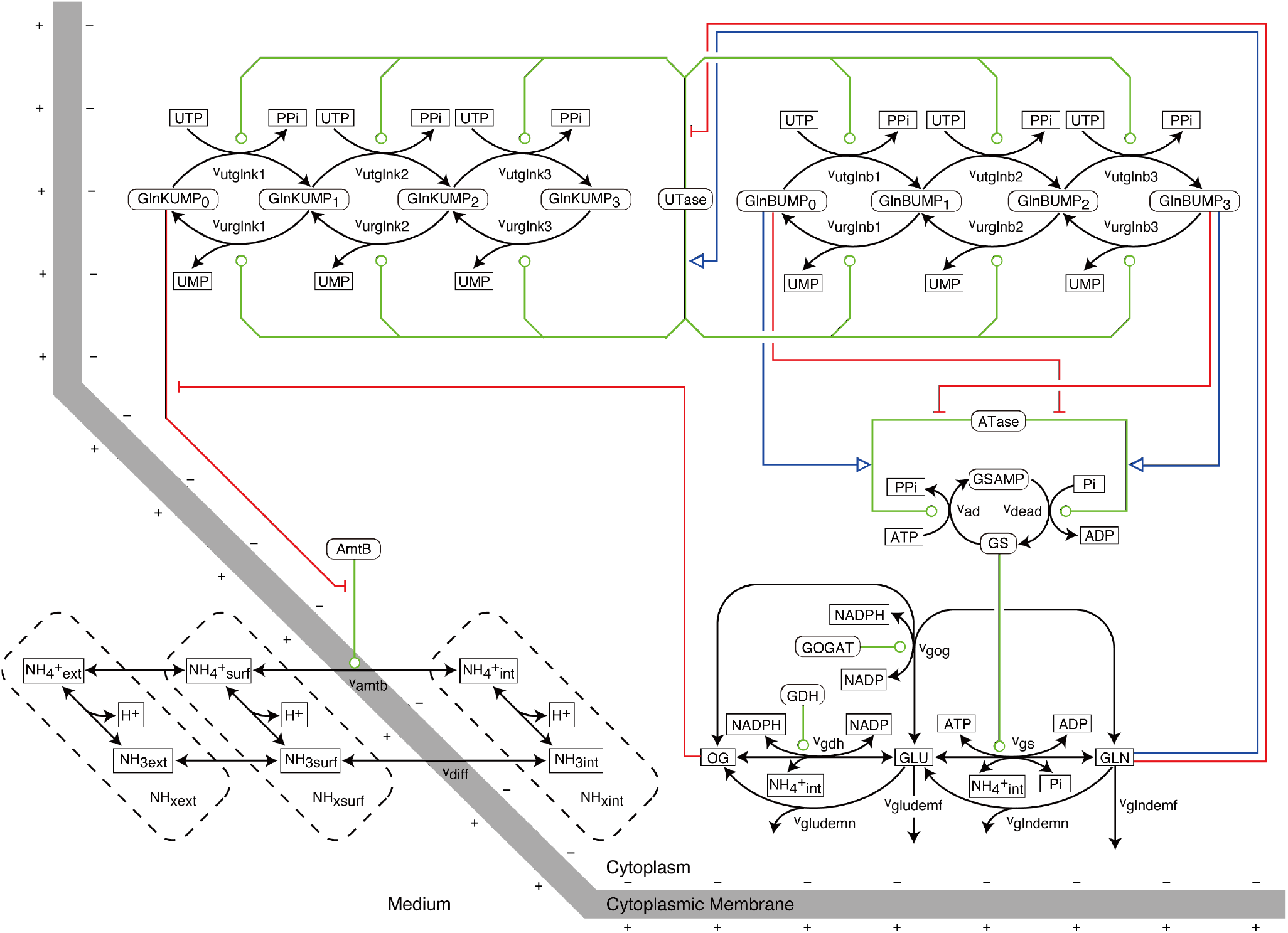
The *E. coli* ammonium transport and assimilation network. Scheme of the *E. coli* ammonium transport and assimilation network. For simplicity, CADLIVE notation was used. Blue lines: activation, red lines: inhibition, green lines: catalytic action, black lines: chemical conversion or physical displacement. The thick grey line indicates the cytoplasmic membrane. The following abbreviations and nomenclature were used. GS: glutamine synthetase (dodecamer), GDH: glutamate dehydrogenase (monomer), GOGAT: glutamate synthase (heterodimer), ATase: adenylyltransferase/adenylyl-removing enzyme (monomer), UTase: uridylyltransferase/uridylyl-removing enzyme (monomer), GlnB and GlnK: nitrogen regulatory proteins (trimers), AmtB: ammonium transporter (trimer), OG: 2-oxoglutarate, GLU: glutamate, GLN: glutamine. int: intracellular, surf: surface, ext: extracellular far away from the cell. In the model, the concentrations of multimeric proteins are treated based on their multimeric forms. Figure adapted and modified from Maeda et al., npj Syst Biol Appl, 2019, published under a Creative Commons Attribution 4.0 International License (http://creativecommons.org/licenses/by/4.0/).

### How is AmtB-mediated NH_4_^+^ transport energized by the transmembrane electric potential?

#### Two model classes: potential-dependent ammonium binding affinity or potential-dependent conformational flip

The reaction scheme of AmtB-mediated NH_4_^+^ transport across the cytoplasmic membrane is shown in **Fig. 2**. We employed the following rate equation for the transport:

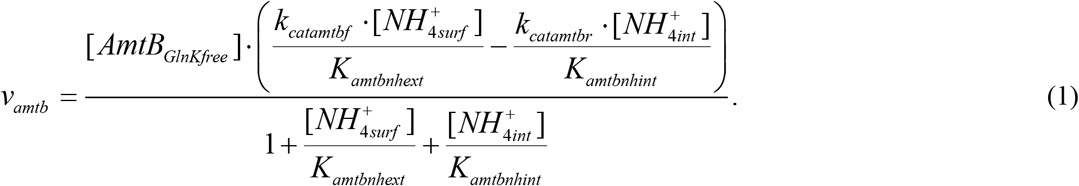

**Fig. 2.**
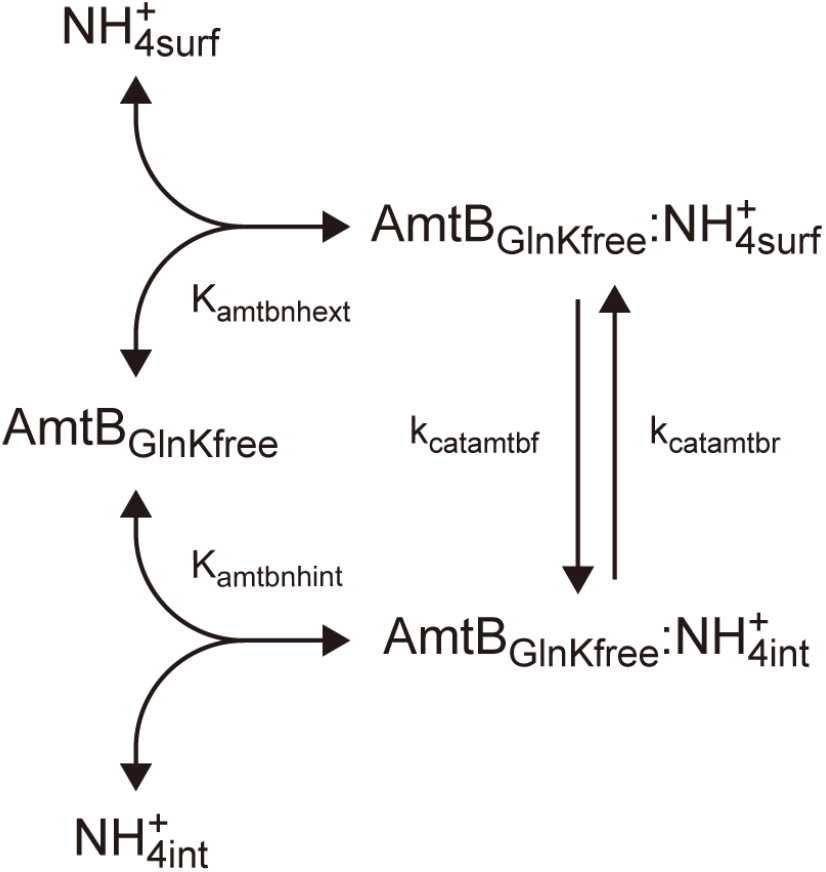
Reaction scheme of active transport across the cytoplasmic membrane. First, NH_4_^+^ binds to AmtB that is not associated with GlnK at the membrane surface. The AmtB–NH_4_^+^ complex is assumed to exist in two conformational states: one open to the outside and the other open to the inside. After binding, NH_4_^+^ is released into the cytoplasm.

Here, [*AmtB*_*GlnKfree*_] represents the concentration of AmtB that is not bound to GlnK (i.e., the active form of AmtB). [*NH* ^+^] and [*NH* ^+^] denote the NH ^+^ concentration at the cell surface and inside the cell, respectively. *k*_*catamtbf*_ and *k*_*catamtbr*_ are the rate constants at which the binding site for NH_4_^+^ flips in the inward or outward direction, respectively, taking bound NH_4_^+^ along; this may correspond to the ammonium moving through the Phe gate ^10^. *K*_*amtbnhext*_ and *K*_*amtbnhint*_ represent the NH_4_^+^ dissociation equilibrium constants of the NH_4_^+^ binding site when it is accessible to periplasmic-surface NH_4_^+^ (‘S1’) or intracellular NH_4_^+^ (presumed ‘S2’), respectively ^9^ . A positive *v*_*amtb*_ value indicates NH_4_^+^ is on average moving from the extracellular to the intracellular space, while a negative value indicates the reverse. For a full derivation of Eq. (1), see Section 7.1 of the Supplementary Information of our previous work ^14^.

At transporter equilibrium (*v*_*amtb*_ = 0), the ratio of the internal to the external NH_4_^+^ is given by:

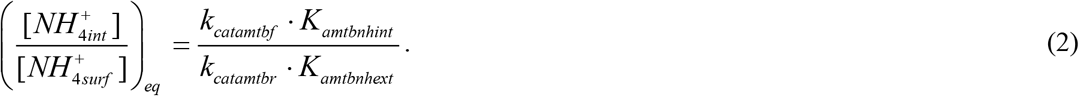

However, the four parameters in Eq. (2) are not quite independent. At transporter equilibrium, the electrochemical potential difference across the cytoplasmic membrane for NH_4_^+^ is given by:

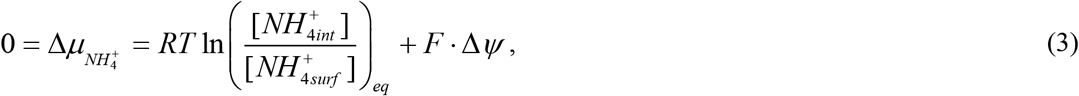

where *F* is the Faraday constant, Δ*ψ* is the transmembrane electric potential of approximately -150 mV, *R* is the gas constant, and *T* is the absolute temperature. According to Eq. (3), the theoretical accumulation factor of NH_4_^+^ (*φ*) is given by:

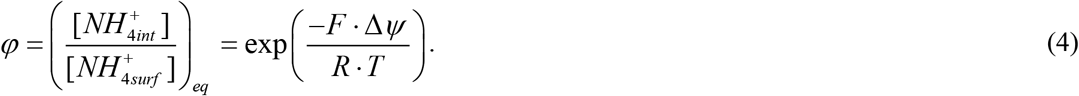

For reference, *φ =* 275 at *T* = 310 K and Δ*Ψ = -150 mV. k*_*catamtbf*_, *k*_*catamtbr*_, *K*_*amtbnhext*_, and *K*_*amtbnhint*_ must satisfy the following equation:

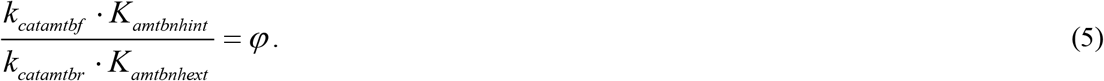

Which of these four parameters are influenced by the transmembrane electric potential? Identifying this could provide insights into the NH_4_^+^ transport mechanism of AmtB. We specifically consider two classes of hypotheses: the electro-binding and the electro-flipping hypotheses.

For the electro-binding hypotheses, we assume

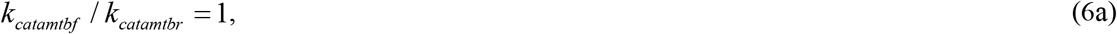

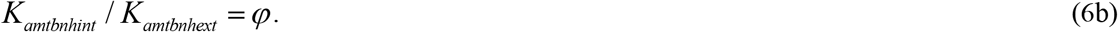

This corresponds to a scenario in which the transmembrane electric potential affects the ammonium affinity of the periplasmic and/or cytoplasmic AmtB-NH_4_^+^ binding sites but does not influence the conformational flip of the transporter. Within the electro-binding model, many variants can be considered depending on how strongly the membrane potential acts on the periplasmic and cytoplasmic AmtB-NH_4_^+^ binding sites. Here, we focus on the following three representative cases:

- Internal-only *φ*-dependent case, EB-Int-*φ* (electro cytoplasmic binding): *K*_*amtbnhint*_= *φ* · *K*_*EB* -*Int* -*φ*_ and *K*_*amtbnhext*_= *K*_*EB* -*Int* -*φ*_
- Symmetric *φ*-dependent case, EB-Sym-*φ* (electro mixed binding): *K* = *φ*^−0.5^ · *K*_*amtbnhext EB*-*Sym*-*φ*_ and *K*_*amtbnhint*_ = *φ* ^0.5^ · *K*_*EB*-*Sym*-*φ*_
- External-only *φ*-dependent case, EB-Ext-*φ* (electro periplasmic binding): *K*_*amtbnhint*_ = *φ*^−1^*K*_*EB* -*Ext* -*φ*_ and *K*_*amtbnhext*_ = *K*_*EB*-*Ext* -*φ*_

*K*_*EB-Int-φ*_, *K*_*EB-Sym-φ*_, and *K*_*EB-Ext-φ*_ are the case-specific values of *K*_*amtbnhext*_ and *K*_*amtbnhint*_ at *φ* = 1 (i.e., zero membrane potential). For a fair comparison among models, the values of *K*_*EB-Int-φ*_, *K*_*EB-Sym-φ*_, and *K*_*EB-Ext-φ*_ are chosen different such that *K*_*amtbnhext*_ = 5 μM ^3,6^,29 in each case under the reference simulation condition (Δ*Ψ* = - 150 mV, *T* = 310 K, and thus *φ* = 275). Thus, under the reference condition, *K*_*amtbnhext*_ and *K*_*amtbnhint*_ = *φ* · *K*_*amtbnhext*_ are identical across the three electro-binding models. However, when the models deviate from the reference condition, i.e., *φ* ≠ 275, the three models yield different values for *K*_*amtbnhext*_ and *K*_*amtbnhint*_, reflecting their distinct dependencies on *φ*. For complete description of the electro-binding model variants, see Section 3 of Supplementary Notes.

For the electro-flipping hypotheses, we assume

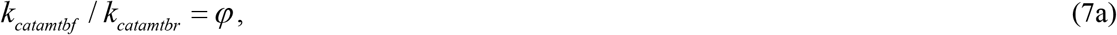

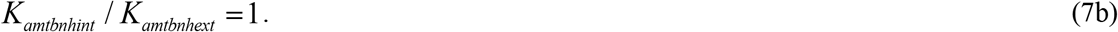

This corresponds to a scenario in which the transmembrane electric potential affects the conformational flip of the transporter but does not influence AmtB-NH_4_^+^ binding. Within the electro-flipping model, many variants can be considered depending on how strongly the membrane potential acts on the inward and outward conformational flips associated with NH_4_^+^ transport. Here, we focus on the following three representative cases:

- Reverse-only *φ*-dependent case, EF-Rev-*φ* (electro outward flipping): *k*_*catamtbf*_ = *k*_*EF* -*Rev* -*φ*_ and *k* = *φ* ^−1^*· k* _*catamtbr EF* -*Rev*-*φ*_
- Symmetric *φ*-dependent case, EF-Sym-*φ* (electro bidirectional flipping): *k*_*catamtbf*_ = *φ* ^0.5^*· k* _*EF* -*Sym*-*φ*_ and *k*_*catamtbr*_ = *φ* ^−0.5^*· k* _*EF* -*Sym*-*φ*_
- Forward-only *φ*-dependent case, EF-Fwd-*φ* (electro inward flipping): *k*_*catamtbf*_= *φ · k*_*EF* -*Fwd* -*φ*_ and *k*_*catamtbr*_= *k*_*EF* -*Fwd* -*φ*_

*k*_*EF-Rev-φ*_, *k*_*EF-Sym-φ*_, and *k*_*EF-Fwd-φ*_ are the case-specific values of *k*_*catamtbf*_ and *k*_*catamtbr*_ at *φ* = 1 (i.e., zero membrane potential). For a fair comparison among models, the same reference value of *k*_*catamtbf*_ is used for all the three cases under the reference simulation condition. After estimating the value of *k*_*catamtbf*_ through parameter estimation (see the next section), *k*_*EF-Rev-φ*_, *k*_*EF-Sym-φ*_, and *k*_*EF-Fwd-φ*_ are backcalculated from this estimated value. As a result, both *k*_*catamtbf*_ and *k*_*catamtbr*_ are nearly identical across the three electro-flipping models under the reference condition. However, outside the reference condition, i.e., *φ* ≠ 275, the three models can produce significantly different values for *k*_*catamtbf*_ and *k*_*catamtbr*_, reflecting their distinct dependencies on *φ*. For complete description of the electro-flipping model variants, see Section 3 of Supplementary Notes.

#### Transmembrane electric potential affects AmtB-NH_4_^+^ dissociation

In the previous study ^14^, we developed a constrained optimization-based parameter estimation technique (see Methods). Briefly, this technique estimates model parameters so that the model can fit experimental data. The model that explains the experimental data with physiologically reasonable parameter values has high model plausibility (*MP*). In the present study, we applied this technique to six models in total, consisting of three electro-binding cases and three electro-flipping cases. We then evaluated which case is the most plausible based on currently available experimental data.

As in the previous study^14^, in the parameter estimation we treated *k*_*catamtbf*_ and *K*_*amtbnhext*_ as Class I and unsearched (US) parameters, respectively. For Class I parameters, measured values are available, whereas US parameters were fixed during parameter estimation. As reference values, we used 4.86 × 10^5^ min^-1^ ^6^ and 5 μM ^3,6^,29 for *k*_*catamtbf*_ and *K*_*amtbnhext*_, respectively, under the reference simulation condition (Δ*Ψ* = -150 mV, *T* = 310 K, and *φ* = 275). For each of the six cases, *k*_*catamtbr*_ and *K*_*amtbnhint*_ were calculated from *φ, k*_*catamtbf*_, and *K*_*amtbnhext*_. We employed the same constraint functions as those used in our previous study ^14^ (i.e., *g*_1_-*g*_58_ in Section 4.2 of Supplementary Information of our previous study^14^). These constraint functions are mainly based on culture experiments with agarose medium ^23^, in a microfluidic chamber ^22^, and with liquid medium ^21^ (See Methods). For each of three electro-binding cases and three electro-flipping cases, we performed five independent parameter estimation runs.

We found parameter sets that satisfied all constraints for all six cases (**Table S6** in Supplementary Tables). In other words, all electro-binding models (EB-Int-*φ*, EB-Sym-*φ*, and EB-Ext-*φ*) and electro-flipping models (EF-Rev-*φ*, EF-Sym-*φ*, and EF-Fwd-*φ*) successfully reproduced experimental data obtained by three independent research groups ^21-23^. In **Figs. S2–S4** (Supplementary Figures), we present simulation results only for the EB-Int-*φ* model, as the other models produced essentially identical outcomes.

The objective function value *f* represents the deviation of model parameters from their experimentally measured values [Eq. (14)]. The model plausibility (*MP*) was calculated from the *f* values [Eq. (15) in Methods]. Among the three electro-binding models, no significant difference in *MP* was observed (Kruskal–Wallis test, p = 0.97). Similarly, no significant difference in *MP* was detected among the three electro-flipping models (Kruskal–Wallis test, p = 0.78). In contrast, a significant difference in *MP* was observed between the electro-binding and the electro-flipping models (Wilcoxon rank-sum test, p = 3 × 10^−6^). When averaging the results of five independent parameter estimation runs across the three variants of the electro-binding model, the *MP* of the electro-binding models was 1.2 × 10^-4^. Likewise, the *MP* of the electro-flipping models was 4.3 × 10^-6^. Thus, the *MP* for the electro-binding models was 28-fold higher than that for the electro-flipping models. Notably, the *MP* of the electro-binding models (*MP* = 1.2 × 10^-4^) was not significantly different from that of our previous model ^14^ (the refined active transporter model; *MP* = 1.1 × 10^-4^; Wilcoxon rank-sum test, p = 0.29).

The *MP* values of the electro-binding models are markedly higher than those of the electro-flipping models. This is because, in order to achieve the ammonium transport rate required for rapid growth, the electro-flipping models require larger deviations of AmtB-related parameter values from their experiment-based reference values.

For the three electro-binding models, using Eqs. (6), the rate equation for the AmtB-mediated NH_4_^+^ transport [Eq. (1)] can be rewritten as

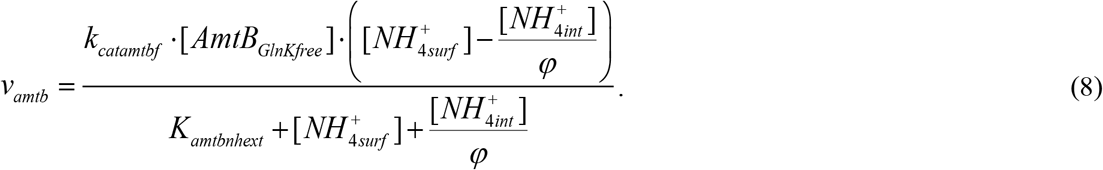

Irrespective of the specific electro-binding variant (EB-Int-*φ*, EB-Sym-*φ*, and EB-Ext-*φ*), the rate equation for the AmtB-mediated NH_4_^+^ transport is given by Eq. (8).

For the three electro-flipping models, using Eqs. (7), Eq. (1) becomes

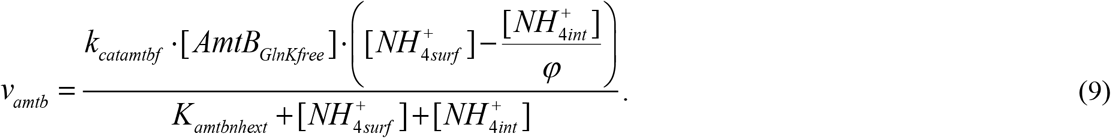

Irrespective of the specific electro-flipping variant (EF-Rev-*φ*, EF-Sym-*φ*, and EF-Fwd-*φ*), the rate equation for the AmtB-mediated NH_4_^+^ transport is given by Eq. (9).

The key difference between the two rate equations [Eqs. (8)-(9)] lies in whether the term for internal NH_4_^+^ appears with a factor of *φ* in the denominator. We examine the requirements for the electro-binding and electro-flipping models to achieve the same AmtB-mediated NH_4_^+^ transport rate:

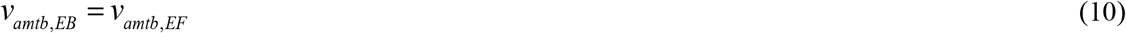

The subscript *EB* denotes the electro-binding models, and the subscript *EF* denotes the electro-flipping models. Here, we consider the experiment executed by Kim et al.^22^, which is the most important among the experimental datasets used for our parameter estimation with respect to AmtB-mediated NH_4_^+^ transport. Specifically, we consider the condition where the extracellular NH_4_^+^ concentration is 4 μM ([*NH*_4_^+^_*surf*_] = 4 μM), i.e., the lowest concentration in these experiments ^22^. Under this condition, the intracellular NH_4_^+^ concentration is 24 μM ([*NH*>_4_^+^_*int*_] = 24 μM) ^22^. Replacing *v* and *v* with Eq. (8) and Eq. (9), respectively, we obtain:

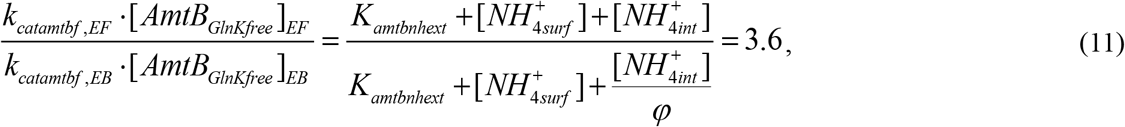

where *K*_*amtbnhext*_ = 5 μM and *φ* = 275 are used. Variables and parameters without either *EB* or *EF* subscripts have identical values in the electro-binding and electro-flipping models. Eq. (11) indicates that, for the electro-flipping model to achieve the same AmtB-mediated NH_4_^+^ transport rate as the electro-binding model, the product *k*_*catamtbf*_ [*AmtB*_*GlnKfree*_] must be 3.6-fold higher than in the electro-binding model. As a consequence, to reproduce the same *v*_*amtb*_ value, the electro-flipping model requires a higher *k*_*catamtbf*_ and/or a higher AmtB concentration than the electro-binding model. This, in turn, leads to an increased concentration of GlnK, because GlnK binds to AmtB in a 1:1 stoichiometric ratio.

The deviations of the estimated parameter values from their respective reference values are shown in **Fig. S5** (Supplementary Figures). The value of *k*_*catamtbf*_ in the electro-flipping model is 2.6-fold higher than that in the electro-binding model. In addition, the estimated AmtB and GlnK levels (for Kim’s experiment) in the electro-flipping model are 1.5-fold and 1.7-fold higher, respectively, than those in the electro-binding model. Indeed, the AmtB level required for the electro-flipping models is comparable to, or even higher than, that of the ATP synthetase—one of the most abundant transporter proteins in *E. coli*, with approximately 10000 ATP synthetase complexes per cell at the specific growth rate of 0.8 h^-1^ ^30^. These larger deviations of the electro-flipping model parameters from their reference values lead to small *MP* values. In contrast, the electro-binding models exhibit smaller deviations of the estimated parameter values from their reference values (blue circles in **Fig. S5**), which leads to higher *MP* values. For further discussion on AmtB-related parameter values, see Section 4 of Supplementary Notes.

In summary, the electro-binding models show a smaller discrepancy with experimental data regarding *k*_*catamtbf*_, AmtB and GlnK levels. This leads to a higher *MP*. Therefore, the transmembrane electric potential likely affects the affinity between AmtB and NH_4_^+^. No significant differences in *MP* were observed among the three electro-binding model variants. Because the membrane potential is constant in the experimental data ^21-23^, all the variants can be fitted to a similar extent, converging to comparable parameter sets through the parameter estimation, which results in similar *MP*. Thus, based on currently available experimental data and *MP*, we cannot determine whether the membrane potential preferentially acts on the periplasmic or cytoplasmic AmtB– NH_4_^+^ binding site. For the analyses that follow, we assume *φ* = 275 unless otherwise explicitly stated. We use the EB-Int-*φ* model as a representative model because the EB-Sym-*φ* and EB-Ext-*φ* models behave indistinguishably from the EB-Int-*φ* model under this condition.

### The importance of the coordinated regulation of GS and AmtB Mechanism of GS and AmtB coordination

The coordinated regulation of GS and AmtB, previously described by Kim et al. ^22^ and us ^14^, is part of our model: When ambient NH_4_^+^ is abundant, passive NH_3_ diffusion supports cell growth. Both AmtB and GlnK are then typically repressed. If expressed, GlnK binds to AmtB and blocks its NH_4_^+^ transport activity (**Fig. 3a**). When NH_4_^+^ is scarce, AmtB and GlnK are highly expressed. The uridylylation of GlnK and its binding to 2-oxoglutarate prevent GlnK from binding to AmtB (**Fig. 3b**). This allows AmtB to transport NH_4_^+^ (or NH_3_ + H^+^) into the cell, raising intracellular NH_4_^+^ levels, possibly over extracellular NH_4_^+^ levels due to the impact of the electric potential across the membrane. However, as the NH_3_ concentration becomes higher in the cytoplasm, much of the transported NH_4_^+^ diffuses back out as NH_3_, creating a futile cycle. This process dissipates the proton motive force because active repeated uptake of the ammonium cation requires one energized proton every cycle, which consumes both constituents of the proton-motive force. In the present study, we assume conditions under which the carbon source is abundant, such that the energetic cost of futile ammonium cycling does not influence growth. Under these conditions, the membrane potential and intracellular pH are treated as constants.

**Fig. 3.**
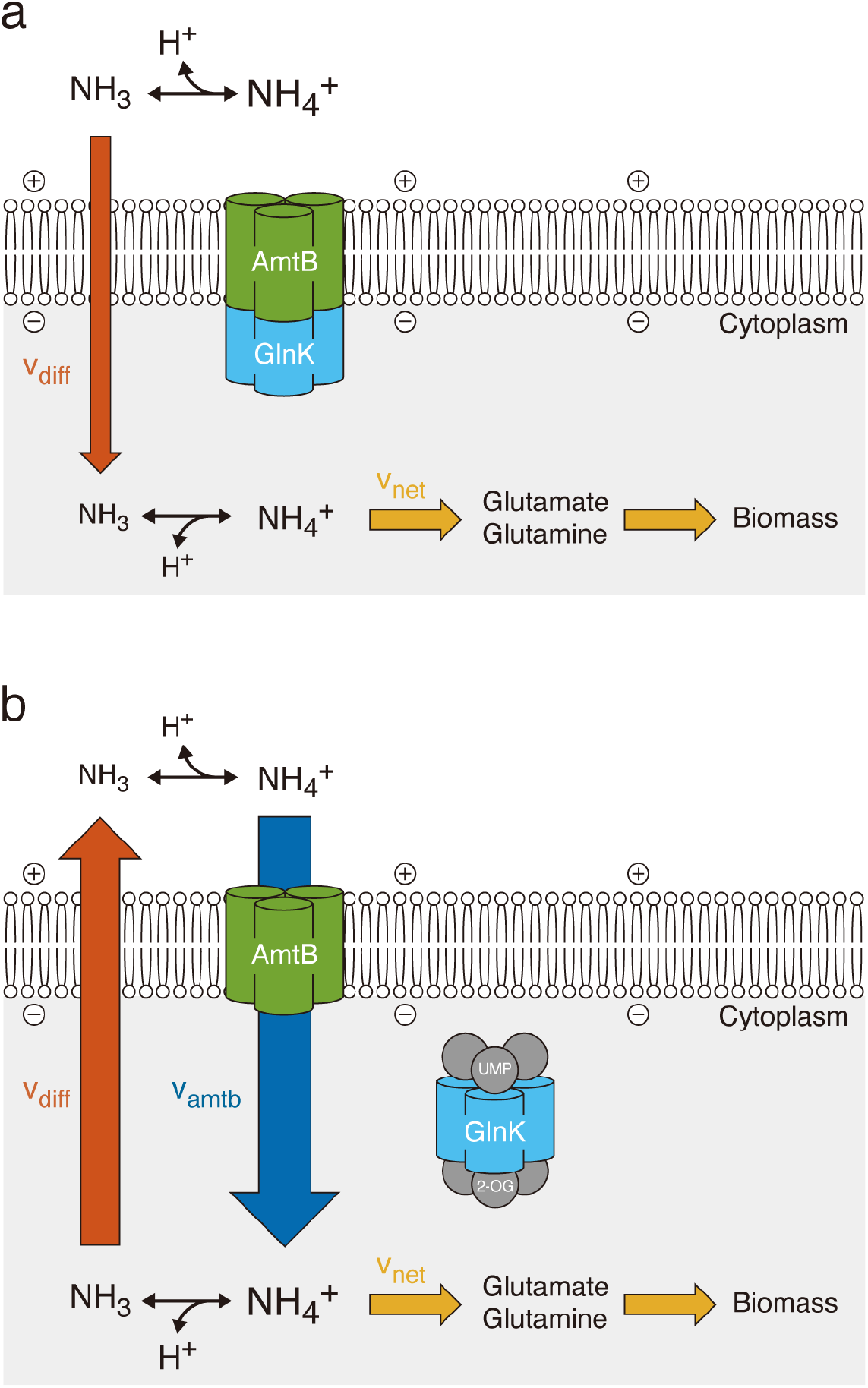
The two modes of ammonium/ammonia transport and the regulation by GlnK. (a) When the extracellular NH_4_^+^ is high, the AmtB expression is low and GlnK binds to AmtB thereby blocking the AmtB-mediated NH_4_^+^ transport. The passive diffusion of NH_3_ alone can then support cell growth. (b) When the extracellular NH_4_^+^ is low, the AmtB expression is high. The uridylylation of GlnK and its binding to 2-oxoglutarate prevent GlnK from binding to and inactivating AmtB. AmtB then transports NH_4_^+^ down its electrochemical gradient, hence towards an increased intracellular NH_4_^+^ concentration, supporting the cell’s growth. The intracellular NH_4_^+^ is de-protonated and produces NH_3_ and H^+^. Since the intracellular NH_3_ concentration thereby becomes higher than the extracellular one, NH_3_ diffuses out of the cell, which is called ‘back diffusion’. The diffused NH_3_ is re-protonated. The NH_4_^+^ transport, NH_4_^+^ de-protonation, NH_3_ back diffusion, and NH_3_ re-protonation constitute a futile cycle which is called futile because it has no effect other than dissipating the proton motive force. Figure adapted and modified from Maeda et al., npj Syst Biol Appl, 2019, published under a Creative Commons Attribution 4.0 International License (http://creativecommons.org/licenses/by/4.0/).

#### Simulation of the coordinated regulation

How does *E. coli* manage the potential futile cycling surrounding nitrogen uptake? To investigate this, we performed simulations at various extracellular NH_x_ (NH_4_^+^ + NH_3_) concentrations. The simulation results are shown in **Fig. 4**. As we are modeling conditions of ammonium limitations, all NH_x_ taken up by the cell minus the NH_x_ flowing back out will finally end up in new biomass. Since the composition of the nitrogen component of biomass (enzymes and nucleic acids) is relatively constant, the rate of glutamate and glutamine incorporation into the biomass should be equal to the specific growth rate multiplied by fixed stoichiometries (see Methods). Consequently, the net NH_x_ import (*v*_*net*_) is directly proportional to the specific growth rate, a net import of 44 mM/min (yellow dotted line in **Fig. 4f**) being required to sustain the specific growth rate (μ) of 0.8 h^-1^ (**Fig. 4c**).

**Fig. 4.**
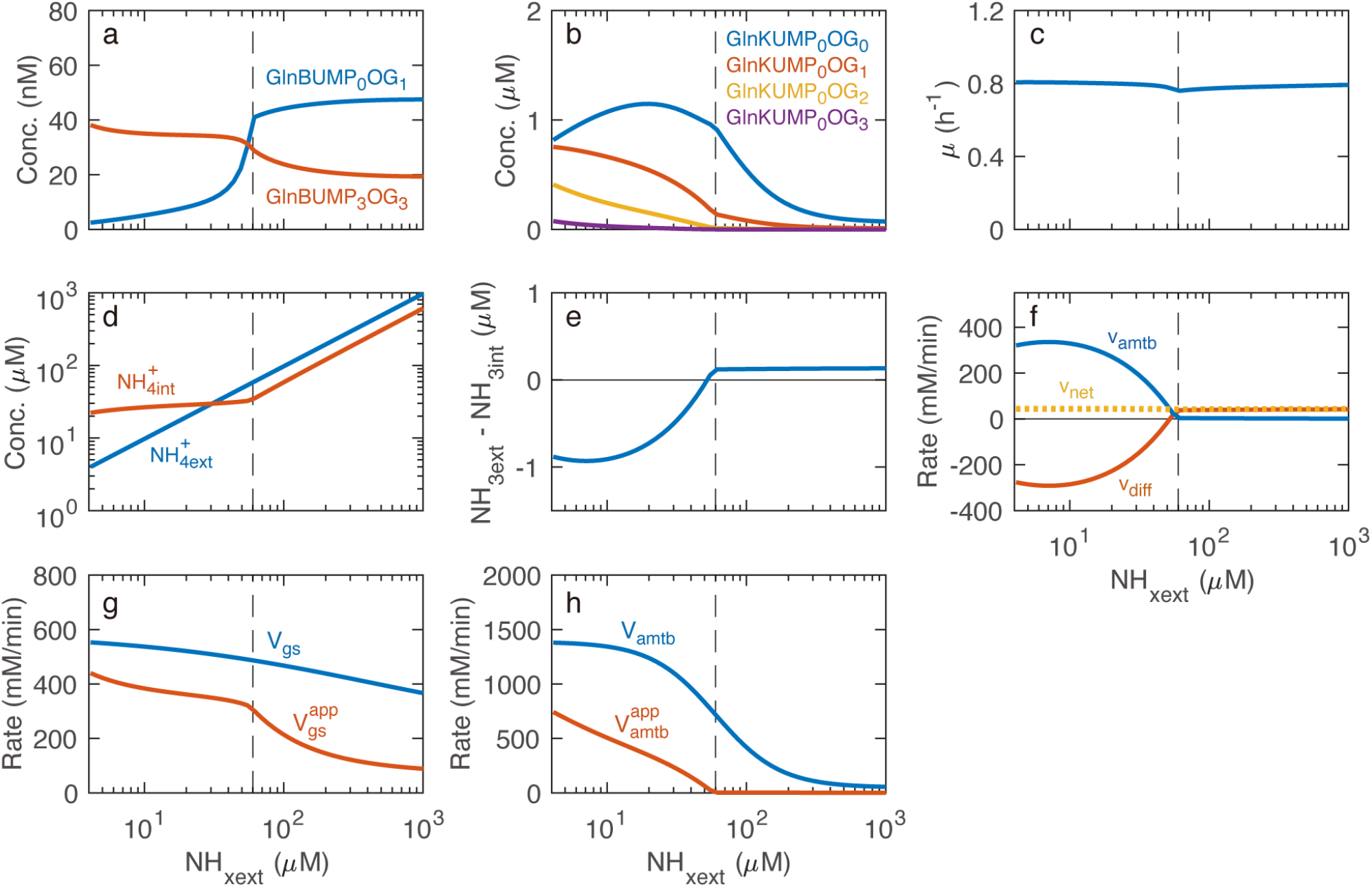
Predicted steady-state behaviors of the wild-type. (a, b) Important GlnB and GlnK species for the coordinated regulation of GS and AmtB. (c) Specific growth rate. (d) Extracellular and intracellular NH_4_^+^ concentrations. (e) Concentration difference of NH_3_, i.e. driving force of unmediated passive diffusion of NH_3_. (f) Ammonium assimilation fluxes. *v*_*amtb*_, *v*_*diff*_, and *v*_*net*_ indicate the AmtB-mediated NH_4_^+^ transport, the unmediated NH_3_ diffusion via the cytoplasmic membrane, and the net flux of nitrogen (*v*_*net*_ = *v*_*amtb*_ + *v*_*diff*_), respectively. Positive and negative values indicate inward and outward fluxes, respectively. (g) The true (*V*_*gs*_) and apparent (*V*_*gs*_^*app*^) maximum rates of the GS reaction. *V*_*gs*_ is proportional to [*GS*_*total*_] which is based on the measured promoter activity. *V*_*gs*_^*app*^ is determined based on *V*_*gs*_ and the adenylylation state of GS. (h) The true (*V*_*amtb*_) and apparent (*V*_*amtb*_^*app*^) maximum rates of AmtB-mediated NH_4_^+^ transport. *V*_*amtb*_ is proportional to [*AmtB*_*total*_] which is based on the measured promoter activity. *V*_*amtb*_^*app*^ is determined based on [*AmtB*^*GlnKfree*^], the active form of AmtB. For details on *V*^*gs*^, *V*_*gs*_^*app*^, *V*_*amtb*_, and *V*_*amtb*_^*app*^, see Methods.

At high extracellular NH_x_ concentrations (above ∼60 μM, i.e., to the right of the vertical dashed line in **Figs. 4a-h**), unmediated NH_3_ inward diffusion was sufficient to maintain a high specific growth rate of 0.8 h^−1^ (**Fig. 4c**). The high concentration of GlnBUMP_0_OG_1_ at the high extracellular NH_x_ concentrations (**Fig. 4a**) implies an increased rate of NRII-GlnBUMP_0_OG_1_-catalysed dephosphorylation of NRI-P and hence repression of AmtB (not shown), GlnK (**Fig. 4b**), and GS (not shown) expression. Moreover, most GlnK remained unuridylylated and unbound to 2-oxoglutarate (**Fig. 4b**). GlnKUMP_0_OG_0_ is bound to the existing AmtB, blocking AmtB-mediated NH_4_^+^ transport. As a result, both the apparent *V*_*max*_ (*V*_*amtb*_^*app*^) and the actual rate (*v*_*amtb*_) of AmtB-mediated NH_4_^+^ transport were effectively zero (**Figs. 4h** and **4f**, respectively). At these extracellular NH_x_ concentrations above ∼60 μM, no futile cycling occurred, because influx through the AmtB (*v*_*amtb*_) was absent (**Fig. 4f**). Although a decrease in extracellular NH_x_ strongly and proportionally reduced intracellular NH_4_^+^ levels (**Fig. 4d**), this did not lower the ammonium assimilation rate: It was compensated for by an increase in the apparent *V*_*max*_ of GS (*V*_*gs*_ ^*app*^) with decreasing extracellular NH_x_ level (**Fig. 4g**), which was achieved somewhat through enhanced GS expression and even more through deadenylylation of already present GS. Adenylylation of GS is catalyzed by ATase and regulated through UTase and GlnB. When the intracellular NH_4_^+^ level decreased, GlnB became uridylylated and bound to 2-oxoglutarate (**Fig. 4a**), leading to increased deadenylylation of GS by ATase.

When extracellular NH_x_ dropped below ∼60 μM, the activation of GS with further reduction in NH_x_ became less intensive (**Fig. 4g**). The model *E. coli* began to express AmtB with GlnK dissociated from it (**Fig. 4h**, both the *V*_*amtb*_ and the *V*_*amtb*_^*app*^ increase with further decreasing *NH*_*xext*_). Furthermore, as extracellular NH_x_ decreased, intracellular 2-oxoglutarate levels increased, promoting the binding of 2-oxoglutarate to GlnK (**Fig. 4b**, red line increasing more than the blue line especially at very low *NH*_*xext*_). This prevented GlnK from blocking AmtB-mediated NH_4_^+^ import. The consequent increase in *V*_*amtb*_^*app*^ (**Fig. 4h**) enabled AmtB-mediated NH_4_^+^ import: *v*_*amtb*_ increased with decreasing extracellular NH_x_ concentrations (**Fig. 4f**). In this range of extracellular NH_x_ concentrations (60 dropping to 4 μM), the intracellular NH_4_^+^ concentration decreased only slightly and kept around 20–30 μM for extracellular NH_4_^+^ concentrations decreasing a further 15-fold (**Fig. 4d**). Since the intracellular pH was assumed to be constant, this meant that the intracellular NH_3_ concentration did not decrease much either, causing a stronger outward NH_3_ gradient with decreasing extracellular NH_x_ concentrations (**Fig. 4e**, blue line dropping below zero). Consequently, more NH_3_ back diffusion occurred (**Fig. 4f**, red line dropping below zero) simultaneously with increased AmtB uptake flux (blue line in **Fig. 4f**), constituting futile cycling.

Overall, our *E. coli* model maintained its growth rate (and hence the net nitrogen uptake rate, *v*_*net*_) within 5 % of μ = 0.8 h^-1^ across a wide NH_x_ range (4–1000 μM) (**Fig. 4c**). As extracellular NH_x_ was decreased and intracellular NH_x_ followed suit, the cell first activated GS while continuing to rely on passive NH_3_ diffusion for nitrogen uptake. This activation compensated for the reduced availability of its substrate, intracellular NH_4_^+^, and kept AmtB inactive, thereby preventing futile cycling. Only when intracellular NH_4_^+^ dropped below ∼60% of the K_m_ of GS (0.1 mM), AmtB was activated. At extracellular NH_x_ concentrations below ∼50 µM, nitrogen was then imported as NH_4_^+^ via AmtB, driven by the transmembrane electric potential. This prevented further declines in intracellular NH_4_^+^ (red line in **Fig. 4d**) despite decreasing extracellular NH_x_ levels, but at the cost of increased futile cycling (**Fig. 4f**; the *v*_*amtb*_ becoming much higher than the *v*_*net*_). GS is activated at higher NH_x_ concentrations than AmtB because GlnB and GlnK respond to metabolic conditions in different ways: GlnB undergoes uridylylation at higher external NH_x_ concentrations than GlnK. At NH_x_ concentrations above the micromolar range, GlnK remains largely unuridylylated but tunes AmtB-mediated NH_4_^+^ transport through its binding to 2-oxoglutarate. Model parameters related to GlnB and GlnK were estimated in the parameter estimation process. For the model to consistently explain the experimental datasets from three independent groups ^21-23^, it was essential that GlnB undergoes uridylylation at higher external NH_x_ concentrations than GlnK. This differential sensitivity is therefore not an arbitrary assumption, but a necessary condition imposed by the combined experimental data. Notably, this behavior emerged as an interesting and non-trivial prediction of the model itself.

To confirm the importance of coordinated regulation between GS and AmtB, we simulated virtual mutants in which the uridylylation states and 2-oxoglutarate binding affinities of GlnB and GlnK were altered (**Fig. S6** in Supplementary Figures). The simulations showed that abnormalities in the uridylylation or 2-oxoglutarate binding states of GlnB and GlnK either impaired the maintenance of rapid growth at low external NH_x_ concentrations or caused excessive ammonium futile cycling. These results highlight that coordinated regulation of GS and AmtB achieves maximal growth while minimizing free-energy loss. For details on the virtual mutants, see Section 5 of Supplementary Notes.

#### AmtB-mediated NH_4_^+^ transport is more free energy-intensive than the GS reaction

In order to quantify the energy cost of futile cycling, we defined *r*_*amtb*_ as the ratio of the AmtB-mediated NH_4_^+^ transport rate to the ammonium assimilation rate [Eq. (24) in Methods]. *r*_*amtb*_ thereby represents the number of times NH_4_^+^ is transported via AmtB before being assimilated. **Fig. 5a** shows how *r*_*amtb*_ depends on the extracellular NH_x_ level. The futile cycling occurred when the extracellular NH_x_ concentration was lower than 50 μM and reached a maximum at 7 μM of almost 8, meaning that out of every 8 NH_4_^+^ molecules transported by AmtB, only 1 was assimilated.

**Fig. 5.**
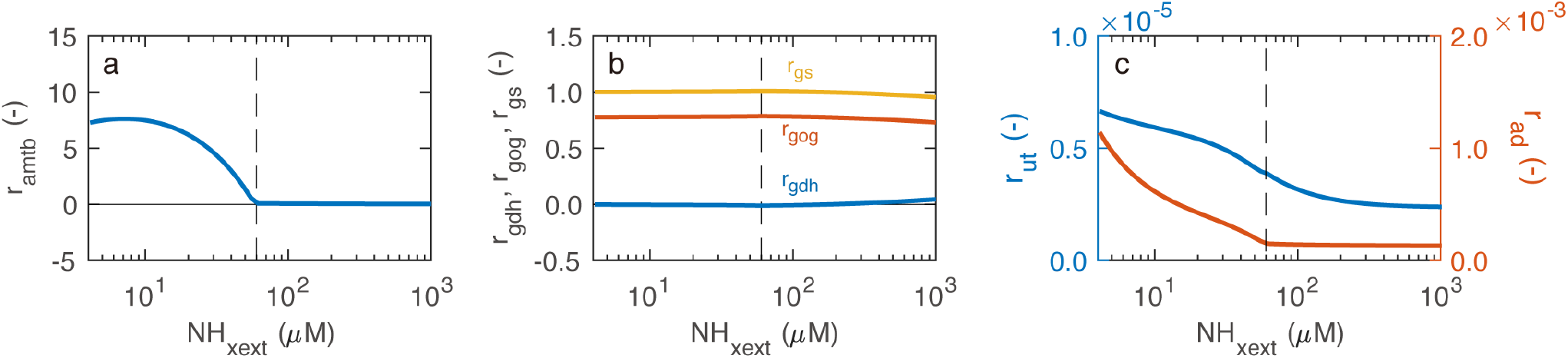
Predicted ratios of the reaction rates to the ammonium assimilation rate. (a) The ratio of the AmtB-mediated NH_4_^+^ transport rate to the ammonium assimilation rate (*r*_*amtb*_). (b) The ratios of GDH, GOGAT, and GS reaction rates to the ammonium assimilation rate (*r*_*gdh*_, *r*_*gog*_, and *r*_*gs*_, respectively). (c) The ratios of uridylyl and adenylyl transfer reaction rates to the ammonium assimilation rate (*r*_*ut*_ and *r*_*ad*_, respectively). For details on *r*_*amtb*_, *r*_*gdh*_, *r*_*gog*_, *r*_*gs*_, *r*_*ut*_, and *r*_*ad*_, see Methods.

The ammonium transport and assimilation network is home to yet another futile cycle (**Fig. 1**): the joint activity of GOGAT and GS leads to the synthesis of glutamate from 2-oxoglutarate and NH_4_^+^ at the cost of ATP hydrolysis. The GDH reaction catalyzes the same conversion without requiring ATP hydrolysis. Theoretically, a metabolic futile cycle could occur, in which glutamate is synthesized from 2-oxoglutarate via the GS/GOGAT cycle and then converted back to 2-oxoglutarate through the reverse GDH reaction. We define *r*_*gs*_ as the flux through the GS reaction divided by *v*_*net*_, representing how many times NH_4_^+^ is incorporated into glutamate to form glutamine before being assimilated into biomass [Eq. (27) in Methods]. Similarly, we define *r*_*gdh*_ and *r*_*gog*_ for the GDH and GOGAT reactions for comparison [Eqs. (25)-(26) in Methods]. **Fig. 5b** shows that *r*_*gs*_ remains close to 1 across a wide range of extracellular NH_x_ concentrations (4 µM to 1000 μM), whereas *r*_*gdh*_ is nearly zero. This indicates that nearly all ammonium assimilation occurs via GS rather than GDH, consistent with previous findings that *E. coli* predominantly employs the GS/GOGAT pathway for glutamate synthesis ^23^. The result also indicates metabolic futile cycling did not occur.

The signal transduction in this system also comes with futile cycling. In a cycle of uridylylation and deuridylylation of either GlnK or GlnB, a UTP is hydrolyzed to UMP and pyrophoshate, ultimately costing two ATP equivalents (**Fig. 1**). We define *r*_*ut*_ as the flux through the uridylyl transfer/uridylyl removing cycle divided by *v*_*net*_ [Eq. (28) in Methods]. Similarly, in a cycle of adenylylation and deadenylylation of GS, an ATP is hydrolyzed to ADP and phosphate (**Fig. 1**). We define *r*_*ad*_ as the flux through the adenylylation/deadenylylation cycle divided by *v*_*net*_ [Eq. (29) in Methods]. **Fig. 5c** shows that *r*_*ut*_ and *r*_*ad*_ are more than three orders of magnitude lower than *r*_*gs*_, suggesting that the energy costs associated with the signal transduction futile cycles are negligible.

The maximum value of *r*_*amtb*_ is 8 (**Fig. 5a**). Assuming that NH_3_ is symported with one proton (H^+^) and considering the H^+^/ATP coupling ratio, the resulting proton motive force dissipation would be equivalent to a loss of over 2.6 ATP per NH_4_^+^ molecule assimilated. This energy cost arises from dissipation of the proton motive force rather than from direct ATP hydrolysis. Since *r*_*gs*_ is consistently close to 1 (**Fig. 5b**), this implies that at low NH_x_ concentrations, the energy cost of AmtB-mediated NH_4_^+^ transport and the associated futile cycling exceeds that of the GS reaction itself. The classic view of ammonium assimilation considered GS as the most energy-intensive step, primarily because it directly consumes ATP. Accordingly, *E. coli* was thought to utilize GDH (K_m_ = 1.0 mM) in nitrogen-rich environments and GS (K_m_ = 0.1 mM) under nitrogen-limited conditions. However, this view has since been refuted ^23^. As our **Fig. 5b** confirms, *E. coli* ‘s ammonium assimilation is always centered around GS rather than GDH, at least under the conditions that we examined. Our analysis further shows that, under low NH_x_ conditions, the dominant energy cost of ammonium assimilation does not stem from biosynthetic ATP consumption by GS, but rather from proton motive force dissipation caused by repeated AmtB-mediated transport. Although proton motive force-driven transport is a common feature of nutrient uptake, our results quantitatively demonstrate that in this system its energy cost can surpass that of the canonical ATP-consuming assimilation reactions.

#### Control of ammonium assimilation is distributed

To investigate whether ammonium assimilation is governed by a single rate-limiting step or if control is distributed across multiple processes, we calculated control coefficients. **Fig. 6** presents these control coefficients as a heatmap, and the corresponding numerical values are summarized in **Table S7** in Supplementary Tables.

**Fig. 6.**
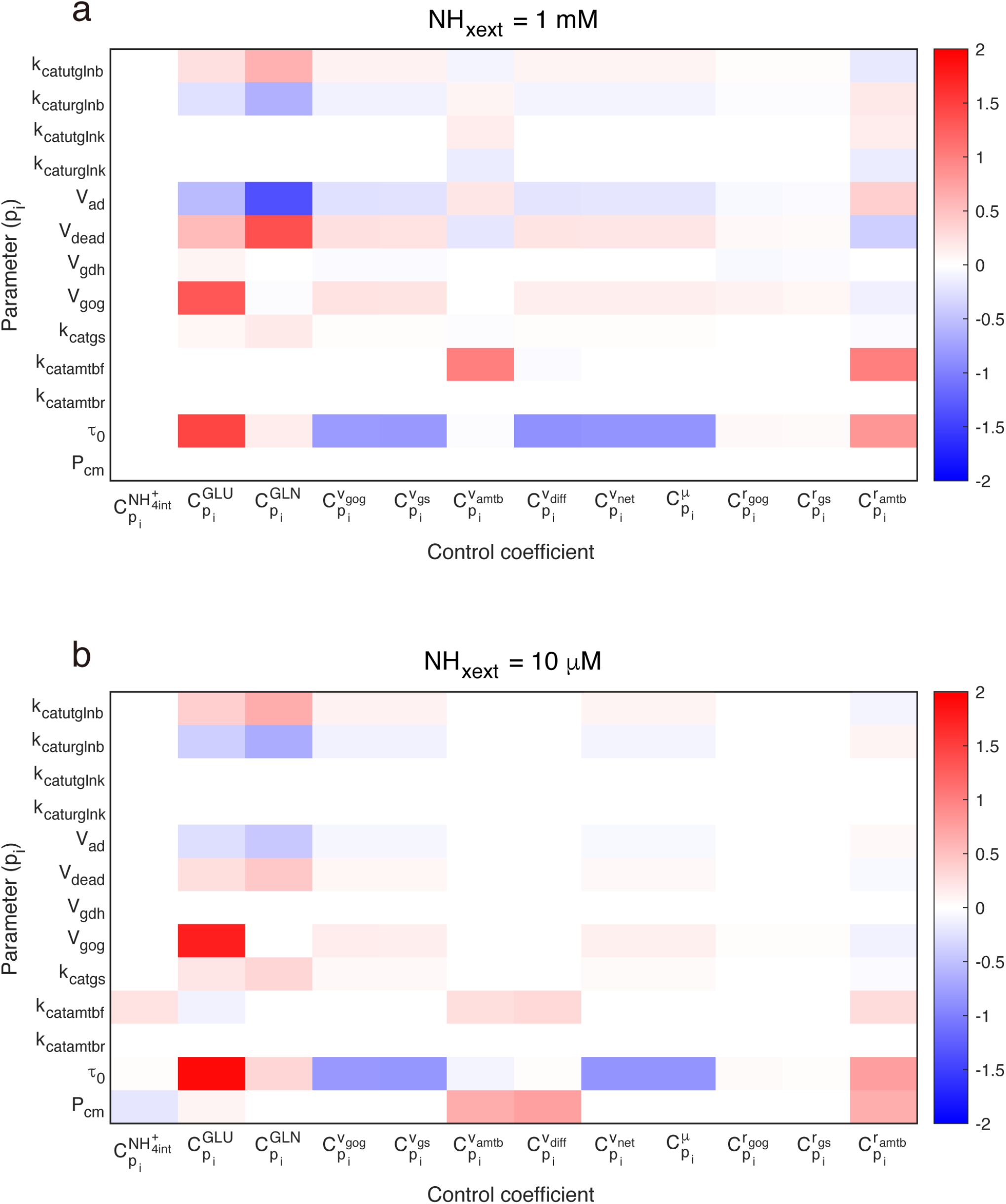
Control coefficients. Control coefficients 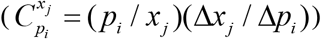 were calculated at steady state. (a) At 1 mM extracellular NH_x_. (b) At 10 μM extracellular NH_x_. The coefficients were obtained by slightly perturbing each parameter (Δ*p*_*i*_/*p*_*i*_ = 10^-6^). Numerical values are provided in **Table S7**.

The summation laws ^31^ were met as shown in **Table S7**, i.e., 1 for fluxes and specific growth rate, and 0 for concentrations and flux ratios. This validates the calculation. We found that the ammonium assimilation (*v*_*net*_) lacks a single rate-limiting step and instead exhibits distributed control. The control is also hierarchical ^32^: the summation laws are met separately at the level of metabolism (*V*_*gdh*_, *V*_*gog*_, *k*_*catgs*_, *k*_*catamtbf*_, *k*_*catamtbr*_, *τ*_0_, and *P*_*cm*_ were included in the sum as they should be ^31^), at the level of GlnB uridylylation (*k*_*catutglnb*_ and *k*_*caturglnb*_), at the level of GlnK uridylylation (*k*_*catutglnk*_ and *k*_*caturglnk*_) as well as at the level of GS adenylylation (*V*_*ad*_ and *V*_*dead*_); signal transduction at steady state involves components with positive and negative controls of equal magnitude ^33^.

The control mechanisms differ significantly depending on the external NH_x_ levels, highlighting dynamic adjustments in the ammonium transport and assimilation network in response to ammonium availability. The absolute values of control coefficients of *V*_*ad*_ and *V*_*dead*_ with respect to glutamate (*GLU*) and glutamine (*GLN*) concentrations were larger under higher NH_x_ concentrations (**Fig. 6a**), indicating that the regulation of GS activity by ATase was more important under ammonium-rich conditions. At *NH*_*xext*_ = 1 mM (**Fig. 6a**), all the control coefficients with respect to intracellular NH_4_^+^ concentration were close to zero. In contrast, at *NH*_*xext*_ = 10 μM (**Fig. 6b**), the absolute values of control coefficients of *k*_*catamtbf*_ and *P*_*cm*_ with respect to intracellular NH_4_^+^ were moderately high (0.23 and -0.20, respectively), indicating that AmtB-mediated NH_4_^+^ transport and unmediated NH_3_ outward passive diffusion had more impact on the intracellular NH_4_^+^ level under the ammonium-limited conditions. The control coefficients of *k*_*catutglnb*_, *V*_*dead*_, and *V*_*gog*_ with respect to *r*_*amtb*_ are negative, suggesting that strengthening the GS/GOGAT cycle reduces the relative magnitude of ammonium-transport futile cycling. For further discussion of the control coefficients, see Section 6 of Supplementary Notes.

### Effect of environmental and physiological parameters Extracellular pH

The external pH significantly influences NH_x_ transport and futile cycling due to its impact on the ratio of NH_4_^+^ to NH_3_. Simulations revealed that at NH_x_ concentrations below 10 µM the growth rate of the ΔAmtB strain was highly sensitive to external pH, making growth impossible at pH 6.9 (**Fig. 7c**). Note that following Kim et al.^22^, we deleted the entire *glnK-amtB* operon (i.e. the code for both AmtB and its regulator GlnK) for what we call the ΔAmtB mutant throughout this study. At lower pH (e.g., 6.9), extracellular NH_3_ concentrations become low (for the same external NH_x_ concentration), limiting unmediated NH_3_ inward diffusion and reducing growth unless NH_x_ concentrations are sufficiently high. For the wild-type strain, which expresses AmtB, growth is robust across a wide range of external pH values (**Fig. 7c;** the solid lines for the various pHs almost coincide). However, at lower pH levels, futile cycling becomes more prominent in the wild-type cells (**Fig. 7d-f;** futile cycling is absent for the ΔAmtB mutant). We define *r*_*amtb,max*_ as the maximum values of *r*_*amtb*_ when the external NH_x_ was changed. We also define *NH*_*xext,futile*_ as the highest external NH_x_ concentration at which futile cycling occurs. *r*_*amtb,max*_ and *NH*_*xext,futile*_ increase as the external pH decreases (**Fig. 7gh**), meaning that futile cycling becomes more significant at lower external pH values.

**Fig. 7.**
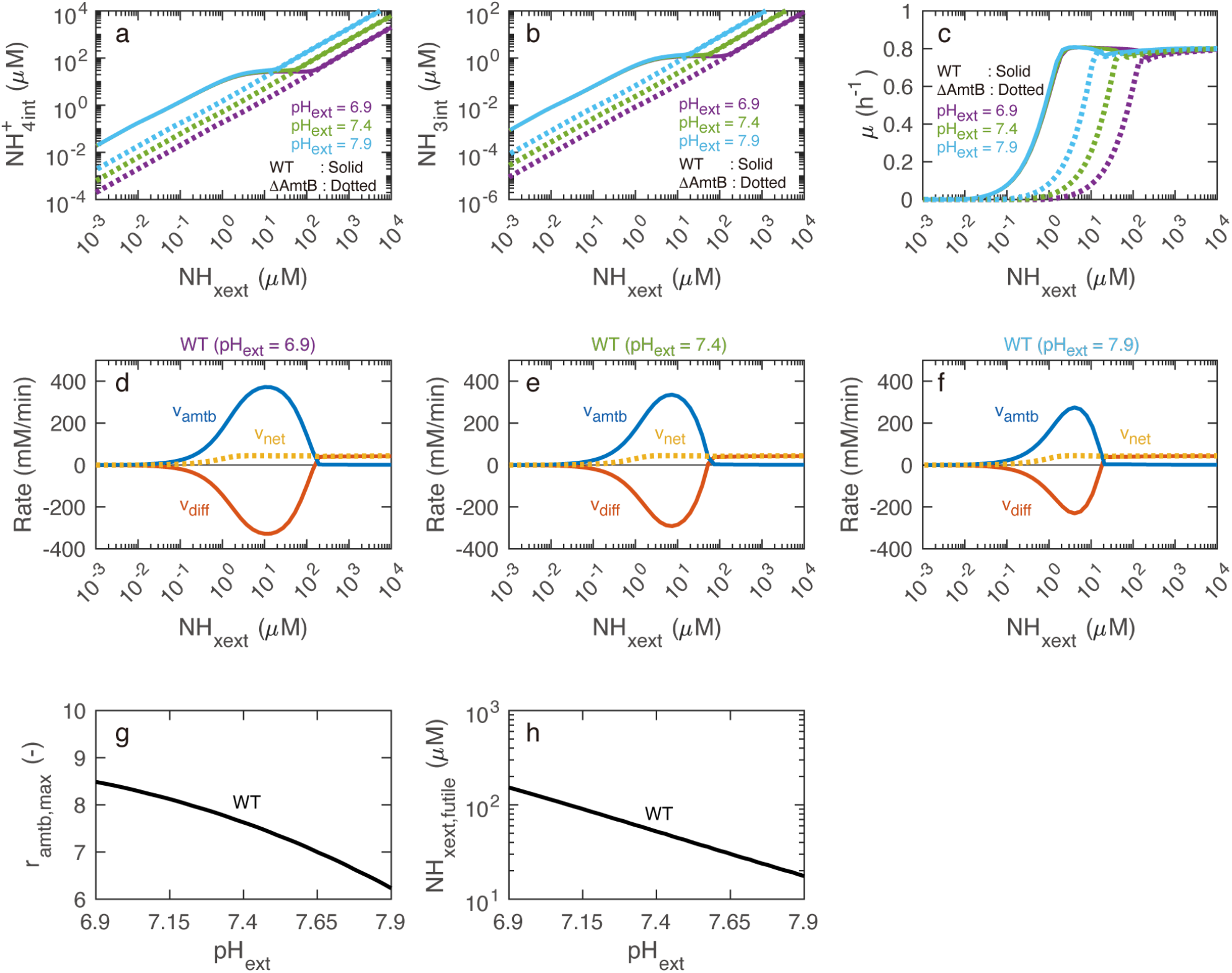
Effects of changes in extracellular pH on steady-state behaviors of the wild-type and ΔAmtB strains. (a) Intracellular NH_4_^+^ concentration. (b) Intracellular NH_3_ concentration. (c) Specific growth rate. (d–f) AmtB-mediated NH_4_^+^ transport (*v*_*amtb*_), unmediated NH_3_ diffusion (*v*_*diff*_), and net flux (*v*_*net*_) at external pH of 6.9, 7.4, and 7.9. Positive flux values indicate the movement of NH_x_ from the extracellular environment into the cytoplasm. (g) The maximum values of *r*_*amtb*_ when the external NH_x_ was changed (*r*_*amtb,max*_). (h) The highest external NH_x_ concentration at which futile cycling occurs (*NH*_*xext,futile*_). The intercellular pH was kept at its normal value (7.6).

#### Intracellular pH

The typical cytoplasmic pH of *E. coli* is 7.6 ^34^, close to the typical pH of 7.4 for media, used for example in Kim et al.^22^. However, in principle, by lowering its intracellular pH ^35^, *E. coli* can reduce internal NH_3_ concentrations, promoting NH_3_ inflow down its concentration gradient—a process known as acid trapping. Simulations showed that lowering intracellular pH by up to one unit enabled ΔAmtB cells to sustain growth at lower external NH_x_ concentrations (**Fig. 8c**). However, further lowering the internal pH provided little additional benefit, as the NH_3_ concentration gradient became less responsive to changes, i.e., the internal NH_3_ concentration was near zero (**Fig. 8b**). Therefore, the effect of acid trapping on NH_x_ import is limited compared with the AmtB-mediated NH_4_^+^ transport. In wild-type cells with AmtB, reducing intracellular pH further improves growth at low NH_x_ concentrations (**Fig. 8c**) while decreasing futile cycling (**Fig. 8d-f**). Both *r*_*amtb,max*_ and *NH*_*xext,futile*_ decreased as the internal pH was lowered (**Fig. 8gh**).

**Fig. 8.**
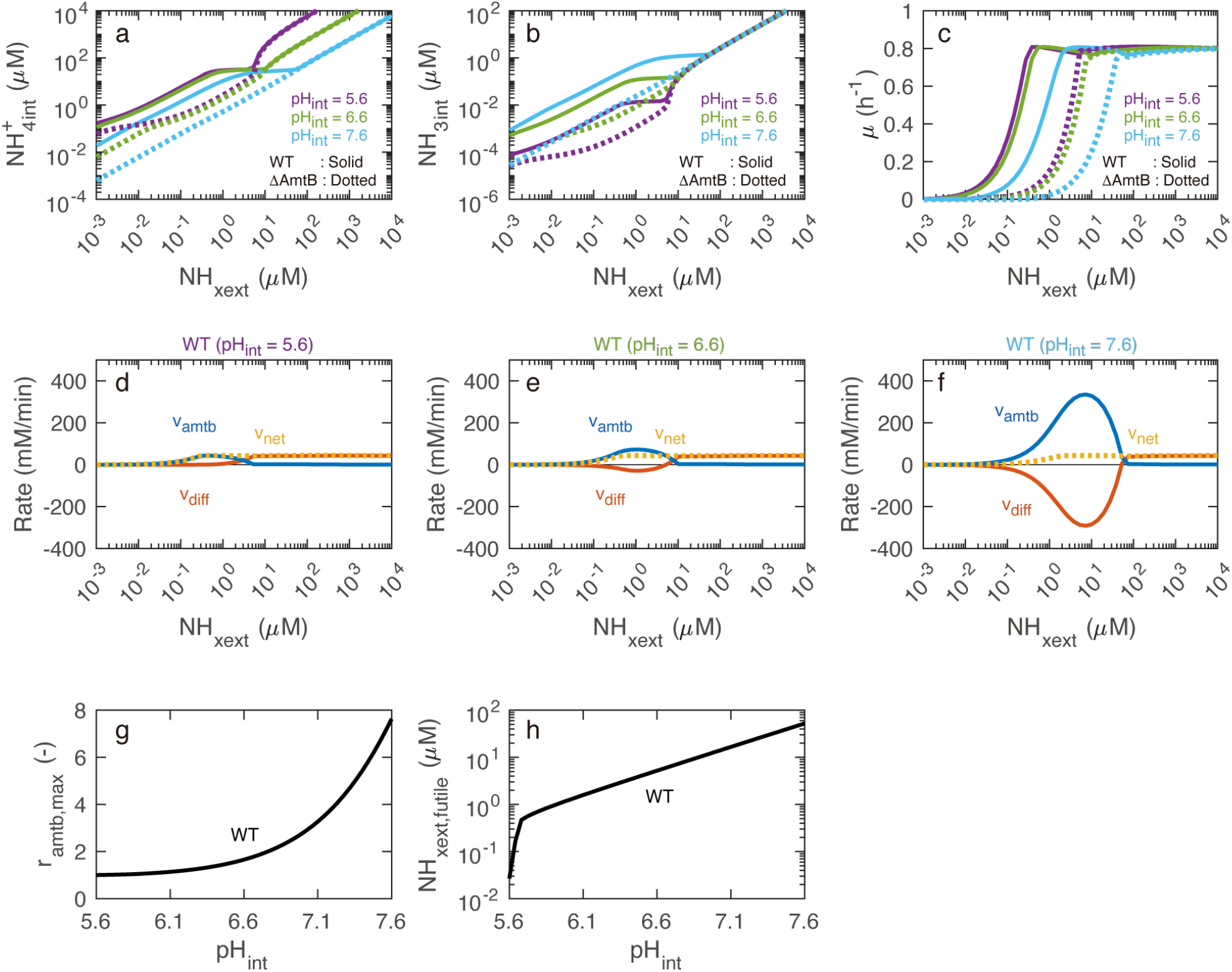
Effects of changes in intracellular pH on steady-state behaviors of the wild-type and ΔAmtB strains. (a) Intracellular NH_4_^+^ concentration. (b) Intracellular NH_3_ concentration. (c) Specific growth rate. (d–f) AmtB-mediated NH_4_^+^ transport (*v*_*amtb*_), unmediated NH_3_ diffusion (*v*_*diff*_), and net flux (*v*_*net*_) at internal pH values of 5.6, 6.6, and 7.6. Positive flux values indicate the movement of NH_x_ from the extracellular environment into the cytoplasm. (g) The maximum values of *r*_*amtb*_ when the external NH_x_ was changed (*r*_*amtb,max*_). (h) The highest external NH_x_ concentration at which futile cycling occurs (*NH*_*xext,futile*_). The extracellular pH was kept at its normal value (7.4).

#### Transmembrane electric potential

In the simulations presented so far, we have employed the EB-Int-*φ* model, i.e., one of the electro-binding models. This is because, under the reference condition (Δ*Ψ* = -150 mV, *T* = 310 K, and thus *φ* = 275), the three electro-binding model variants (EB-Int-*φ*, EB-Sym-*φ*, and EB-Ext-*φ*) exhibit identical behavior. In this section, we investigate how model behavior changes in response to variations in the transmembrane electric potential (Δ*Ψ*). Therefore, we consider not only the EB-Int-*φ* model but also the EB-Sym-*φ* and EB-Ext-*φ* models.

The transmembrane electric potential (Δ*Ψ*) ranges from -85 to -220 mV, depending on growth conditions ^36-38^. We performed simulations at Δ*Ψ* values of -85, -150, and -220 mV, corresponding to NH_4_^+^ accumulation factors (*φ*) of 24, 275, and 3774, respectively. We found that with respect to the intracellular NH_4_^+^ and NH_3_ concentrations, specific growth rate, and futile cycling the EB-Int-*φ* model is insensitive to changes in the membrane potential (**Fig. 9a-c** and **Fig 9jk**). In contrast, *r*_*amtb,max*_ of the EB-Ext-*φ* model was highly sensitive to changes in the membrane potential (**Fig. 9g-i**). This result implies that robust cellular growth can be maintained when the membrane potential interferes with the binding between AmtB and NH_4_^+^ on the cytoplasmic side. For reference, among the electro-flipping models (the less plausible models), the EF-Rev-*φ* model showed the greatest insensitivity to changes in the membrane potential (**Fig. S7** in Supplementary Figures).

**Fig. 9.**
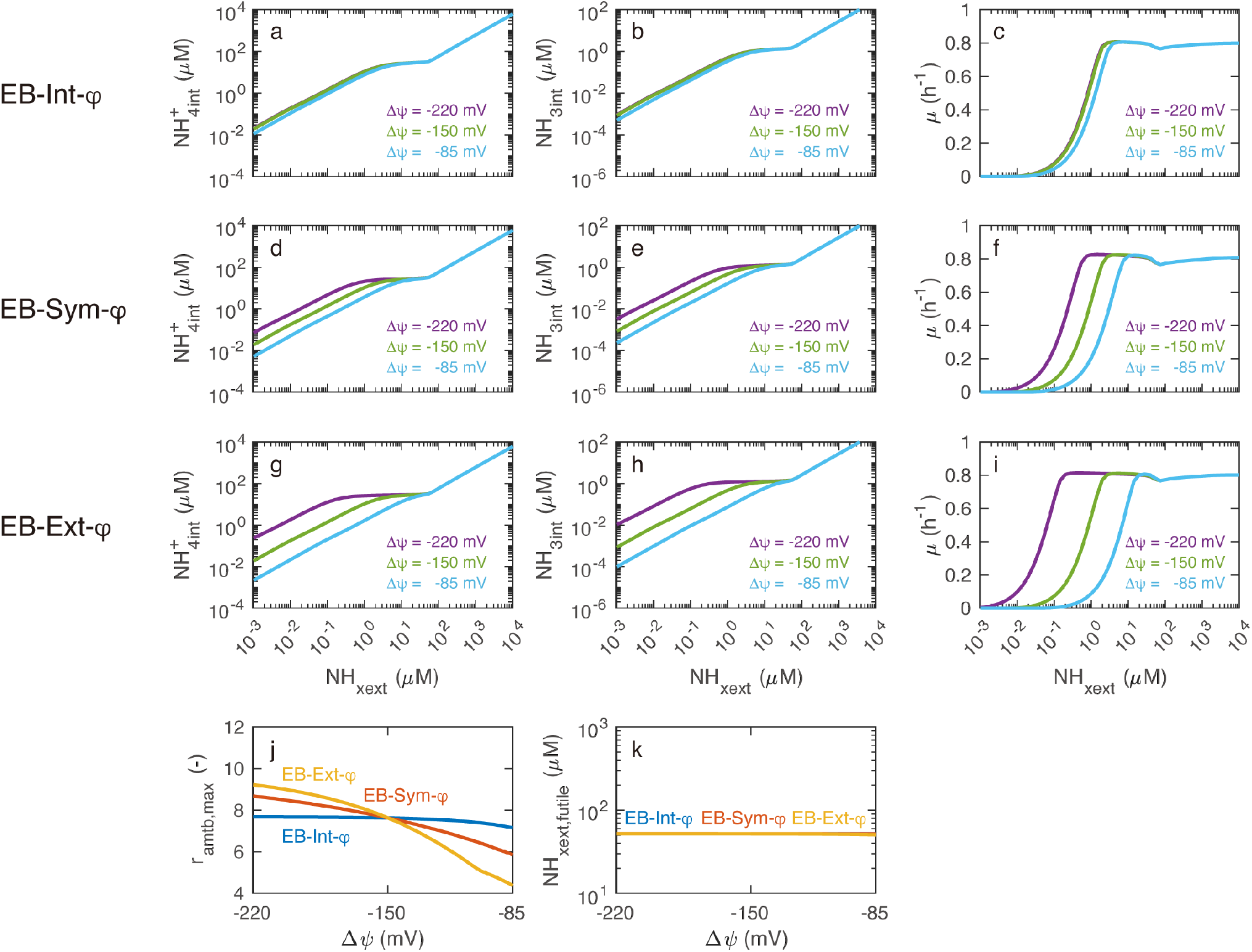
Effects of changes in transmembrane electric potential on steady-state behaviors of the electro-binding model variants. Transmembrane electric potential was assumed to affect the ammonium affinity of the periplasmic and/or cytoplasmic AmtB-NH_4_^+^ binding sites but does not influence the conformational flip of the transporter. EB-Int-*φρ*: Transmembrane electric potential affects the cytoplasmic AmtB-NH_4_^+^ binding. EB-Sym-*φρ*: Transmembrane electric potential affects both the periplasmic and cytoplasmic AmtB-NH_4_^+^ binding. EB-Ext-*φρ*: Transmembrane electric potential affects the periplasmic AmtB-NH_4_^+^ binding. (a, d, g) Intracellular NH_4_^+^ concentration. (b, e, h) Intracellular NH_3_ concentration. (c, f, i) Specific growth rate. (j) The maximum values of *r*_*amtb*_ when the external NH_x_ was changed (*r*_*amtb,max*_). (k) The highest external NH_x_ concentration at which futile cycling occurs (*NH*_*xext,futile*_). In panel (k), the blue, red, and yellow lines coincide.

## Discussion

Ammonium is a primary nitrogen source for bacteria, making its transport and assimilation critical to understanding bacterial physiology. This transport does not only raise intracellular NH_4_^+^ concentrations but may also lead to free-energy-consuming futile cycling, where NH_3_ diffuses out and NH_4_^+^ is re-imported. Despite its energetic cost, this futile cycling is not truly futile: it is a side effect of useful active transport. The active transport enables *E. coli* to grow under ammonium-limited conditions by maintaining the intracellular NH_4_^+^ levels necessary for metabolism and growth, whilst the passive NH_3_ transport is a biophysical feature stemming from the high membrane permeability for gases such as ammonia. The present study shows that this interpretation is fully realistic in view of the most relevant experimental data that exist in three quantitative studies ^21-23^.

This study used a detailed, previously calibrated and substantially validated kinetic model ^14^ to explore the mechanism by which the transmembrane electric potential energizes ammonium transport by AmtB, the regulation of GS and AmtB, and the response of the transport plus assimilation system to environmental and physiological changes. First, we compared the electro-binding and electro-flipping models. The model comparison showed that the electro-binding models are more consistent and have a much higher plausibility vis-à-vis the experimental data. Notably, the electro-binding models reproduce the ammonium transport rates required for growth while remaining closer to experimentally supported parameter values and requiring lower levels of cellular components such as AmtB and GlnK. This resource efficiency may be biologically advantageous. We also found that all the three electro-binding models were equally plausible. This is because the existing experimental data were obtained at the physiological value of the membrane potential. Among three variants of the electro-binding model, the EB-Int-*φ* model was the most robust to variations in transmembrane electric potential however (**Fig. 9**). This suggests that transmembrane electric potential drives the ejection of ammonium from the ‘cytoplasmic’ ammonium binding site in AmtB to the cytoplasm. Experiments performed under controlled variations of membrane potential will be required to directly test this hypothesis.

Using the most plausible and robust model, i.e., the EB-Int-*φ* model, we demonstrated that as external NH_x_ decreases, *E. coli* first upregulates GS and then activates AmtB to maintain nitrogen assimilation (**Fig. 4**): which of GS or AmtB plays the dominant role in ammonium assimilation depends on the external NH_x_ level, as also indicated by the control coefficients (**Fig. 6** and **Table S7**). While AmtB enables robust growth across a wide range of NH_x_ concentrations and pH levels, it also causes futile cycling—particularly under acidic conditions (**Fig. 7**). Some of these phenomena have been reported previously, for example in Fig. 6 of our earlier study ^14^. However, in the present study, we performed a more detailed analysis demonstrating that GS and AmtB regulation is coordinated through both UTase activity and direct binding of 2-oxoglutarate. Impairment of UTase activity or 2-oxoglutarate binding results in reduced growth or elevated futile cycling (**Fig. S6** in Supplementary Figures). By comparing the magnitude of futile cycling in ammonium transport, metabolism, and signal transduction, we found that the free-energy cost of futile cycling in ammonium transport, driven by AmtB-mediated NH_4_^+^ uptake and NH_3_ back diffusion, exceeds that of the GS reaction itself (**Fig. 5**). Finally, our simulations reveal that AmtB enables robust cellular growth under varying ammonium concentrations and pH levels, albeit at the cost of futile cycling (**Fig. 7 and 8**).

Where structural studies often inform kinetic models, the present study illustrates an interesting reversal of this logic: kinetic and thermodynamic constraints restrict the class of transport mechanisms that can be consistent with structural observations. Structural methodologies cannot directly resolve electric potentials or proton dynamics, neither can they reliably distinguish ammonium from ammonia, nor protonated from unprotonated histidine residues. This limitation is critical, because whether nitrogen is transported as ammonia or as ammonium has profound consequences for the very function of AmtB at low external ammonium concentrations^12^, as well as for futile cycling and Gibbs energy dissipation. With respect to possible histidine binding sites for ammonium/ammonia, this has some extra relevance: In the AmtB structure (PDB ID: 1U7G), 3 ‘ammonia binding sites’ are mentioned that are close to what has been called the hydrophobic pore between H168 and F31. According to Bizior et al. ^9^, two of these are histidines (i.e., H168 and H318). These are called ammonia, i.e., NH_3_, binding sites, but they may just as well be called ammonium binding sites: the same bound state would arise from binding of ammonium to unprotonated histidine as from binding of ammonia to protonated histidine. For H168 and H318 it is likely that in the absence of ammonia they are unprotonated *in vivo*: due to the high electric field existing between their locations and the cytosol their pKa should be shifted downward by 2.5 units. And, in the absence of continued supply of ammonium from the periplasm such as in the crystals, the sites may be occupied by water ^10^ rather than ammonium.

Williamson et al., proposed a two-lane mechanism for AmtB-mediated ammonium transport ^16^ illustrated by their Fig. 7, as well as Fig. 2 of Bizior et al. ^9^. In these graphical representations of this transport mechanism, we note a potential shortcoming: ammonia (NH_3_) is depicted as passing through the Phe gate, while the proton is shown to move separately around that gate via a water wire. Subsequent passage to H318 is also depicted as two independent lanes. The spatial separation of ammonia and proton fluxes in the diagram of the two-lane mechanism suggests a lack of coupling between the two or at least refrains from addressing the mechanism of this coupling. We emphasize that this proposed spatial separation between protons and ammonia is not supported by structural observation of the actual positions of the protons and ammonia being transported. It is a consequence of an uncertain mechanistic interpretation and highlights the need to integrate kinetic and thermodynamic constraints when inferring transport mechanisms from structural models. We will now attempt this integration.

For the function of AmtB, which is intracellular accumulation of ammonium, the coupling between ammonia and proton transport is essential ^12^. Therefore the mechanism of coupling should preferably be addressed in proposed mechanisms of transport. In free-energy transduction, strict coupling of two processes has at least two inextricable features ^39^: (i) the one flux cannot proceed without the other and (ii) each process can be driven by the thermodynamic force of the other process notwithstanding its own perhaps counteracting force. As protons nor proton flux can be observed by the structural methods, the question remains how the membrane potential (which is the dominant component of the protonmotive force) could energize ammonium transport through AmtB. Building on both structural information and kinetic–thermodynamic constraints, we here propose such a strictly coupled transport mechanism in which ammonia and proton translocation are causally and energetically linked, even if they are not always chemically bound together as in NH_4_^+^ (**Fig. 10**). In this view, the species progressing through the hydrophobic pore is protonated ammonia, i.e., ammonium in which two of its protons are shared between ammonia and two weak hydrogen-bonding residues lining that pore, notably H168, W212, or H318 ^9-10^; we are proposing a dual complex between the ammonium and two proton accepting residues lining the pore. Such shared binding delocalizes charge widely (over two aromatic rings and the ammonia core between them) and transiently allows this complex to occupy the low-dielectric environment of the hydrophobic pore into which it is coerced by the very strong electric field at the gate (F215 + F107) and in the hydrophobic pore (H168 - F31).

**Fig. 10.**
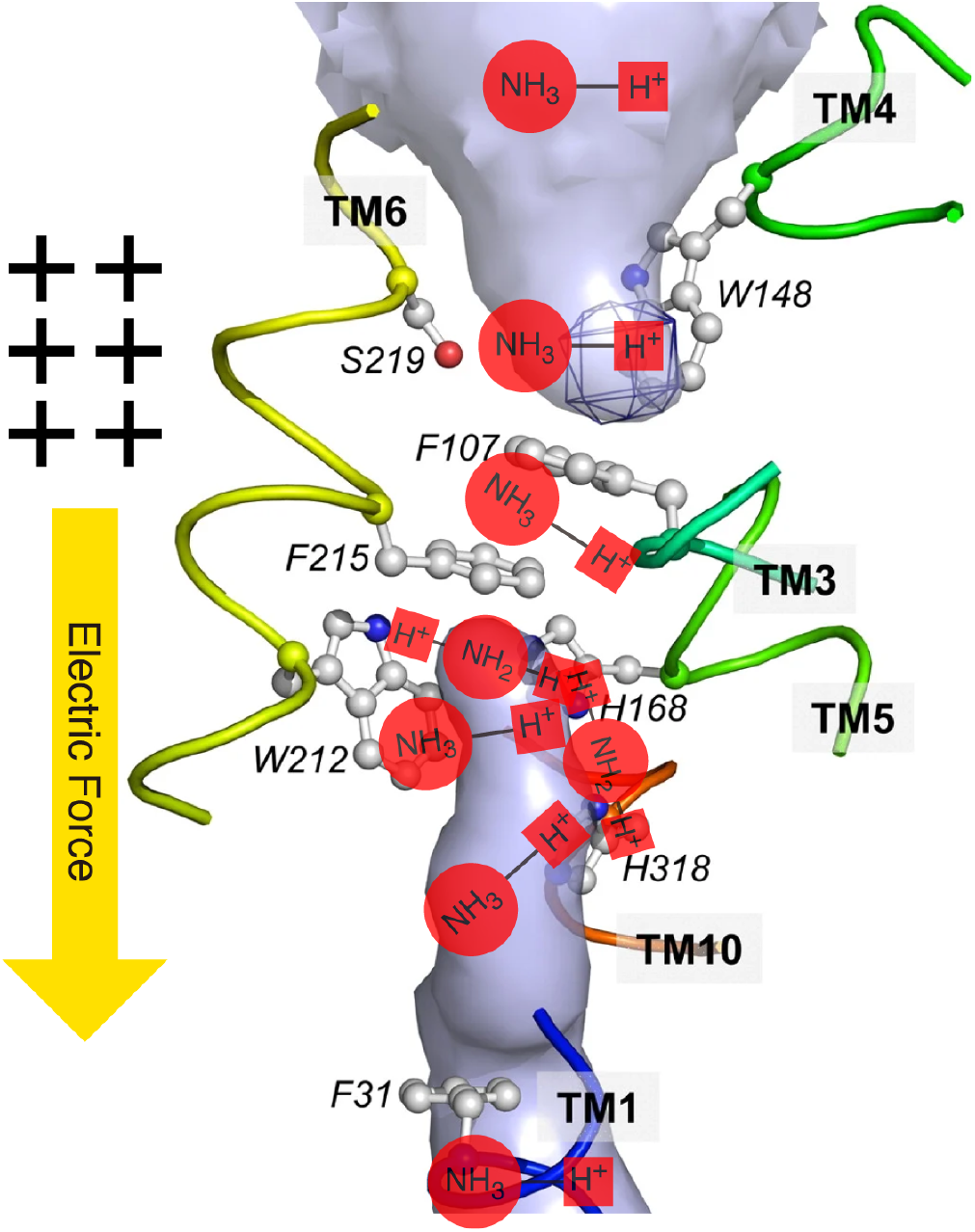
Proposed strictly coupled mechanism for ammonia and proton transport by AmtB. NH_4_^+^ equilibrates with the periplasm and binds at the S1 periplasmic binding site (W148 and S219). Upon stochastic opening of the Phe gate (F107 and F215), NH_4_^+^ flies into the hydrophobic pore, where it encounters more of the strong electric field. Within the pore, the ammonium forms a partially protonated dual complex with W212 and H168 or H318, or with H168 and H318, or with H318 and a water at the bottom of the pore. Transport proceeds as a stepwise handover of this weakly bound dual complex along the His–Trp–His motif, driven by the membrane potential. The state of the ammonium is typically the two-armed dual complex (A-H-NH_2_-H-B)^+^, where A and B represent a neutral Trp or His lining the hydrophobic pore or water at the bottom of or inside the hydrophobic pore. The single positive charge is delocalized over A and B (i.e., the Histidine 5-ring(s) and the Tryptophan 9-double-ring), the two hydrogens, and the ammonia core (-NH2-). The dual complex displaces itself by swinging one of its arms in protonated form to the next His or water, like a monkey swinging through the jungle. This movement only happens when driven by the strong electric field in the hydrophobic pore and when the proton charge is delocalized between Trp, His, and ammonia core. H^+^ and NH_3_ translocation are herewith strictly coupled, preventing independent H^+^ slippage or NH_3_ diffusion. Full re-protonation occurs when the dual complex releases both its arms and becomes ammonium ion near the cytoplasmic side prior, upon which it is released into the cytosol, although also in the water the ammonium may continue to swing in similar ways. Figure adapted and modified with kind permission from Arnaud Javelle, Domenico Lupo, Pierre Ripoche, Tim Fulford, Mike Merrick, Fritz K Winkler, “Substrate binding, deprotonation, and selectivity at the periplasmic entrance of the *Escherichia coli* ammonia channel AmtB,” PNAS, 105(13):5040–5045 (2008), DOI:10.1073/pnas.0711742105. Copyright (2008) National Academy of Sciences, U.S.A.

This electric field is likely to be very strong because the space (the periplasmic vestibule) separating the periplasmic binding site of ammonia (S1^10^) near W148 appears to be wide and hydrophilic enough ^10^ to allow periplasmic ions such as K^+^ to move and water dipoles to reorient in response to any electric field, with the effect of virtually annihilating such a field in the vestibule. Consequently, S1 is likely to be at almost the same electric potential as the periplasm. This further makes the electro *periplasmic binding* model (EB-Ext-*φ*) in which the membrane potential influences ammonium binding at the periplasmic site, unlikely.

For the same reasons, there should be no electric field in the vestibule between the cytoplasm and F31 either. However, that cytoplasmic vestibule only runs up to the cytoplasmic side of the so-called ‘hydrophobic pore’, i.e., to F31. As a consequence, virtually the entire transmembrane electric field should be concentrated in the area between the S1 binding site and F31, i.e., across the gate and the hydrophobic pore itself (**Fig. 10**). This combination of structural information with electrostatic theory makes the electro *cytoplasmic binding* model (EB-Int-*φ*) the most likely amongst the electro-binding models, confirming the conclusion we drew above already: the electric potential primarily reduces the affinity of cytoplasmic ammonium for the S2 binding site. The effect of the membrane potential on the affinity of the S2 site for cytoplasmic ammonium thereby provides a rectification mechanism that prevents rebinding of ammonium from the cytoplasm and enforces net inward transport, provided the membrane potential acts on the AmtB, i.e., not in the crystal.

**Fig. 10** depicts the mechanism we propose in the structure defined by Fig.3 in Javelle et al.^40^. In this mechanism transport begins with spatial equilibration of NH_4_^+^ between the bulk periplasm and the periplasmic binding site S1 of AmtB. NH_4_^+^ transiently binds to W148 (and S219) while awaiting stochastic opening of the Phe gate. We propose that the gating residue F107 oscillates between interaction with F215 and interaction with the aromatic ring of W148. When F107 associates with W148, it interferes with NH_4_^+^ binding at this site. This dynamic competition creates a correlation between NH_4_^+^ release from its occupancy at the S1 site (near W148) at the border of the periplasmic vestibule and gate opening, thereby contributing to conduction specificity for NH_4_^+^ over similarly sized ions such as Na^+^ or K^+^ that do not bind to W148. Upon transient opening of the Phe gate —when F107 flips away from F215 (and may even associate transiently with W212)— NH_4_^+^ can move through the gate; it ‘flies’ as if in a mass spectrometer, from S1 (W148) to S2 (H168-W212) and perhaps even further (see below). Here motion is already biased by the transmembrane electric potential focused on the area between F107 and F31 due to the vestibules on either side. After passing the gate as ammonium ion and entering the hydrophobic pore the ammonium ion continues to encounter the strong electric field that drops across this region (see above). Within the pore, charge localization on ammonia as in ammonium is energetically unfavorable because the ammonium is so small. We therefore propose that the ammonium forms a partially protonated, weakly bound dual complex with two pore-lining residues. An unprotonated H168 shares a proton with the ammonia moiety, becoming partially protonated, while a second ammonium proton transiently interacts with the aromatic nitrogen of W212, which has meanwhile moved back to bind to F107, thereby closing the gate and preventing passage of K^+^ and H^+^ ions. In this configuration, the positive charge is delocalized over ammonium, histidine, and tryptophan, allowing the complex to occupy the otherwise low-dielectric environment of the pore. Occasionally the interaction with W212 is released, and the corresponding partially charged ammonium arm with the proton flies deeper into the pore whilst its other arm holds on to H168. The ammonium ion subsequently engages both H168 and H318, again through shared protons, further delocalizing the charge over the two histidines and the ammonia core. Driven by the strong electric field, this charged species advances vectorially toward the cytoplasmic side while remaining transiently tethered, now to H318: this will look like a monkey swinging through the jungle. Near the cytoplasmic exit, the electric force promotes reassociation of the proton arms of the ammonia with water molecules, regenerating the structure of NH_4_^+^ in water, which may correspond to a monkey likewise swinging through the water. The fully protonated ammonium then enters the cytoplasmic vestibule, where the electric field is negligible and NH_4_^+^ equilibrates with the cytoplasm.

Because the successive bindings at the histidine–tryptophan–histidine motif within the pore are loose, the movement of the ammonium through gate and pore can almost be seen as a flight through a mass spectrometer, perhaps explaining the observed isotopic fractionation in favor of ^14^N over ^15^N ^41^, which used to be taken ^10^ to evidence dissociation of ammonium into ammonia plus a proton (the half dissociations of two ammonium protons may provide an alternative explanation).

Although inspired by and in many aspects similar to this earlier proposed mechanism, the strictly coupled mechanism we propose differs fundamentally from the two-lane mechanism proposed by Bizior et al. ^9^ and Williamson et al. ^16^, in which ammonia and proton translocate via spatially separate pathways. Any such structural separation would permit proton slippage and dissipation of the proton-motive force, in contradiction with thermodynamic requirements and would not provide a driving force for inward transport of the ammonia in the ammonia lane. The hydrophobic nature of the pore has been taken as making the movement of ammonium through that pore impossible^10^. That argument foregoes the effect of the very strong electric field (amounting to up to 15 kJ/mol in energy) helping to push the ammonium into the pore whilst the binding to histidines and tryptophan, two at a time, and concomitant extensive delocalization of a single positive charge, should further reduce the energy of ammonium in the pore.

In keeping with the strictly coupled mechanism, the most plausible kinetic model in the present study (i.e., the EB-Int-*φ* model) independently predicts that ammonium binding to the cytoplasmic-facing site of AmtB is disfavored by the membrane potential. This prediction emerged from the kinetic analysis without additional structural assumptions. This shows how kinetic/thermodynamics may add information to structural information and how that is useful when examining possible mechanisms. With all this we note that more experimental work is needed before we can definitively decide in favor of (or against) the transport mechanism we propose here. In particular measurements at various transmembrane electric potentials should be informative, and mutations altering the distances between the presumed water-proton wires and the hydrophobic pore.

There are two sets of mutants that may seem to question the mechanism we propose here^16,42^. One is that of mutation of F107 to A107, which leaves the AmtB active. We reckon that in that mutant the F215 may still flip periodically to W212 thereby periodically opening the gate for ammonium to flow through in the strong electric field. The other is single mutations of H168 or H318 not inactivating the AmtB but merely reducing its specificity for ammonium over K^+^. We reckon that in this mutant the ammonium would be more on a par with the K^+^, explaining the reduced specificity; the NH_3_ monkey could still swing if one of the lianas is missing.

Javelle et al. ^43^ proposed that GS and AmtB are metabolically coupled, based on *in vivo* kinetic analyses showing an apparent increase in GS affinity for ammonium compared with *in vitro* measurements. However, their conclusions were derived from experiments using methylammonium (MA) as a substrate analogue, such that the reported kinetic parameters reflect the apparent K_m_ of GS for MA rather than for NH_4_^+^. In addition, AmtB was at that time interpreted as an NH_3_-conducting channel, an assumption that has since been revised. The GS–AmtB coordination described in the present study is conceptually distinct from the metabolic coupling proposed by Javelle et al. Rather than invoking an intrinsic enhancement of GS substrate affinity through AmtB-mediated ammonium import, our model emphasizes regulatory coordination between GS and AmtB activities without the two proteins touching each other. This coordination enables cells to maintain rapid growth under ammonium-limiting conditions while minimizing energetic costs, by appropriately modulating the activities of GS and AmtB in response to ammonium limitation stress.

Without active ammonium transport, there should be neither ammonium futile cycling, nor the associated free energy loss. Is there any way for *E. coli* to grow rapidly without AmtB under ammonium-limited conditions? We tested the “acid trapping” strategy *in silico* and found that, even with a 2-unit pH decrease, the calculated growth rate of the ΔAmtB mutant did not surpass that of wild-type *E. coli* with AmtB (**Fig. 8c**). Similarly, reducing cell size or increasing NH_3_ permeability showed limited effects on growth (**Figs. S8-S9** in Supplementary Figures and see also Sections 7-8 of Supplementary Notes). Therefore, active ammonia transport (such as transport of NH_4_^+^, or NH_3_ transport coupled to a transported proton) remains essential for maintaining the high intracellular NH_4_^+^ levels required for rapid growth. There is no evidence of transport of NH_x_ directly coupled to ATP hydrolysis.

Our investigation suggests that, in wild-type *E. coli*, lowering the internal pH should be an effective strategy to reduce intracellular NH_3_ levels and ammonium futile cycling (**Fig. 8g**). Even a small decrease of 0.5 pH units from the normal internal pH (7.6) should significantly reduce futile cycling. The feasibility of this approach is supported by a paper ^35^ demonstrating that *E. coli* can acidify its cytoplasm to cope with stress and thereby facilitate the development of antibiotic resistance. This strategy of lowering the internal pH could also apply to other bacteria facing high levels of ammonium futile cycling (see refs ^44-46^ for a general review on intentional bacterial behavior).

The insights gained from *E. coli* may extend beyond this species and help understand ammonium transport and assimilation in other bacteria, as the GS/GOGAT system for ammonium assimilation is widely conserved. The BRENDA database ^47^ (consulted on 7-10-2025) contains only four K_m_-values for ammonium that are lower than the 100 µM of *E. coli*’s GS (EC 6.3.1.2): 15-50 µM for *Anabaena variabilis, Paracoccus denitrificans, Azospirillum brasilense*, and *Trichormus azollae*. In addition, the K_m_ values of *Anabaena* 7120 ^48^ and *Hyphomicrobium* X ^49^ were within this range. The K_m_ values shown in BRENDA for 13 other bacteria are higher than 100 µM ammonium. This suggests that many bacteria using the GS/GOGAT system may face even greater challenges with futile cycling than *E. coli*. For more discussion of the K_m_ of GS, see Section 9 of Supplementary Notes and **Fig. S10** in Supplementary Figures.

Extrapolating our results obtained at external pH 6.9 (**Fig. 7d**), our prediction would be that moderate (pH of 3-5) and extreme (pH of <3) acidophiles with active ammonium transport face notable free-energy loss due to futile cycling. However, it would not be much worse than for *E. coli* at pH 6.9 as the driving force for outward ammonia diffusion is primarily determined by internal NH_3_ levels. Moreover, acidophiles maintain an internal pH of approximately 6.5, implying that to the extent that NH_x_ transport attains equilibrium, their internal NH_3_ concentration should be 10-fold lower than that in *E. coli* ^50^.

Thermophilic bacteria may also experience intensified futile cycling due to higher NH_3_ concentrations and permeability caused by elevated temperatures (see Sections 8 and 10 of Supplementary Notes and **Figs. S9** and **S11** in Supplementary Figures). At 70°C, the pKa of ammonium is as low as 8, increasing NH_3_ levels by 10-fold compared to 35°C, while higher membrane permeability further exacerbates NH_3_ back diffusion. Similarly, bacteria with small cell sizes face greater free-energy loss because of their high surface-area-to-volume ratio, which amplifies NH_3_ back diffusion and futile cycling (see Section 7 of Supplementary Notes and **Fig. S8** in Supplementary Figures). For example, *Prochlorococcus marinus*, a marine cyanobacterium is small (0.1 µm^3^ ^51^) and uses GS/GOGAT as the main N-assimilation pathway (see ref ^52^ and references therein) with a low GS affinity for ammonium (K_m_ = 300 µM ^53^). Its growth rate is quite slow (0.01-0.02 h^-1^ ^54^).

Regulation of cell metabolism involves multiple tiers of the cell’s hierarchy, which may all act at significant though different strengths ^55^. In ‘metabolic regulation’ changes in metabolite levels lead to a new balance of reaction rates. In signal-transduction regulation of metabolism, covalent modification of enzymes steered by signal transduction cascades alters enzyme activities. In gene-expression regulation of metabolism, changes in metabolite levels adjust gene-expression resulting in altered concentrations of enzymes. Modeling all three tiers dynamically and at the same time, tends to reproduce the confusion that one already has in direct experimental observation of the regulation. For gaining understanding it may be better therefore to model the slower tiers non-dynamically, by having them defined by empirical information. This is the approach we took here: we inserted gene expression leading to AmtB, GlnK, and GS levels as fixed functions of the extracellular ammonium concentrations. Thereby our results reveal that already the dynamics at the metabolic and signal transduction levels suffice to produce robustness of growth against changes in extracellular pH and ammonium concentrations.

Our kinetic model enables “perfect experiments” in which individual parameters can be varied independently—something difficult to achieve in wet experiments. In the model, the number of NH_4_^+^ transport events via AmtB before assimilation (*r*_*amtb,max*_) is 8 for normal cells (**Fig. 5a**). Since each transport costs about one-third of an ATP equivalent, futile cycling imposes an additional 2–3 ATP cost per assimilation. This may present cells with a substantial energy burden, especially under carbon-limited conditions where ATP and proton motive force generation are compromised. Our simulations assume excess carbon and thus may underestimate this effect. Moreover, the growth rate was computed based on intracellular glutamate and glutamine concentrations following Yuan et al. ^23^: this does not explicitly account for the energetic consequences of ammonium futile cycling and may therefore overestimate growth under energy-limited conditions. Despite these simplifications, our model captures key aspects of ammonium metabolism, provides mechanistic insight into how cells adapt to environmental changes, and provides a new mechanism of energy coupling in transport. We hope that the *in silico* discoveries made in the present paper, can inspire others and us to the design of new experiments that can then further test, extend, and where necessary improve the model and understanding.

## Methods

### Parameter estimation and model plausibility analysis

Kinetic models contain model parameters such as maximum velocities and Michaelis constants, and they have not been measured in most cases. Even when model parameters have been measured, they are often obtained under different experimental conditions, most commonly *in vitro*. Therefore, to simulate *in vivo* cellular behavior, parameter values must be estimated or fine-tuned. In a previous study ^14^, we developed a constrained optimization-based parameter estimation technique that allows a kinetic model to fit experimentally observed behaviors while maintaining realistic parameter values. Here, we provide a brief overview of this method. For a detailed description, please refer to our previous study ^14^.

The parameter estimation problem can be formulated as a constrained optimization problem:

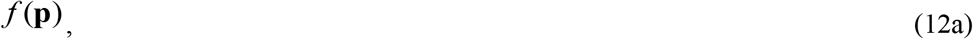

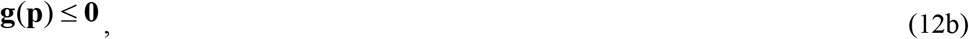

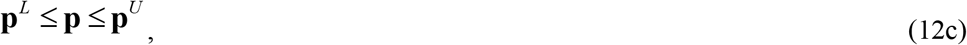

In this framework, **p** = (*p*_1_, *p*_2_, …) denotes the vector of parameters to be estimated, where each *p*_*i*_ represents the *i*th search parameter. **p**^*L*^ and **p**^*U*^ denote the lower and upper bounds of the parameter search space, respectively. The objective function *f* quantifies the deviation of the estimated parameters from their reference values—these reference values correspond to the initial guesses, which may be based on experimental data, educated guesses, or rough guesses. The vector **g** = (*g*_1_, *g*_2_, …) represents the constraint functions that assess how well the model reproduces the training data, i.e., the experimental observations used for calibration. For example, a constraint function that evaluates model fitting to time course data is given by

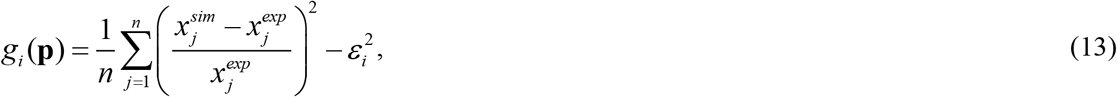

where *x*_*j*_^*sim*^ and *x*_*j*_^*exp*^ are simulated and experimental data points of a model variable, *n* is the number of data points, and *ε*_*i*_ is the allowable error. If the fit is inadequate, the corresponding *g*_*i*_ takes a positive value.

The goal of this constrained optimization problem is to minimize deviations from the reference parameter values while maintaining an acceptable fit to the training data (**g** ≤ **0**). The constraint functions used here are identical to those employed in our previous study (see Section 4.2 of Supplementary Information of our previous study^14^) and were constructed based on experimental datasets from three earlier works ^21-23^ (see the following section). To solve the parameter estimation problem, we employed a genetic algorithm named IS-SR-REX^star^/JGG^14,56-57^, which efficiently explores the parameter space under multiple constraints. Parameter estimation was performed on a Mac Studio (M3 Ultra) using parallel computation to evaluate multiple candidate solutions simultaneously. A single run required 6 hours.

We divided search parameters into three classes (see **Table S4** in Supplementary Tables): Class I, II, and III parameters are those for which measured values (I), educated guesses (II), and rough guesses (III) are available. In this study, the objective function *f* is given by:

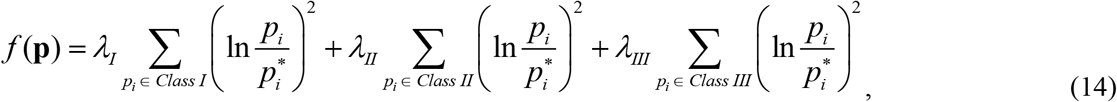

where 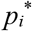 is the reference value of the *i*th parameter, and *λ*_*j*_ (*j* = *I, II, III*) is the class-related penalty weight for a parameter change (*λ*_*I*_ > *λ*_*II*_ > *λ*_*III*_ ≥ 0). We used *λ*_*I*_ = 1.0407, *λ*_*II*_ = 0.1930, and *λ*_*III*_ = 0. These values correspond to assuming that ln(*p*_*i*_/*p*_*i*_^∗^) follows a normal distribution with standard deviations of ln(2), ln(5), and infinity for Class I, II, and III parameters, respectively. This choice reflects the assumption that deviations of Class I and Class II parameters from their reference values are unlikely to exceed twofold and fivefold changes, respectively. In contrast, Class III parameters are allowed to vary freely, because their values have not been experimentally measured and are difficult to guess reliably.

A model is considered valid only if all constraints satisfy (*g*_*i*_ ≤ 0 for all *i*), ensuring consistency with experimentally observed behaviors. For such models, *f* remains almost always positive, allowing model plausibility (*MP*) to be quantified based on deviations from reference values. Since larger deviations from the reference values indicate a less realistic model, *MP* decreases as parameters diverge from their reference values. We can calculate model plausibility (*MP*) from *f*:

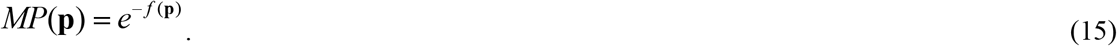

Thus, minimization of *f* is equivalent to maximization of *MP*. When all parameters are equal to their reference values (*p*_*i*_ = *p*_*i*_^∗^), *MP* attains its maximum value of 1. As discussed in Section 3 of the Supplementary Information of our previous work ^14^, *MP* can become substantially small even when only a single parameter deviates markedly from its reference value. Consequently, *MP* generally takes small values in practice. For reference, when one of Class I parameters is 19-fold away from its reference value, and the other parameters are exactly equal to their reference values, then *f*(**p**) = 9.0 and *MP*(**p**) = 1.2 × 10^-4^ (which is close to the *MP* of the electro-binding models of the present study). On the other hand, all Class I parameters deviate twofold from their reference values, and all Class II parameters deviate fivefold from their reference values, we obtain *f*(**p**) = 50 and *MP*(**p**) = 1.9 × 10^-22^. Furthermore, as discussed in Section 14 of the Supplementary Information of our previous work ^14^, the choice of the assumed probability distribution affects the absolute value of model plausibility. However, it does not affect qualitative conclusions, such as which model exhibits higher plausibility.

In summary, our constrained optimization-based parameter estimation technique ensures that the model reproduces experimentally observed behaviors while minimizing the deviation of estimated parameters from their reference values. This approach allows us to construct realistic models that maximize *MP*. When comparing different models, *MP* serves as a quantitative measure of model reliability. For a complete description on *f, MP*, and parameter estimation, please refer to Sections 3, 4, and 14 of Supplementary Information of our previous study ^14^.

### Experimental data for parameter estimation and model plausibility analysis

As in our earlier study ^14^, experimental data used for parameter estimation and model plausibility analysis were primarily obtained from three previous studies ^21-23^. Yuan et al. ^23^ cultivated *E. coli* strains (wild-type, ΔGDH, and ΔGOGAT) on filters placed atop solid agarose media, which enabled rapid and noninvasive sampling of intracellular metabolites. Nitrogen limitation was achieved by lowering the NH_4_^+^ concentration, which reduced but did not stop growth, thereby confirming nitrogen-limited conditions. When these N-limited cultures were transferred to plates containing 10 mM NH_4_^+^ (an N-upshift), the growth rate was restored partially. Metabolite samples were collected at multiple time points for up to 30 minutes after the N-upshift. Kim et al. ^22^ used a microfluidic cultivation system to culture *E. coli* under stable ammonium-limited conditions. From measured growth rates, they estimated intracellular NH_4_^+^ concentrations, as well as rates of ammonium transport via AmtB, non-facilitated ammonia diffusion, and ammonium assimilation. Radchenko et al. ^21^ cultured *E. coli* under N-limited conditions and induced an N-upshift by adding 200 μM NH_4_^+^ to liquid cultures. They monitored the uridylylation state of GlnK and association of GlnK with AmtB both before and periodically after the N-upshift. These three sets of experimental data collectively allowed us to thoroughly validate our model across various conditions of ammonium limitation and availability.

### Important variables

In this section, we explain the important players in determining the physiological behavior of ammonium assimilation. For a complete model description, see also **Tables S1–S4** in Supplementary Tables.

Glutamine Synthetase (GS) is an essential enzyme in ammonium assimilation that uses glutamate, NH_4_^+^, and ATP as substrates to produce glutamine, ADP, and inorganic phosphate (P_i_). GS undergoes adenylylation or deadenylylation, depending on the intracellular concentrations of 2-oxoglutarate and glutamine. Adenylylation reduces GS activity, and the apparent *V*_*max*_ of GS (*V*_*gs*_ ^*app*^) is expressed as:

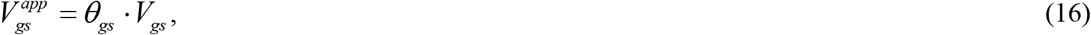

where *V*_*gs*_ is the *V*_*max*_ of the GS reaction when the enzyme is maximally activated in terms of adenylylation, and *θ*_*gs*_ is a factor ranging from 0 to 1, determined by the adenylylation state of GS. The *V*_*max*_ of GS (*V*_*gs*_) is calculated as:

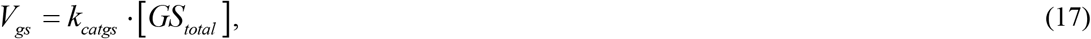

where [*GS*_*total*_] = [*GS*] + [*GSAMP*]. [*GS*] and [*GSAMP*] represent the concentrations of the deadenylylated GS dodecamer and adenylylated GS dodecamer, respectively. [*GS*_*total*_] is the total concentration of GS dodecamer, which is a function of the external NH_4_^+^ concentration. The function is based on promoter activity data ^22^ (see Section 1 of our Supplementary Notes). The rate equations for the adenylylation (*v*_*ad*_) and deadenylylation (*v*_*dead*_) of GS are listed in **Table S1** in Supplementary Tables. As originally proposed in ref ^58^ the function *θ*_*gs*_ reads:

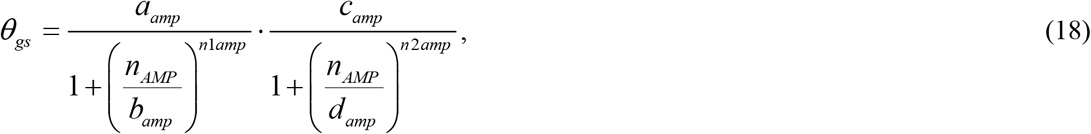

where *a*_*amp*_, *b*_*amp*_, *c*_*amp*_, *d*_*amp*_, *n*_*1amp*_, and *n*_*2amp*_ are model parameters (**Table S4** in Supplementary Tables). *θ*_*gs*_ ranges from 0 to 1 (*a*_*amp*_ *· c*_*amp*_ = 1 when *n*_*AMP*_ = 0). *n*_*AMP*_ is the average number of adenylylated subunits in the GS dodecamer:

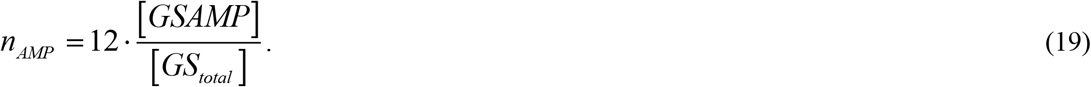

AmtB is a trimeric ammonium transporter, the activity of which is blocked when trimeric GlnK binds to it ^19-21^. The binding of GlnK to AmtB is regulated by UTase depending on the concentrations of glutamine. The apparent *V*_*max*_ of AmtB (*V*_*amtb*_^*app*^) is expressed as:

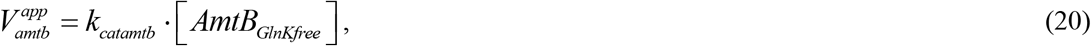

where [*AmtB*_*GlnKfree*_] is the concentration of AmtB that is not bound to GlnK. The maximum AmtB activity (*V*_*amtb*_) is given by:

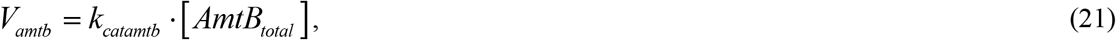

where [*AmtB*_*total*_] = [*GlnKAmtB*] + [*AmtB*_*GlnKfree*_]. [*GlnKAmtB*] is the concentration of AmtB bound to GlnK. [*AmtB*_*total*_] is the total concentration of AmtB. Similarly to [*GS*_*total*_], [*AmtB*_*total*_] is taken to be a function of the external NH_4_^+^ concentration. The actual rate of AmtB-mediated NH_4_^+^ transport (*v*_*amtb*_) is given by Eq. (1). The rate of unmediated NH_3_ diffusion across the cytoplasmic membrane due to the concentration gradient (*v*_*diff*_) is given by

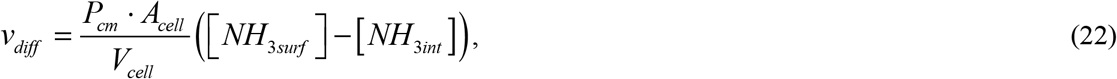

where *P*_*cm*_ is the permeability coefficient of NH_3_ through the cytoplasmic membrane, [*NH*_3*surf*_] is the NH_3_ concentration on the cell surface, and [*NH*_3*int*_] is the intracellular NH_3_ concentration. *A*_*cell*_ and *V*_*cell*_ are the cell surface area and the cell volume, respectively. A positive *v*_*diff*_ value indicates that NH_3_ is moving into the cell, whereas a negative value indicates the reverse. The net NH_x_ transport rate (*v*_*net*_) is given by

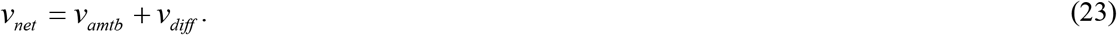

Under steady-state conditions, *v*_*net*_ is equal to the rate of ammonium assimilation, i.e., *v*_*net*_ = *v*_*gdh*_ + *v*_*gs*_.

*r*_*amtb*_ represents the average number of times a single NH_x_ molecule is transported via AmtB before being assimilated inside the cell:

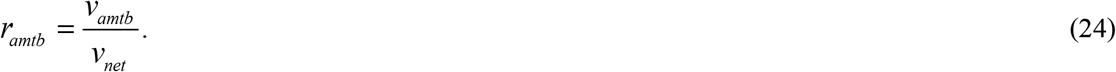

When *r*_*amtb*_ = 0, all intracellular NH_x_ is supplied through passive NH_3_ inward diffusion and subsequent re-protonation. When 0 < *r*_*amtb*_ < 1, both AmtB-mediated NH_4_^+^ transport and passive NH_3_ inward diffusion contribute to NH_4_^+^ acquisition. When *r*_*amtb*_ = 1, all NH_4_^+^ is acquired through AmtB-mediated NH_4_^+^ transport. Under these conditions (*r*_*amtb*_ ≤ 1), no futile cycling occurs. When *r*_*amtb*_ >1, futile cycling occurs because multiple AmtB-mediated NH_4_^+^ transport events occur before NH_4_^+^ is assimilated. *r*_*amtb*_ depends on the extracellular NH_x_ level. When varying the extracellular NH_x_ concentration, the maximum value of *r*_*amtb*_ under the condition that the cell is growing (μ ≥ 0.05 h^-1^) is defined as the physiological maximum value of *r*_*amtb*_ (*r*_*amtb,max*_). By comparing *r*_*amtb,max*_ under different environmental and physiological parameters, we can discuss how these parameters influence maximal futile cycling, i.e. futile cycling at the extracellular NH_x_ concentration that favors futile cycling most. *NH*_*xext,futile*_ will be used to indicate the highest NH_x_ concentration at which futile cycling occurs.

Just like *r*_*amtb*_, we define *r*_*gdh*_, *r*_*gog*_, and *r*_*gs*_ as the ratios of the respective metabolic fluxes (via GDH, GOGAT, and GS) to the ammonium assimilation rate. Comparing these values allows us to quantify the contribution of metabolic futile cycling:

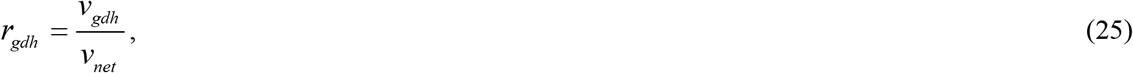

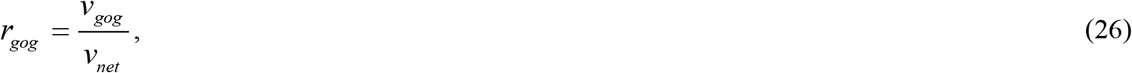

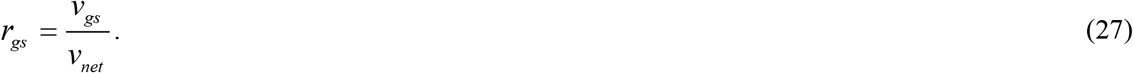

Additionally, to quantify signal transduction futile cycling, we define *r*_*ut*_ and *r*_*ad*_, representing the relative reaction rates of UTase-mediated uridylylation and ATase-mediated adenylylation, respectively, normalized by the ammonium assimilation rate:

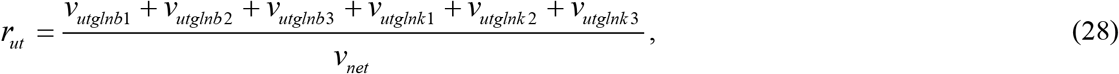

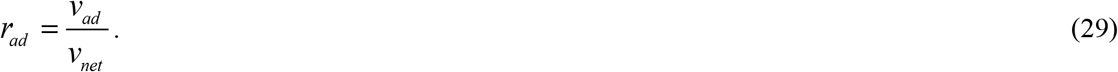

In our model, the synthesis rates of glutamate and glutamine are balanced at steady state by the consumption rates of these molecules in biosynthetic pathways. The consumption rates of these two anabolic substrates are modeled as constant multiples of the specific growth rate. These numbers are assumed to be fixed and are denoted by *GLU*_*demn*_ and *GLU*_*demf*_ for glutamate consumption and by *GLN*_*demn*_ and *GLN*_*demf*_ for glutamine (**Table S4** in Supplementary Tables). *GLU*_*demn*_ and *GLN*_*demn*_ are the biosynthetic requirements of glutamate and glutamine for transamination reactions leading to new cell mass, whilst *GLU*_*demf*_ and *GLN*_*demf*_ refer to the biosynthetic requirements of the entire molecules themselves into the biomass. Accordingly, the specific growth rate (μ) and doubling time (*τ*) are functions of the concentrations of the two metabolites. Here we follow the procedure developed by Yuan et al. ^23^:

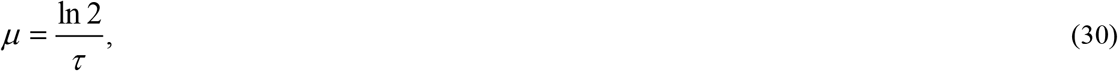

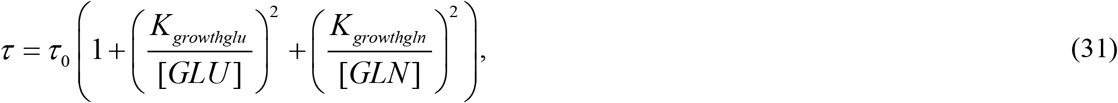

where *τ*_0_, *K*_*growthglu*_, and *K*_*growthgln*_ are model parameters (**Table S4** in Supplementary Tables). Although this growth rate function is phenomenological, **Figs. S2**-**S3** (Supplementary Figures) show that it successfully explained the experimental results from batch cultures on agarose gel media ^23^ and continuous cultures in microfluidic chambers ^22^. The relative magnitudes of *K*_*growthglu*_ and *K*_*growthgln*_ affect the relative extents to which glutamate and glutamine will be increased so as to bring the specific growth rate up to a balance with the synthesis rates of the two amino acids. In our study, we tuned the values of *K*_*growthglu*_ and *K*_*growthgln*_ in the parameter estimation step to fit the experimental data ^22-23^.

At the metabolic steady states that we model, the net uptake of nitrogen will be equal to the assimilation rate of nitrogen into new biomass, as there are no other reactions that produce nitrogenous compounds that are secreted:

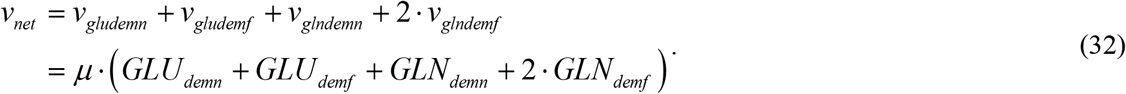

This indicates that the number of nitrogen atoms per liter of cell volume (*N*_0_) remains constant under all conditions investigated:

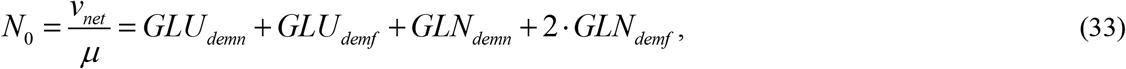

In our model, N_0_ is 3.3 mol/L, which corresponds to 122 g_DW_/mol or 11.5 % (g_N_/g_DW_), assuming an aqueous cytoplasmic volume of 2.5 mL/g_DW_^59.^The nitrogen content of 11.5 % (g_N_/g_DW_) is in good agreement with the experimentally reported value of 10.8 % (g_N_/g_DW_) for ammonium-limited cells ^60^, thereby supporting the validity of our model.

## Data Availability

The authors declare that the data supporting the findings of this study are available within the paper and its supplementary information files.

## Code Availability

Custom C and MATLAB codes used in this study are available upon request. The SBML file for the EB-Int-*φ* model will be made available from the BioModels database upon publication of the final article.

## Acknowledgements

This work was supported by Grant-in-Aid for Scientific Research (C) (22K12247) and Grant-in-Aid for Scientific Research (B) (23K24943) from the Japan Society for the Promotion of Science, and PRESTO (JPMJPR20K8) and CREST (JPMJCR20S1) from the Japan Science and Technology Agency. Additional support was provided by the Systems Biology group of the Faculty of Science, VU, and by research funding from the Foundation of the Amano Institute of Technology.

## Author contributions

K.M., H.V.W., H.K., and F.C.B. conceived the study. K.M. ran simulations with guidance and input from all authors. K.M., H.V.W., and F.C.B. wrote the manuscript with input from H.K. and A.J.. All authors reviewed the manuscript.

## Additional information

### Supplementary information

Supplementary Notes, Supplementary Tables, and Supplementary Figures are available at the journal website.

## Supplementary Notes

### 1. Modeling gene expression

The promoter activities of *amtB* and *glnA* with respect to extracellular NH_4_^+^ concentrations were reported in Supplementary Tables 6–11 of [1], where the activities were presented as fluorescence intensities of GFP and mCherry, respectively. As in our previous study [2], when developing continuous functions describing promoter activities as a function of extracellular NH_4_^+^ concentrations, we employed an approach similar to the one that Kim et al. used [1]. Specifically, we fitted the following Hill function to the *amtB* promoter activity data:

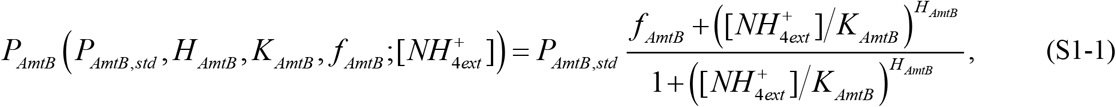

where *P*_*AmtB,std*_ represents the promoter activity at a high extracellular NH_4_^+^ concentration, while *H*_*AmtB*_, *K*_*AmtB*_, *f*_*AmtB*_ are the Hill coefficient, half-inhibition concentration, and maximal fold change, respectively. These parameters vary depending on the strain and carbon source. The fitted parameters are listed in **Table S5** in Supplementary Tables.

Using the promoter activity function above, the AmtB concentration is calculated assuming it is proportional to the promoter activity. Wild-type cells grown in medium containing 4 μM NH_4_^+^ as the nitrogen source and ample glucose as the carbon source are considered as a reference condition. Under this condition, the promoter activity corresponds to AmtB concentration [*AmtB*_*total*_]^*^ (**Table S4** in Supplementary Tables). Thus, the AmtB concentration at any extracellular NH_4_^+^ concentration was obtained using:

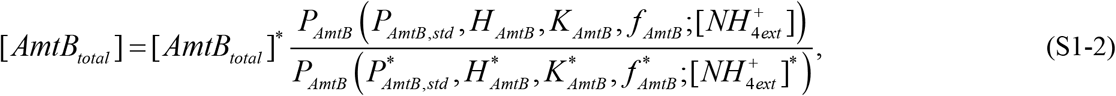

where the asterisks indicate values for the reference condition. Using Eq. (S1-1) and **Table S5**, AmtB concentrations were calculated for any extracellular NH_4_^+^ concentrations.

Since the *glnK* gene is located on the same operon as *amtB*, we assume that the GlnK concentration is also proportional to the *amtB* promoter activity:

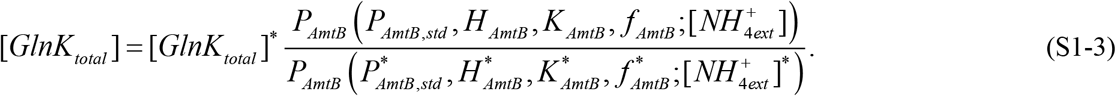

[*GlnK*_*total*_]^*^ is a GlnK concentration estimated in this study (**Table S4**).

Similarly, the following Hill function was fitted to GS promoter activity data from Kim’s study [1]:

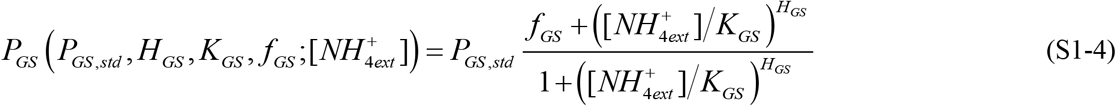

As with AmtB, we assume that for wild-type cells grown under the reference condition (4 μM NH_4_^+^ and ample glucose), the promoter activity corresponds to a reference GS concentration [*GS*_*total*_]^*^. *P*_*GS,std*_, *H*_*GS*_, *K*_*GS*_, *f*_*GS*_ are fitted and listed in **Table S5**. The GS concentration at arbitrary extracellular NH_4_^+^ concentrations was then calculated:

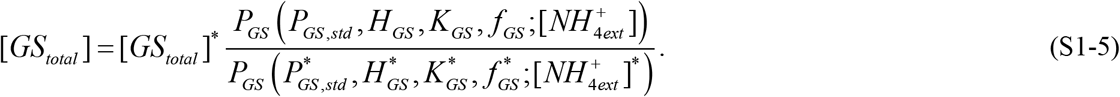

[*GS*_*total*_]^*^ is a GS concentration estimated in this study (**Table S4**).

### 2. Modeling 2-oxoglutarate

The 2-oxoglutarate concentration was not reported in [1]. Thus, in our previous study [2], we developed a function for the dependence of the 2-oxoglutarate concentration on the NH_4_^+^ concentration. We took the same approach in the present study.

Kim predicted a change in 2-oxoglutarate level with respect to the extracellular NH_4_^+^ (Figure 5C of [1]). The 2-oxoglutarate level was predicted to be insensitive to the extracellular NH_4_^+^ level if the extracellular NH_4_^+^ was higher than a certain level indicated as 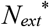, i.e. the external NH_4_ ^+^ level below which AmtB was activated. If the extracellular NH_4_^+^ was less than 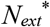, the 2-oxoglutarate level increased as the extracellular NH_4_^+^ decreased. Kim also provided an equation that relates the 2-oxoglutarate level to the intracellular NH_4_^+^ concentration (Eq. (S29) of [1]). When the intracellular NH_4_^+^ is below 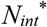, i.e. the internal NH_4_^+^ level below which AmtB is activated, the 2-oxoglutarate level increased as the intracellular NH_4_^+^ decreased. Based on Kim’s predictions, we developed a function of the 2-oxoglutarate concentration with respect to the intracellular NH_4_^+^ concentration:

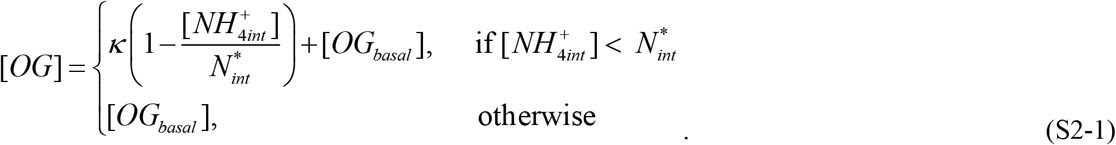

In Eq. (S2-1), there are three parameters 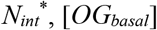, and *κ*. 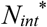 depends on the carbon source: 0.019 mM for glycerol, 0.033 mM for glucose, and 0.057 mM for G6P + gluconate (Table S5 of [1]). [*OG*_*basal*_] and *κ* are assumed to be independent of carbon sources and strains. [*OG*_*basal*_] is the 2-oxogluratate concentration under N-rich condition, and *κ* is the proportional constant. They are class II parameters, and their reference values are 0.7 mM and 10.9 mM, respectively. These reference values were chosen to make the range of [*OG*] congruent with that reported in [3, 4].

### 3. Variants of electro-binding and electro-flipping models

In the electro-binding model (*k*_*catamtbf*_ / *k*_*catamtbr*_ = 1 and *K*_*amtbnhint*_ / *K*_*amtbnhext*_ = *φ*), different model variants can be defined depending on whether the transmembrane electric potential affects the periplasmic or cytoplasmic AmtB-NH_4_^+^ binding site. In this model, the parameters *k*_*catamtbf*_, *k*_*catamtbr*_, *K*_*amtbnhext*_, and *K*_*amtbnhint*_ in Eq. (1) of Main Text are specified as follows:

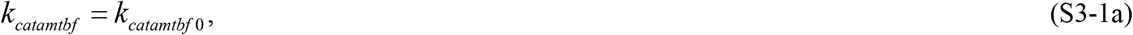

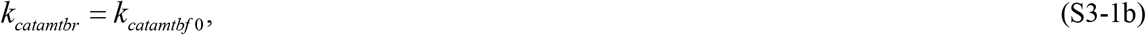

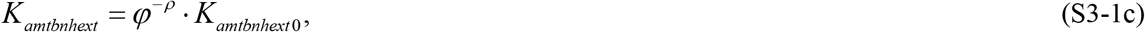

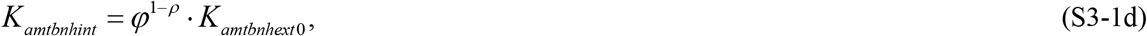

where *k*_*catamtbf*0_ and *K*_*amtbnhext*0_ denote the values of *k*_*catamtbf*_ and *K*_*amtbnhext*_, respectively, for *φ* = 1 (i.e., zero membrane potential). The actual value of *k*_*catamtbf*0_ is determined by parameter estimation and should be within the previously reported range (For reported values, see Section 4). The parameter *ρ* determines how the membrane potential affects *K*_*amtbnhext*_ and *K*_*amtbnhint*_, and takes values in the range 0 ≤ *ρ* ≤ 1. When *ρ* = 0, the membrane potential affects only *K*_*amtbnhint*_, whereas when *ρ* = 1, it affects only *K*_*amtbnhext*_. For a fair comparison of the electro-binding models with different *ρ* values, we assume that, regardless of the value of *ρ*, the value of *K*_*amtbnhext*_ under the reference condition (Δφ = -150 mV, *T* = 310 K and *φ* = 275) is equal to the previously reported value (5 μM [5-7]). To achieve this, the value of *K*_*amtbnhext*0_ (i.e., the value of *K*_*amtbnhext*_ at *φ* = 1) is determined using the following equation for different *ρ* values:

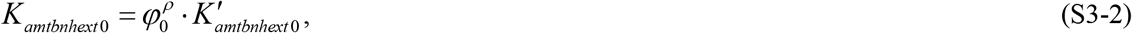

where *φ*_0_ is the accumulation factor under the reference condition, i.e., *φ*_0_ = 275. *K’*_*amtbnhext*0_ indicates *K*_*amtbnhext*_ under the reference condition, and the actual value is 5 μM (as we assumed above). As indicated in Eq. (S3-2), *K*_*amtbnhext*0_ is different for different *ρ* values. This allows for a fair comparison of the electro-binding models with different *ρ* values: Under the reference condition (*φ* = *φ*_0_), *K*_*amtbnhext*_ = *K’*_*amtbnhext*0_ and *K*_*amtbnhint*_ = *φ*_0_ *K’*_*amtbnhext*0_ irrespective of values of *ρ*. In this study, we specifically consider *ρ* = 0 (EB-Int-*φ*), 0.5 (EB-Sym-*φ*), and 1 (EB-Ext-*φ*) cases. **Fig. S1a-c** (Supplementary Figures) illustrates how *K*_*amtbnhext*_ and *K*_*amtbnhint*_ depend on *φ* and *ρ. K*_*amtbnhext*0_ is 5, 83, and 1375 μM for the EB-Int-*φ*, EB-Sym-*φ*, and EB-Ext-*φ* models, respectively. For simplicity, the case-specific *K*_*amtbnhext*0_ constants are denoted as *K*_*EB-Int-φ*_, *K*_*EB-Sym-φ*_, and *K*_*EB-Ext-φ*_ in the section of “Two model classes: potential-dependent ammonium binding affinity or potential-dependent conformational flip” in Main Text.

In the electro-flipping model (*k*_*catamtbf*_ / *k*_*catamtbr*_ = *φ* and *K*_*amtbnhint*_ / *K*_*amtbnhext*_ = 1), the parameters *k*_*catamtbf*_, *k*_*catamtbr*_, *K*_*amtbnhext*_, and *K*_*amtbnhint*_ in Eq. (1) of Main Text are given as follows:

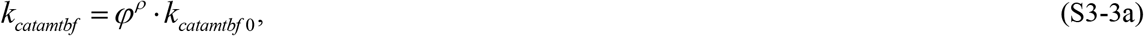

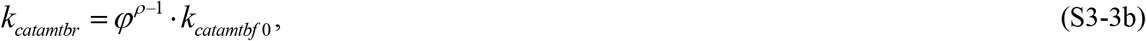

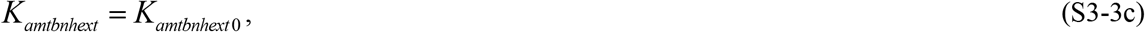

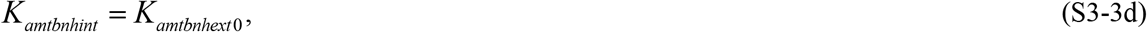

where *K*_*amtbnhext*0_ = 5 μM [5-7], and *k*_*catamtbf*0_ denotes the value of *k*_*catamtbf*_ for *φ* = 1 (i.e. zero membrane potential). As in the electro-binding model, *ρ* (0 ≤ *ρ* ≤ 1) determines how the membrane potential affects the model parameters. When *ρ* = 0, the membrane potential affects only *k*_*catamtbr*_, whereas when *ρ* = 1, it affects only *k*_*catamtbf*_. For a fair comparison of the electro-flipping models with different *ρ* values, we assume that, regardless of the value of *ρ*, the value of *k*_*catamtbf*_ under the reference condition takes the identical value. To achieve this, the value of *k*_*catamtbf*0_ (i.e., the value of *k*_*catamtbf*_ at *φ* = 1) is determined using the following equation.

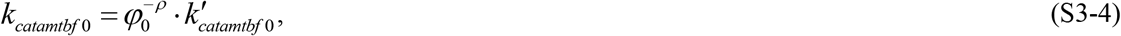

where *φ*_0_ is the accumulation factor under the reference condition, i.e., *φ*_0_ = 275. *k’*_*catamtbf*0_ corresponds to *k*_*catamtbf*_ under the reference condition and is determined by parameter estimation. Under the reference condition (*φ* = *φ*_0_), *k*_*catamtbf*_ = *k’*_*catamtbf*0_ and *k*_*catamtbr*_ = *φ*_0^−1^_ *k’*_*catamtbf*0_ irrespective of values of *ρ* . In this study, we specifically consider *ρ* = 0 (EF-Rev-*φ*), 0.5 (EF-Sym-*φ*), and 1 (EF-Fwd-*φ*) cases. **Fig. S1d-f** (Supplementary Figures) illustrates how *k*_*catamtbf*_ and *k*_*catamtbr*_ depend on *φ* and *ρ. k*_*catamtbf* 0_ is 1.91 × 10^6^, 1.13 × 10^5^, and 7.10 × 10^3^ min^-1^ for the (best) EF-Rev-*φ*, EF-Sym-*φ*, and EF-Fwd-*φ* models, respectively. For simplicity, the case-specific *k*_*catamtbf* 0_ constants are denoted as *k*_*EF-Rev-φ*_, *k*_*EF-Sym-φ*_, and *k*_*EF-Fwd-φ*_ in the section of “Two model classes: potential-dependent ammonium binding affinity or potential-dependent conformational flip” in Main Text.

### 4. Further discussion on *k*_*catamtbf*_, and AmtB and GlnK concentrations

According to Radchenko et al. [8], the total AmtB concentration is 1353 trimers per cell under N-limited condition. For the electro-binding model, the total AmtB concentration was estimated to be 1.8 μM (2357 trimers per cell) whereas it was 2.8 μM (3631 trimers per cell) for the electro-flipping model under the reference condition (i.e., Kim’s experiment with NH_4_ ^+^_ext_ = 4 μM and glucose as a carbon source). The number of AmtB monomers for the electro-flipping models is comparable to or even exceeds that of the ATP synthetase—one of the most abundant transporter proteins in *E. coli*, with approximately 10000 ATP synthetase complexes per cell at the specific growth rate of 0.8 h^-1^ (Circles in Figure 3F of [9]). Therefore, both the electro-binding and electro-flipping models require total AmtB concentrations that are higher than the AmtB level reported by Radchenko et al. [8]. However, the required concentration is substantially higher for the electro-flipping models, reaching a level comparable to that of the ATP synthetase complex.

van Heeswijk et al. [10] reported that the GlnK concentration was 1.7 times higher than GlnB under N-limited conditions while Gosztolai et al. [11] reported higher ratios of 3.8. In our study, the GlnK/GlnB ratio was 3.3 and 5.6 for the electro-binding and the electro-flipping models, respectively. Therefore, the GlnK/GlnB ratio for the electro-binding models falls within the reported range, whereas that for the electro-flipping models exceeds it.

Khademi et al. [12] estimated AmtB single channel conductance to be ∼27000 s^-1^ (see Table 3 of ref [6]). However, as pointed out by Javelle et al. [6], the proteo-liposome experiments by Khademi et al. [12] were not reproducible, and Javelle et al. [6] speculated that the single channel conductance is 10–100-fold lower than the value reported by Khademi et al. [12]. Thus, we took 2700 s^-1^ (per monomer), i.e. 4.86 × 10^5^ min^-1^ (per trimer) as the reference value for *k*_*catamtbf*_. This value is within the range estimated by Zheng et al. [13]: 1.8 × 10^3^–1.8 × 10^6^ min^-1^ per trimer. In the electro-binding and electro-flipping models, *k*_*catamtbf*_ was estimated to be 7.49 × 10^5^ min^-1^ and 1.92 × 10^6^ min^-1^, respectively. The value for the electro-flipping models is 2.6-fold higher than that for the electro-binding models and exceeds the ranges suggested by Javelle et al. [6] and Zheng et al. [13].

### 5. Disruption of coordinated regulation of GS and AmtB

The coordinated regulation of GS and AmtB is achieved through the uridylylation of GlnB and GlnK and their binding to 2-oxoglutarate. To explore what happens when these regulatory mechanisms fail, we compared the wild-type strain with four virtual mutants, each representing a specific disruption in this regulation. The results are shown in **Fig. S6** in Supplementary Figures. The metabolic steady state reached after a mutation is introduced into *E. coli* cells that were first grown at the NH_x_ concentrations indicated on the x-axis. For this reason, the gene expression levels of GS, AmtB, and GlnK are identical across all strains.

In the “loss of deuridylylation” mutant (red dotted lines in **Fig. S6a, c, e, h**), UTase lacks uridylyl-removing activity (*k*_*caturglnb*_ = 0 and *k*_*caturglnk*_ = 0), and uridylylated GlnK cannot bind to AmtB. As a result, AmtB-mediated NH_4_^+^ transport occurs continuously. This mutant maintains a high growth rate (μ) across a broad NH_x_ range (**Fig. S6c**), but *r*_*amtb*_ (= *v*_*amtb*_/*v*_*net*_) exceeds 20 (**Fig. S6a**), indicating excessive futile cycling and unnecessary free-energy consumption compared with the wild type.

In the “loss of uridylylation” mutant (yellow dotted lines in **Fig. S6a, c, e, h**), UTase lacks uridylyl transferase activity, or GlnB/GlnK harbor mutations that prevent uridylylation (*k*_*catutglnb*_ = 0 and *k*_*catutglnk*_ = 0). As a result, GS remains almost fully adenylylated and therefore nearly completely inactive (**Fig. S6e**). *r*_*amtb*_ remains zero expect at 50-60 µM NH_x_ where μ and thus *v*_*net*_ are very low but still *µ* > 0.05 h^-1^ (**Fig. S6a**). However, since GS is inactive, growth is severely impaired, and at NH_x_ concentrations below 50 µM, growth is essentially absent (**Fig. S6c**).

The “strong 2-oxoglutarate binding” mutant (purple dotted lines in **Fig. S6b, d, f, i**) shows a phenotype broadly similar to the “loss of deuridylylation” mutant. This mutant retains normal uridylylation and deuridylylation activity, but GlnK and GlnB have mutations in their 2-oxoglutarate binding sites, making them bind 2-oxoglutarate much more strongly (*K*_*glnbog,i*_ → 0, *K*_*glnbumpog,i*_ → 0, and *K*_*glnkog,i*_ → 0 for *i* = 1, 2, 3). The “strong 2-oxoglutarate binding” mutant maintains a high growth rate (μ) across a broad NH_x_ range (**Fig. S6d**), but *r*_*amtb*_ (= *v*_*amtb*_/*v*_*net*_) exceeds 20 (**Fig. S6b**).

In the “no 2-oxoglutarate binding” mutant (green dotted lines in **Fig. S6b, d, f, i**), GlnK and GlnB cannot bind 2-oxoglutarate (*K*_*glnbog,i*_ → ∞, *K*_*glnbumpog,i*_ → ∞, and *K*_*glnkog,i*_ → ∞ for *i* = 1, 2, 3). In this case, GS activation proceeds almost normally (**Fig. S6f**), allowing rapid growth at NH_x_ concentrations above 60 μM (**Fig. S6d)**. However, AmtB remains inactive even below 60 μM NH_x_ (**Fig. S6i**), resulting in reduced growth under nitrogen-limited conditions where AmtB activity is normally required.

These results highlight the critical role of coordinated GS and AmtB regulation in balancing growth and energy efficiency. Without proper regulation, *E. coli* either fails to sustain growth or wastes free energy through excessive futile cycling.

### 6. Further discussion on control coefficients

As shown in **Table S7** in Supplementary Tables, control coefficients of *τ*_0_ (the minimal doubling time) have large absolute values for glutamate (*GLU*) and glutamine (*GLN*) concentrations, rates of GOGAT and GS reactions (*v*_*gog*_ and *v*_*gs*_), the ammonium assimilation rate (*v*_*net*_), specific growth rate (μ), and *r*_*amtb*_. This is expected because μ and the consumption rates of glutamate and glutamine are functions of *τ*_0_ [Eqs. (30)-(31) in Main Text].

When *NH*_*xext*_ = 1 mM, the control coefficients of *k*_*catamtbf*_ with respect to *v*_*amtb*_ and *r*_*amtb*_ are 1.002 (**Table S7**). These control coefficients are expected to be high because, at *NH*_*xext*_ = 1 mM, 2-oxoglutarate is insensitive to the intracellular NH_4_^+^ level (see Section 2), meaning that 2-oxoglutarate-based feedback regulation of AmtB transport is absent. It should be noted, however, at *NH*_*xext*_ = 1 mM, AmtB expression is very low, and thus these high control coefficients are of little biological relevance. By contrast, at *NH*_*xext*_ = 10 µM, an increase in intracellular NH_4_^+^ reduces 2-oxoglutarate concentration, which promotes GlnK binding to AmtB and suppresses NH_4_^+^ transport. This feedback loop effectively reduces the control coefficients of *k*_*catamtbf*_ with respect to *v*_*amtb*_ and *r*_*amtb*_ to 0.262 and 0.270, respectively, well below 1.002 (**Table S7**).

### 7. Effects of changes in cell area-to-volume ratio

The rate of unmediated NH_3_ diffusion flux (in mM/min) is proportional to the surface-to-volume ratio of the cells, *A*_*cell*_/*V*_*cell*_ ratio [see Eq. (22) in Methods of Main Text]. For the “normal cell”, we used a surface area (*A*_*cell*_) of 9.18 μm^2^ and a volume (*V*_*cell*_) of 2.15 μm^3^, values representative of *E. coli* growing exponentially on glucose [14].

To assess the importance of *A*_*cell*_/*V*_*cell*_ ratio for ammonium assimilation and specific growth rate, we simulated three cell sizes:

- Small: *V*_*cell*_ = 1.14 μm^3^, *A*_*cell*_ = 5.43 μm^2^, *A*_*cell*_/*V*_*cell*_ = 4.78 μm^-1^
- Normal: *V*_*cell*_ = 2.15 μm^3^, *A*_*cell*_ = 9.18 μm^2^, *A*_*cell*_/*V*_*cell*_ = 4.27 μm^-1^
- Large: *V*_*cell*_ = 3.25 μm^3^, *A*_*cell*_ = 13.24 μm^2^, *A*_*cell*_/*V*_*cell*_ = 4.08 μm^-1^

A constant cell width of 1.08 μm was used for all three geometries, and the cell lengths were set to 1.60 μm, 2.70 μm, and 3.90 μm for the small, normal, and large cells, respectively. Cells were modeled as cylinders capped by two hemispheres (as in [15] and Section 8.2 of the Supplementary Information in our previous study [2]). The small and large sizes approximate the lower and upper bounds of *E. coli* cell dimensions observed under diverse growth conditions [15].

We maintained the number of AmtB and GlnK molecules per unit cell surface area, while keeping the concentrations of other molecules at a fixed level. For ΔAmtB cells, the changes in cell size had minimal impact on internal NH_4_^+^ and NH_3_ concentrations and growth rates (**Fig. S8a-c** in Supplementary Figures). Although reducing cell size increases the *A*_*cell*_/*V*_*cell*_ ratio and is expected to enhance passive inward NH_3_ diffusion, this strategy alone does not improve growth, because an increased *A*_*cell*_/*V*_*cell*_ ratio does not lead to the accumulation of intracellular NH_4_^+^ in the absence of AmtB. For wild-type cells (with AmtB), smaller cell sizes increased NH_3_ back diffusion and futile cycling due to higher *A*_*cell*_/*V*_*cell*_ ratio (**Fig. S8d-f**) As a result, smaller cells experienced somewhat greater free-energy loss, with *r*_*amtb*_ increased as cell size decreased (**Fig. S8g**). *NH*_*xext,futile*_ are 52 µM for all the cases (**Fig. S8h**). In summary, for ΔAmtB cells, reducing cell size to survive in low NH_x_ environments is not effective: reducing cell size would not be a good alternative to expressing AmtB. In wild-type cells, smaller sizes exacerbate NH_3_ back diffusion and futile cycling.

### 8. Effects of changes in NH_3_ permeability

The permeability coefficient of the cytoplasmic membrane (*P*_*cm*_) determines the rate at which NH_3_ permeates the membrane. For normal *E. coli* cells, we used *P*_*cm*_ of 0.073 m/min, based on experimental measurements [1, 16]. We ran simulations with *P*_*cm*_ = 0.025 m/min (one-third), 0.073 m/min (normal), and 0.220 m/min (triple). For ΔAmtB cells, changes in *P*_*cm*_ had small impact on internal NH_4_^+^ and NH_3_ concentrations or growth rates, even with a nine-fold variation in *P*_*cm*_ (**Fig. S9a-c** in Supplementary Figures). In wild-type cells, higher *P*_*cm*_ increased NH_3_ back diffusion and futile cycling (**Fig. S9d-f**), making it harder to maintain growth at low NH_x_ concentrations (**Fig. S9c**). *r*_*amtb,max*_ increased as *P*_*cm*_ increased. Lower *P*_*cm*_ reduced NH_3_ back diffusion and futile cycling, providing a clear growth advantage but only at extracellular NH_x_ concentrations as low as 1 μM. In summary, for ΔAmtB cells, increasing *P*_*cm*_ to survive in low NH_x_ environments is not effective: increasing *P*_*cm*_ would not be a good alternative to expressing AmtB. In wild-type cells, adjusting membrane composition to decrease *P*_*cm*_ could be beneficial in terms of reducing futile cycling.

### 9. Effects of changes in Michaelis constant of GS for NH_4_^+^

A value of 100 µM for *E. coli’s* GS Michaelis constant (*K*_*gsnh*_) for NH_4_^+^ is frequently used in the literature. Nevertheless other values in the range of 60-200 µM have also been reported [17]. To investigate how *K*_*gsnh*_ affects the futile cycling, we ran simulations with *K*_*gsnh*_ values of 50 µM (half), 100 µM (normal), and 200 µM (double) (**Fig. S10** in Supplementary Figures). We also changed *K*_*gsgln*_ to keep the equilibrium constant (*K*_*gseq*_) unchanged. We used the same normal AmtB and GS expression profiles for all *K*_*gsnh*_ cases. Intracellular NH_4_^+^ levels were largely unaffected by changes in *K*_*gsnh*_ (**Fig. S10a**). As the external NH_x_ concentration was decreased, *V*_*gs*_^*app*^ increased to compensate. As should be expected this compensation was stronger for the case with the higher *K*_*gsnh*_ (**Fig. S10d**). However, when *V*_*gs*_^*app*^ approached its maximum (*V*_*gs*_), further compensation failed, and growth rate decreased in the case of increased *K*_*gsnh*_ (**Fig. S10c**). *θ*_*gs*_ (reflecting GS activity via deadenylylation) varied depending on *K*_*gsnh*_ (**Fig. S10e**). The lower the *K*_*gsnh*_, the lower the external NH_x_ level at which *θ*_*gs*_ reached 1. *θ*_*gs*_ depends on the average number of adenylylated monomers (*n*_*AMP*_) in the GS dodecamer [Eqs. (18)-(19) in Main Text]. The lower the *K*_*gsnh*_, the lower the external NH_x_ level at which *n*_*AMP*_ reached zero (**Fig. S10f**). When *K*_*gsnh*_ was lower than approximately 130 µM, *r*_*amtb,max*_ remained at around 7 (**Fig. S10j**). However, when *K*_*gsnh*_ exceeded 130 µM, *r*_*amtb,max*_ increased. In the case of *K*_*gsnh*_ of 200 µM, *r*_*amtb,max*_ is high because of reduced ammonium assimilation rate (*v*_*net*_) (**Fig. S10i**). Taken together, the changes in *K*_*gsnh*_ strongly affected the growth rates at extracellular NH_x_ levels below 100 µM, the effect is only dampened partly by enhanced deadenylylation of GS at higher *K*_*gsnh*_ values.

### 10. Effects of changes in pKa of ammonium

Changes in pKa significantly affect the NH_4_^+^/NH_3_ ratio, influencing NH_x_ transport and futile cycling. pKa generally depends on temperature and solvent conditions. Here, we performed simulations varying pKa between 8.65 and 9.25, a range equivalent to a 20 °C temperature shift. Simulations showed that in ΔAmtB cells, this pKa variation had little effect on growth rate (**Fig. S11c** in Supplementary Figures). This is because passive NH_3_ diffusion remains the sole mode of NH_x_ uptake, and the internal NH_4_^+^ level remains much lower than the external one, regardless of pKa (**Fig. S11a**). In wild-type cells, a higher pKa facilitated faster growth at lower NH_x_ concentrations (**Fig. S11c**), whereas a lower pKa led to increased NH_3_ levels (**Fig. S11b**), enhancing NH_3_ back diffusion and futile cycling (**Fig. S11d-f**). *NH*_*xext,futile*_ remained 52 µM NH_x_, regardless of pKa (**Fig. S11h**), but *r*_*amtb,max*_ decreased with increasing pKa (**Fig. S11g**). Since pKa decreases with increasing temperature, this suggests that futile cycling becomes more pronounced under higher temperature conditions.

## Supplementary Figures

**Fig. S1.**
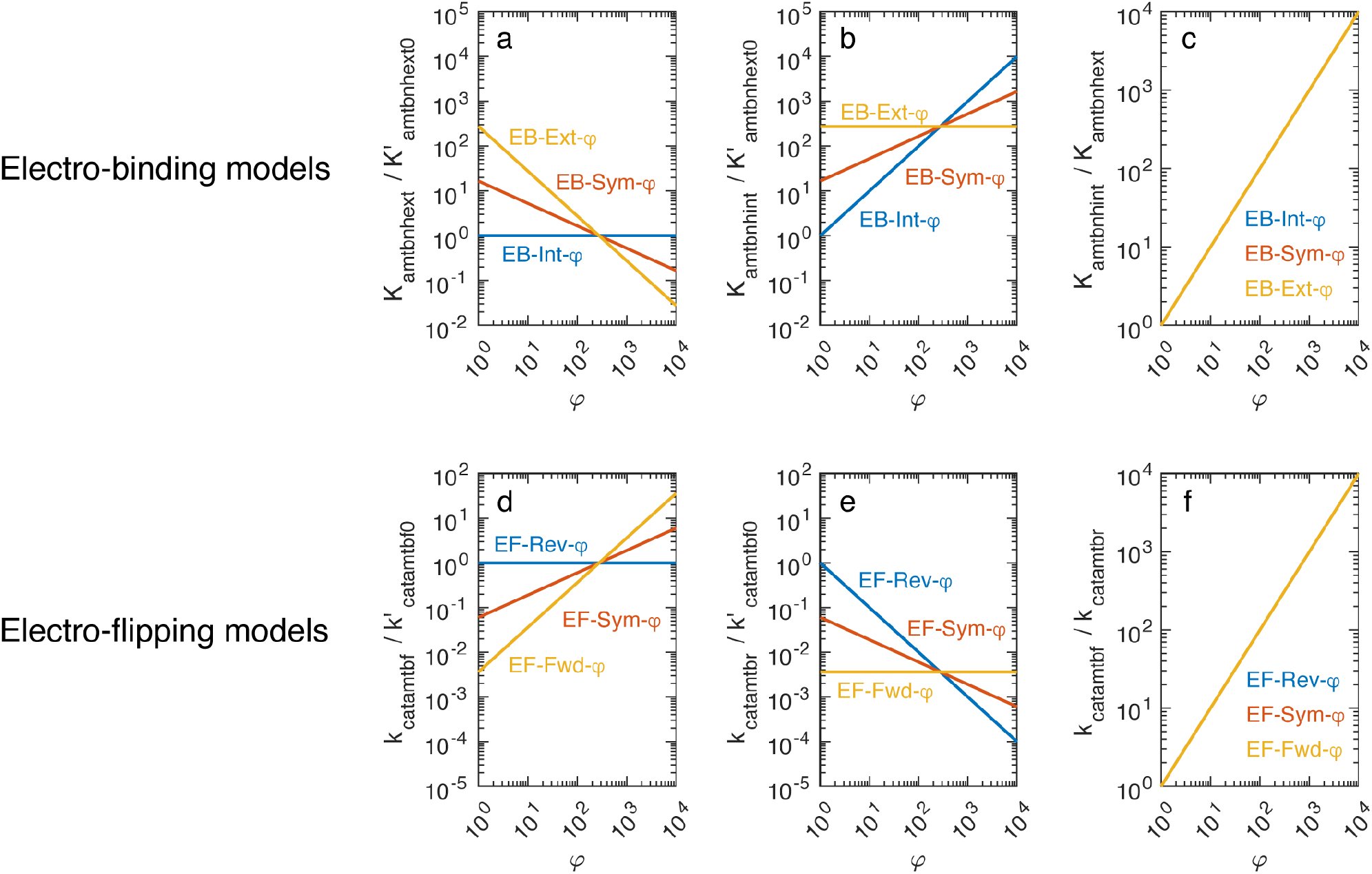
Values of AmtB-related parameters as functions of the accumulation factor of NH_4_^+^ (*φ*). (a) Normalized *K*_*amtbnhext*_ values for the electro-binding models. (b) Normalized *K*_*amtbnhint*_ values for the electro-binding models. (c) *K*_*amtbnhint*_/*K*_*amtbnhext*_ ratio for the electro-binding models. (d) Normalized *k*_*catamtbf*_ values for the electro-flipping models. (e) Normalized *k*_*catamtbr*_ values for the electro-flipping models. (f) *k*_*catamtbf*_/*k*_*catamtbr*_ ratio for the electro-flipping models. In panels (c) and (f), blue, red, and yellow lines coincide.

**Fig. S2.**
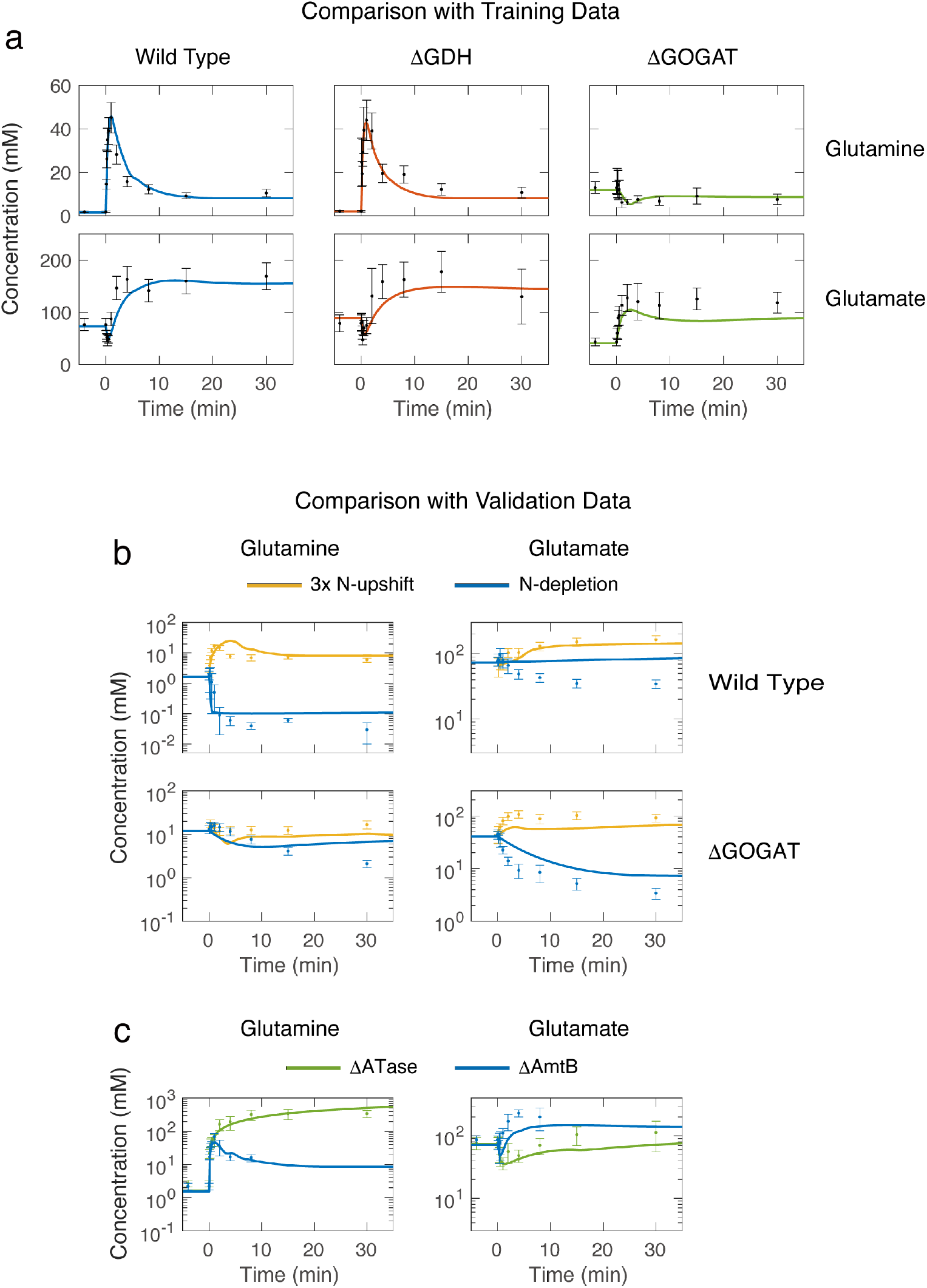
Comparison of model simulations with experimental data obtained by Yuan et al. (a) 13× N-upshift experiment, (b) various N-perturbations in the wild type and ΔGOGAT, (c) 13× N-upshift for ΔATase and ΔAmtB. Symbols and error bars represent experimental measurements as done by Yuan et al., Mol Syst Biol, 2009;5:302. Lines represent the values simulated by the EB-Int-*φ* model. At time zero, N-limited cells were transferred to plates with various NH_4_^+^ concentrations. The experimental data were obtained from Fig. 4 (13× N-upshift), Fig. 7 (various N-perturbations in the wild type and ΔGOGAT, and ΔATase), and Supplementary Fig. 5 (ΔAmtB) of Yuan et al., Mol Syst Biol, 2009;5:302. For details on the simulation settings, see Section 2.1 of Supplementary Information of our previous study (Maeda at al., NPJ Syst Biol Appl, 2019;5:14).

**Fig. S3.**
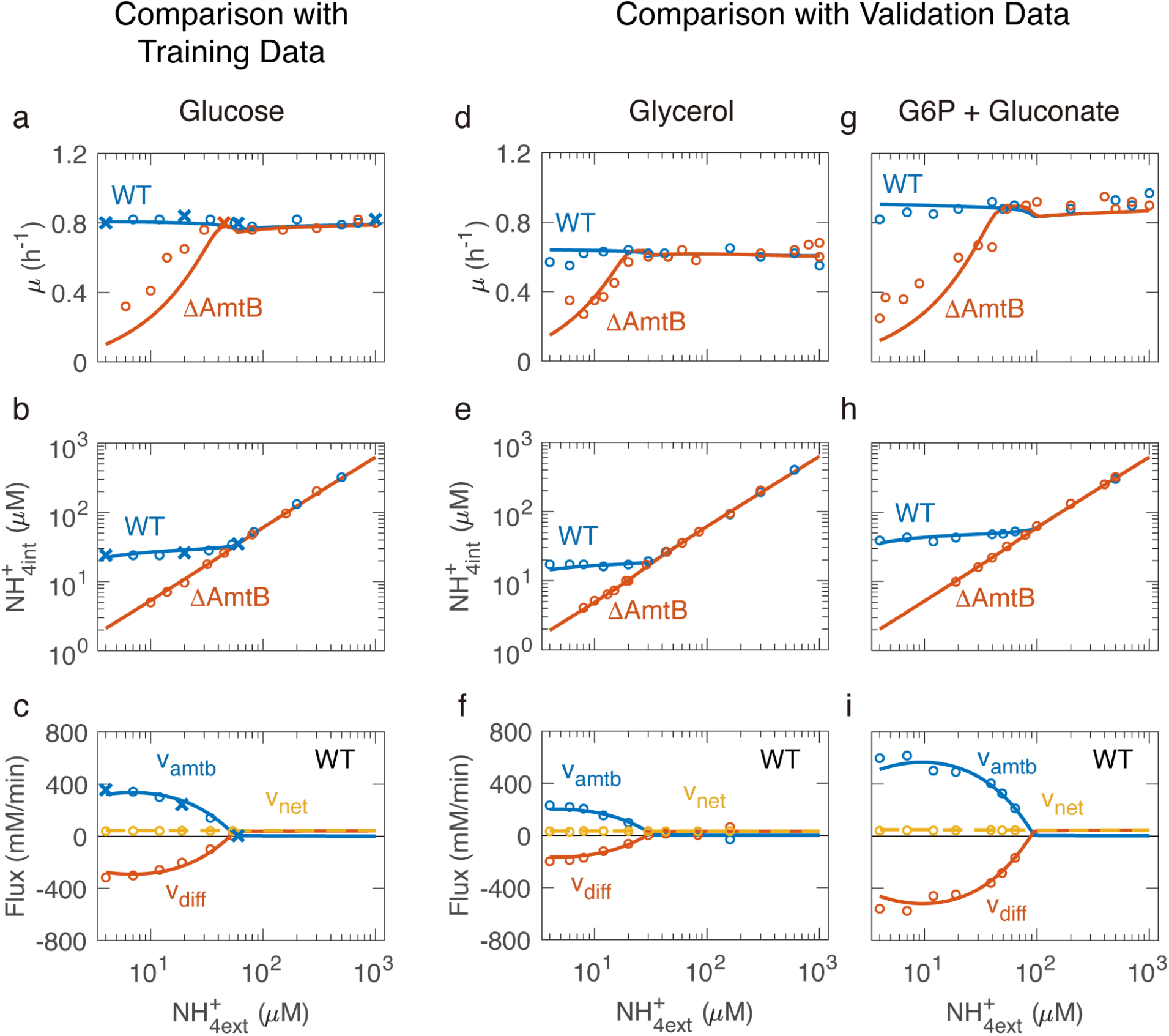
Comparison of model simulations with experimental data obtained by Kim et al. Steady-state changes of various variables as a function of extracellular ammonium. The carbon source is glucose for (a–c), glycerol for (d–f), and glucose-6P + gluconate for (g–i). Red, yellow, and blue circles represent experimental data. Cross symbols are data points used for parameter estimation. Solid lines with a corresponding color represent the values simulated by the EB-Int-*φ* model. The experimental data in (a), (d), and (g) were obtained from Fig. 3A of Kim et al., Mol Syst Biol, 2012;8:616. The experimental data in (b), (e), and (h) were obtained from Fig. 3C of Kim et al., Mol Syst Biol, 2012;8:616. The experimental data in (c), (f) and (i) were kindly provided by Dr. Minsu Kim. For details on the simulation settings, see Section 2.2 of Supplementary Information of our previous study (Maeda at al., NPJ Syst Biol Appl, 2019;5:14).

**Fig. S4.**
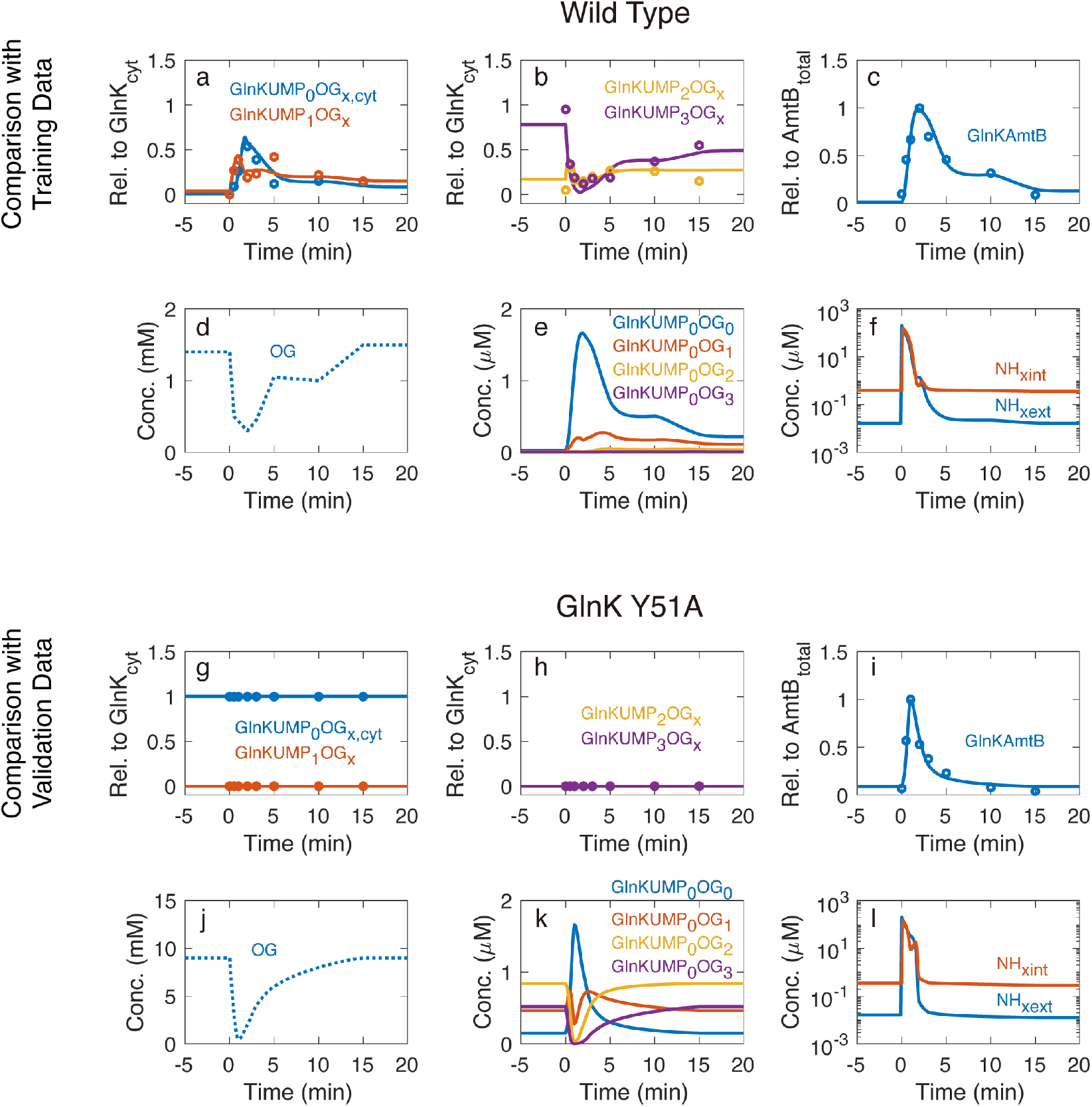
Comparison of model simulations with experimental data obtained by Radchenko et al. (a–f) The wild type and (g–l) GlnK Y51A mutant. Red, blue, yellow, and purple circles represent experimental data. Solid lines with a corresponding color represent the values simulated by the EB-Int-*φ* model. Dotted lines represent assumed dynamic model inputs. “Rel. to GlnK_cyt_” and “Rel. to AmtB_total_” indicate abundance relative to the total cytoplasmic GlnK and that to the total AmtB, respectively. GlnKUMP_0_OG_*x*,cyt_ indicates the sum of cytoplasmic GlnKUMP_0_OG_0_, GlnKUMP_0_OG_1_, GlnKUMP_0_OG_2_, and GlnKUMP_0_OG_3_. The same applies to GlnKUMP_1_OG_*x*_, GlnKUMP_2_OG_*x*_, and GlnKUMP_3_OG_*x*_. In (h), the yellow and purple lines coincide. At time zero, extracellular NH_x_ (NH_4_^+^ + NH_3_) was shifted from 15 nM to 200 μM. We obtained the experimental data in (a–c) and (g–i) by semi-quantifying blackness of the protein bands visible in Fig. 3AB of Radchenko et al., Front Microbiol, 2014;5:731 with ImageJ. We used Fig. 2A of Radchenko et al., J Biol Chem, 2010;285(40):31037-45 to obtain the 2-oxoglutarate profile for the wild type. Since 2-oxoglutarate for GlnK Y51A mutant has not been measured, we optimized the dynamic 2-oxoglutarate model input. For details on the simulation settings, see Section 2.3 of Supplementary Information of our previous study (Maeda at al., NPJ Syst Biol Appl, 2019;5:14).

**Fig. S5.**
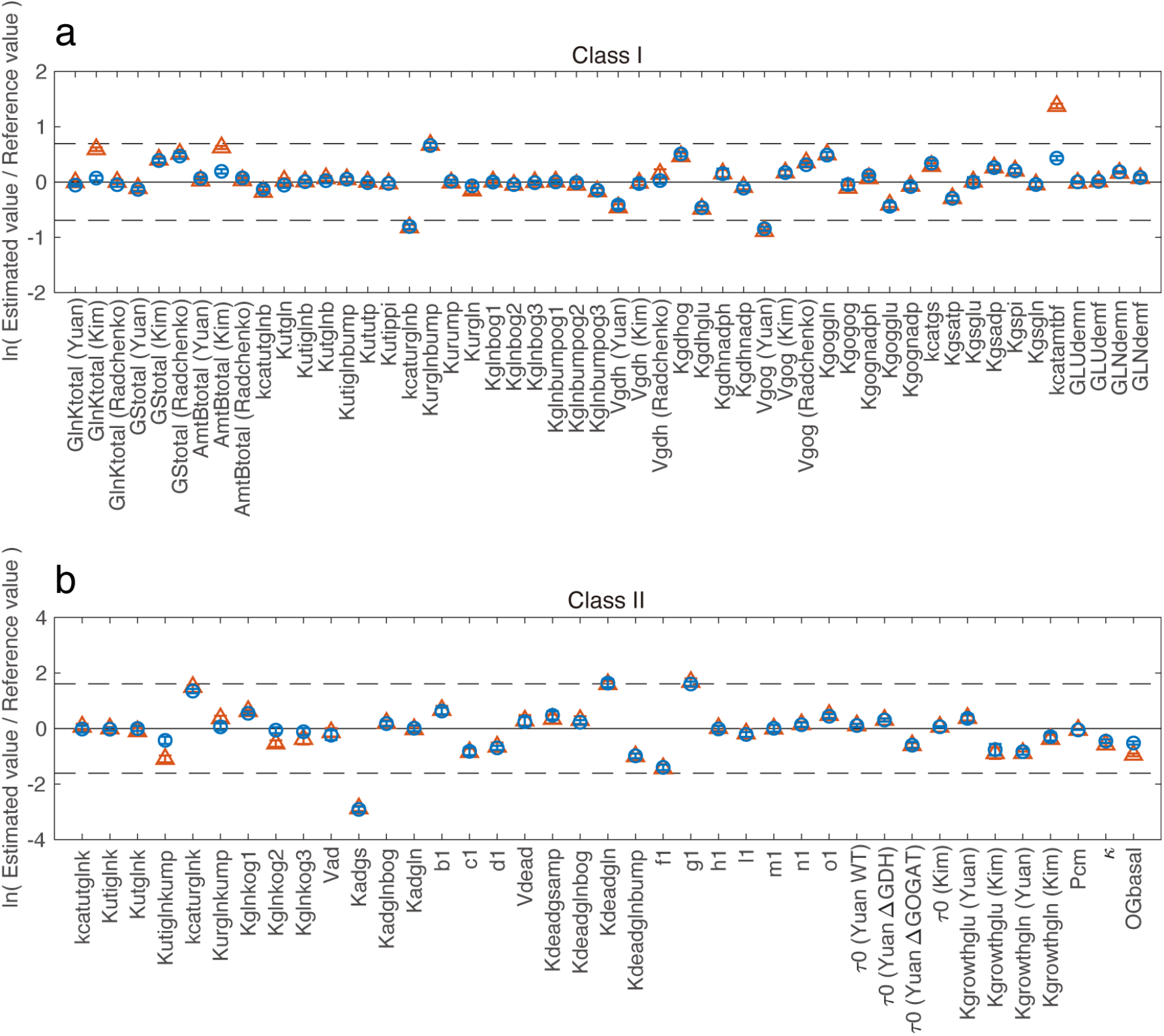
Deviation of estimated parameter values from their reference values. (a) Class I parameters. (b) Class II parameters. Blue circles and red triangles represent the electro-binding model variants (EB-Int-*φ*, EB-Sym-*φ*, and EB-Ext-*φ*) and the electro-flipping model variants (EF-Rev-*φ*, EF-Sym-*φ*, and EF-Fwd-*φ*), respectively. We repeated parameter estimation five times for each variant. Circles and triangles represent mean values (*n* = 15). Error bars represent ± standard deviation. Dashed lines show the boundaries of a twofold change in (a) and a fivefold change in (b) above and below the reference values. It should be noted that in parameter estimation we assumed that ln(*p*_*i*_/*p*_*i*\*_) follows the normal distribution with the standard deviation of ln(2) for class I and ln(5) for class II. The parameter values used for the figures are shown in **Table S6**.

**Fig. S6.**
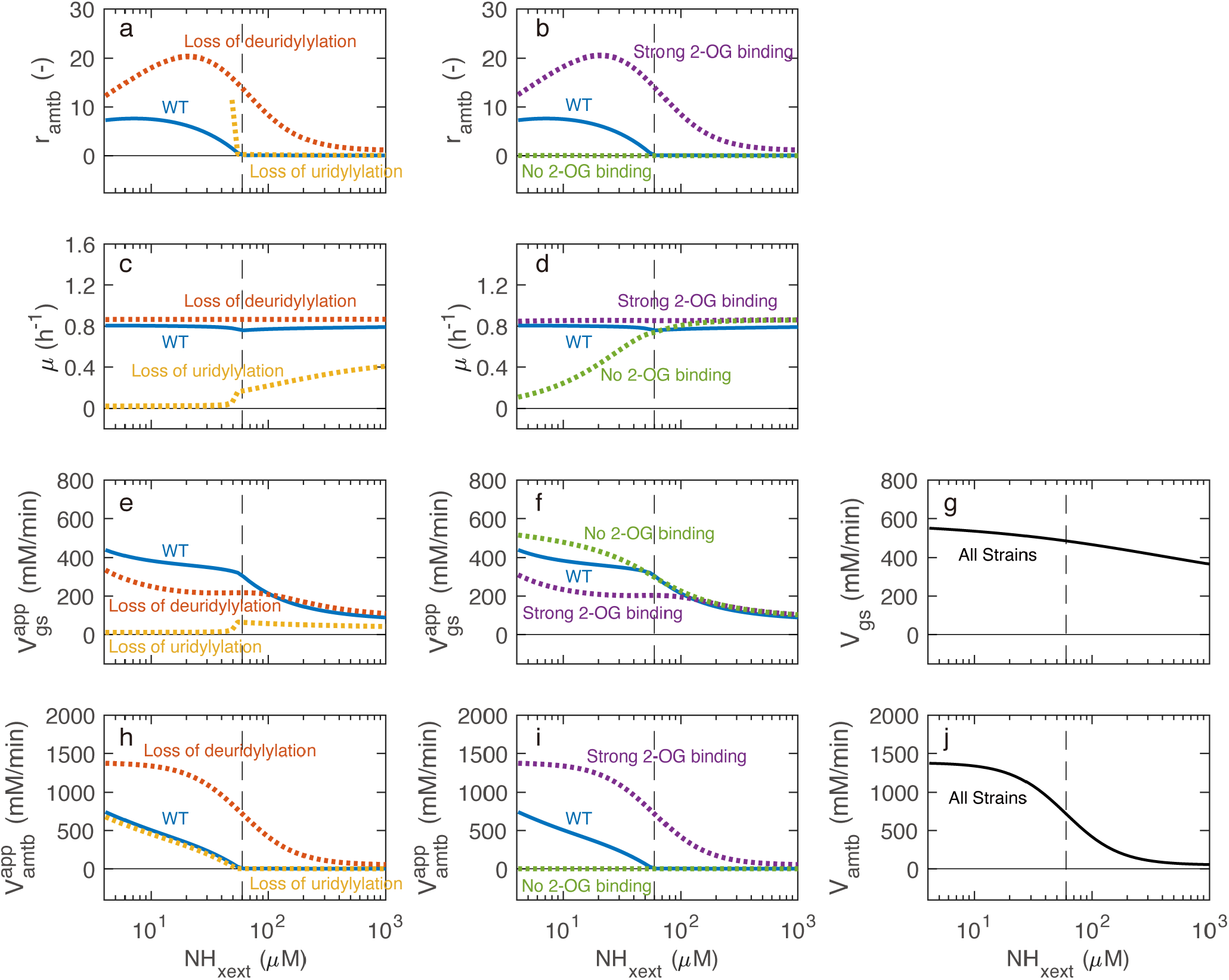
Predicted steady-state behaviors of the wild-type and virtual mutant strains. (a-b) The ratio of the AmtB-mediated NH_4_^+^ transport rate to the ammonium assimilation rate (*r*_*amtb*_). *r*_*amtb*_ is the measure of the magnitude of ammonium futile cycling and given as *r*_*amtb*_ = *v*_*amtb*_/*v*_*net*_. Note that *r*_*amtb*_ is defined only for growing cells (μ > 0.05 h^-1^) (see Methods), and thus the yellow solid line for the “loss of uridylylation” mutant is shown only for *NH*_*xext*_ > ∼50 μM. (c-d) Specific growth rate. (e-f) Apparent maximum rate of GS reaction (*V*_*gs*_^*app*^). (g) Maximum rate of GS reaction (*V*_*gs*_). (h-i) Apparent maximum rate of AmtB-mediated NH_4_^+^ transport (*V*_*amtb*_^*app*^). (j) Maximum rate of AmtB-mediated NH_4_^+^ transport (*V*_*amtb*_).

**Fig. S7.**
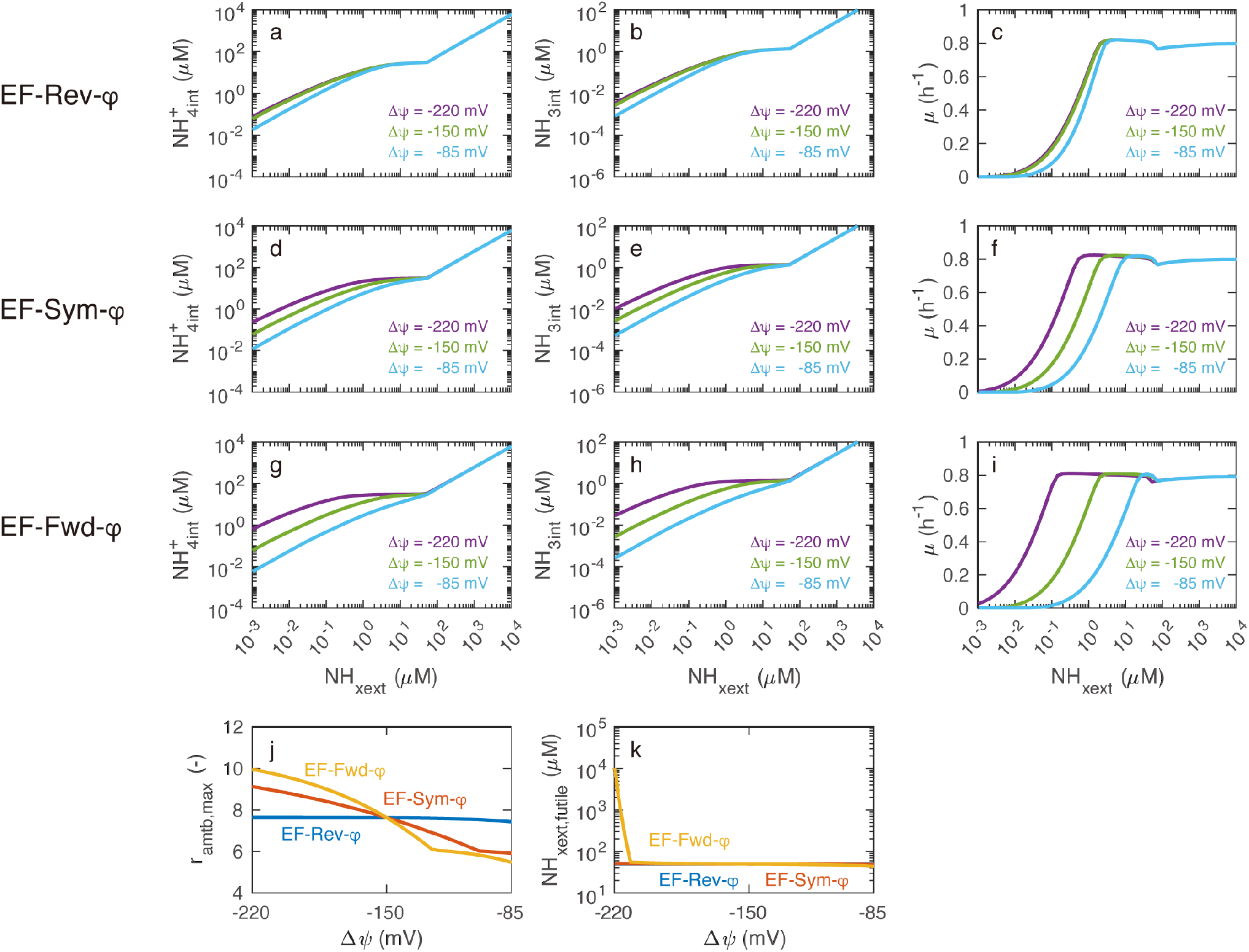
Effects of changes in transmembrane electric potential on steady-state behaviors of the electro-flipping model variants. Transmembrane electric potential was assumed to affect the conformational flip of the transporter but does not influence AmtB-NH_4_^+^ binding. EF-Rev-*φ*: Transmembrane electric potential affects the outward conformational flip. EF-Sym-*φ*: Transmembrane electric potential affects both the inward and outward conformational flip. EF-Fwd-*φ*: Transmembrane electric potential affects the inward conformational flip. (a, d, g) Intracellular NH_4_^+^ concentration. (b, e, h) Intracellular NH_3_ concentration. (c, f, i) Specific growth rate. (j) The maximum values of *r*_*amtb*_ when the external NH_x_ was changed (*r*_*amtb,max*_). (k) The highest external NH_x_ concentration at which futile cycling occurs (*NH*_*xext,futile*_). In panel (k), the blue and red lines coincide.

**Fig. S8.**
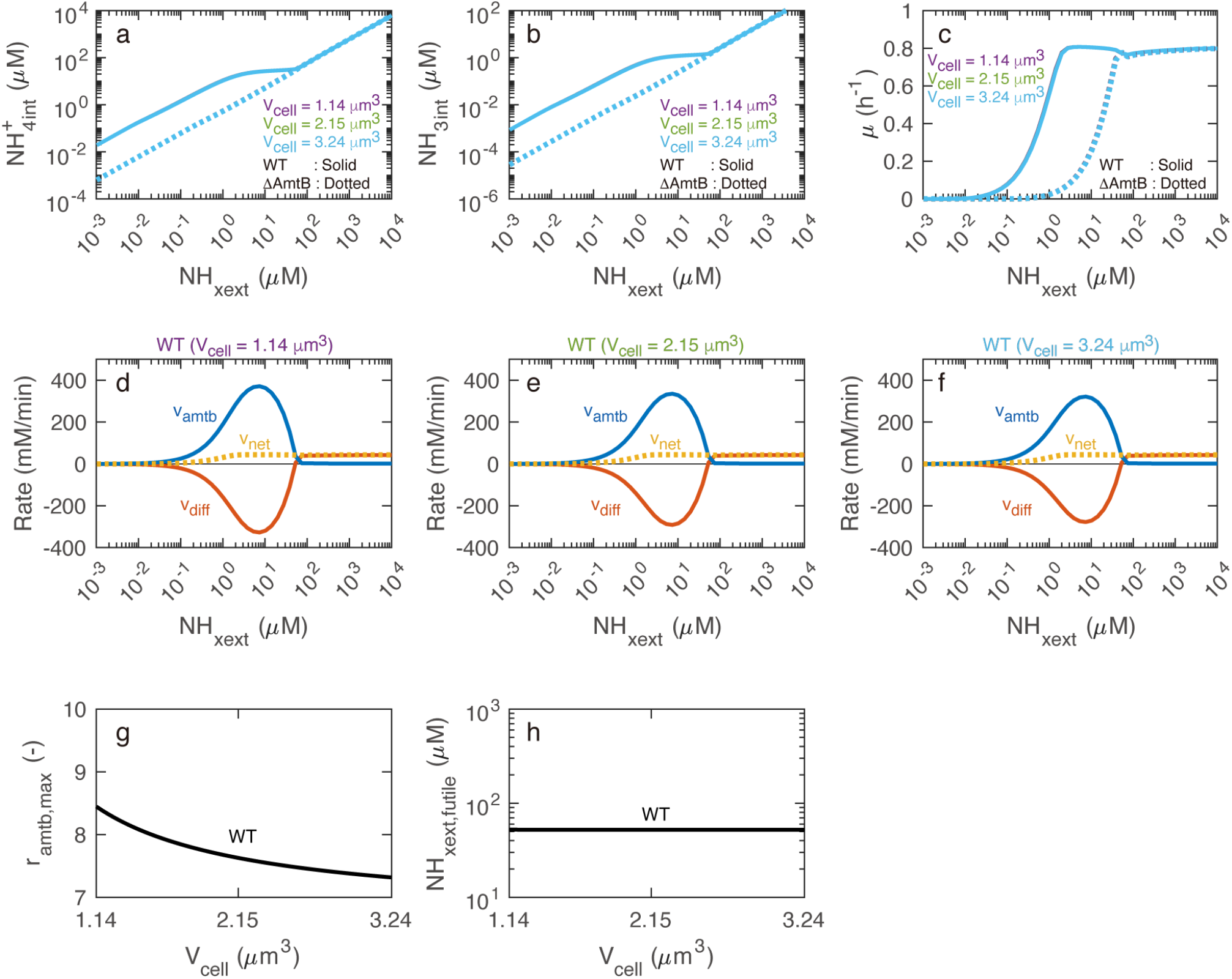
Effects of changes in cell volume on steady-state behaviors of the wild-type and ΔAmtB strains. (a) Intracellular NH_4_^+^ concentration. (b) Intracellular NH_3_ concentration. (c): Specific growth rate. (d–f) AmtB-mediated NH_4_^+^ transport (*v*_*amtb*_), unmediated NH_3_ diffusion (*v*_*diff*_), and net flux (*v*_*net*_) at cell volume of 1.14, 2.15, and 3.24 μm^3^. Positive flux values indicate the movement of NH_x_ from the extracellular environment into the cytoplasm. (g): The maximum values of *r*_*amtb*_ when the external NH_x_ was changed (*r*_*amtb,max*_). (h): The highest external NH_x_ concentration at which futile cycling occurs (*NH*_*xext,futile*_). In panels (a–c), the three solid lines coincide, and the three dotted lines also coincide.

**Fig. S9.**
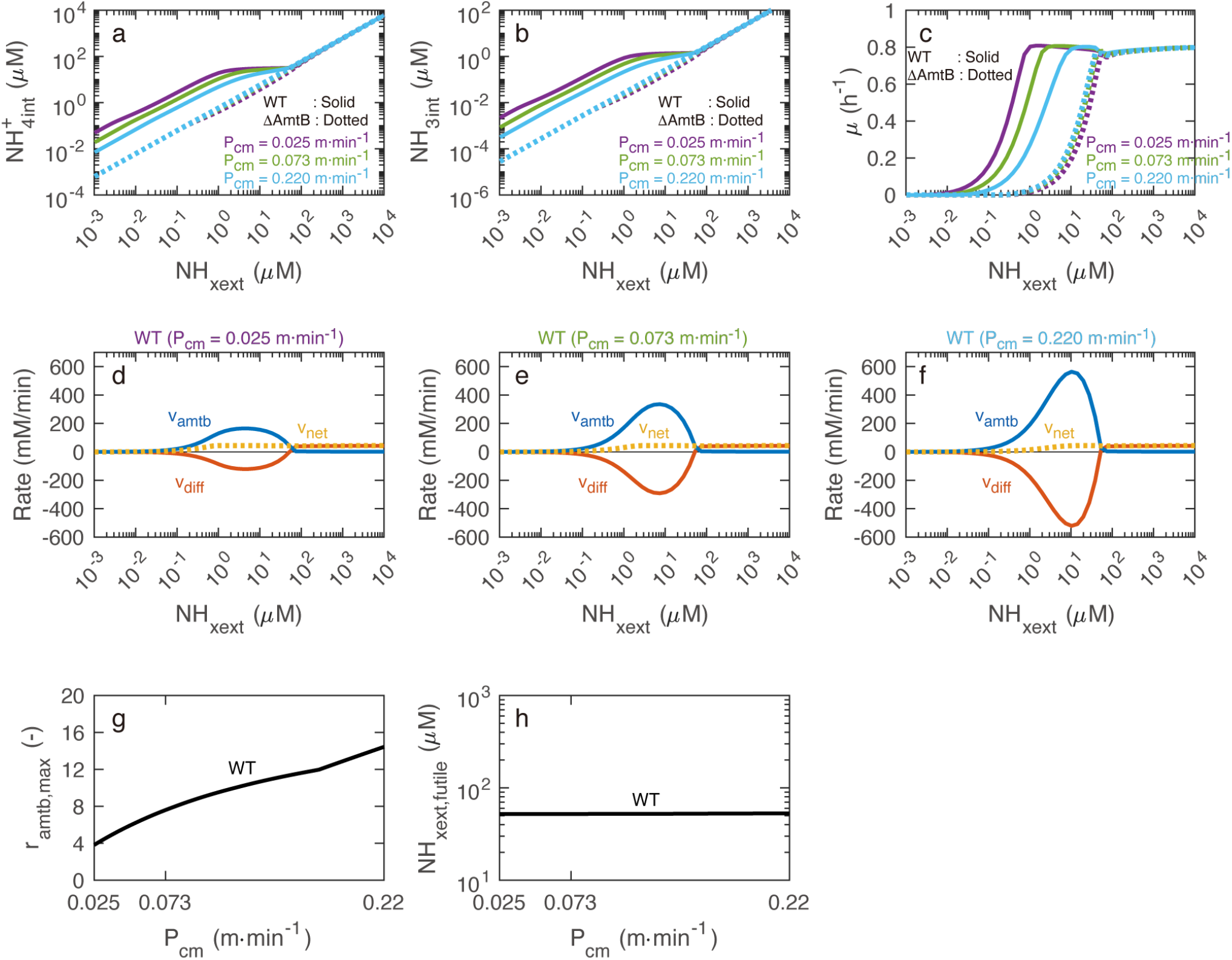
Effects of changes in NH_3_ permeability coefficient on steady-state behaviors of the wild-type and ΔAmtB strains. (a) Intracellular NH_4_^+^ concentration. (b) Intracellular NH_3_ concentration. (c) Specific growth rate. (d–f) AmtB-mediated NH_4_^+^ transport (*v*_*amtb*_), unmediated NH_3_ diffusion (*v*_*diff*_), and net flux (*v*_*net*_) at NH_3_ permeability coefficient of 0.025, 0.073, and 0.220 m min^-1^. Positive flux values indicate the movement of NH_x_ from the extracellular environment into the cytoplasm. (g) The maximum values of *r*_*amtb*_ when the external NH_x_ was changed (*r*_*amtb,max*_). (h) The highest external NH_x_ concentration at which futile cycling occurs (*NH*_*xext,futile*_). In panels (a–c), dotted lines mostly coincide.

**Fig. S10.**
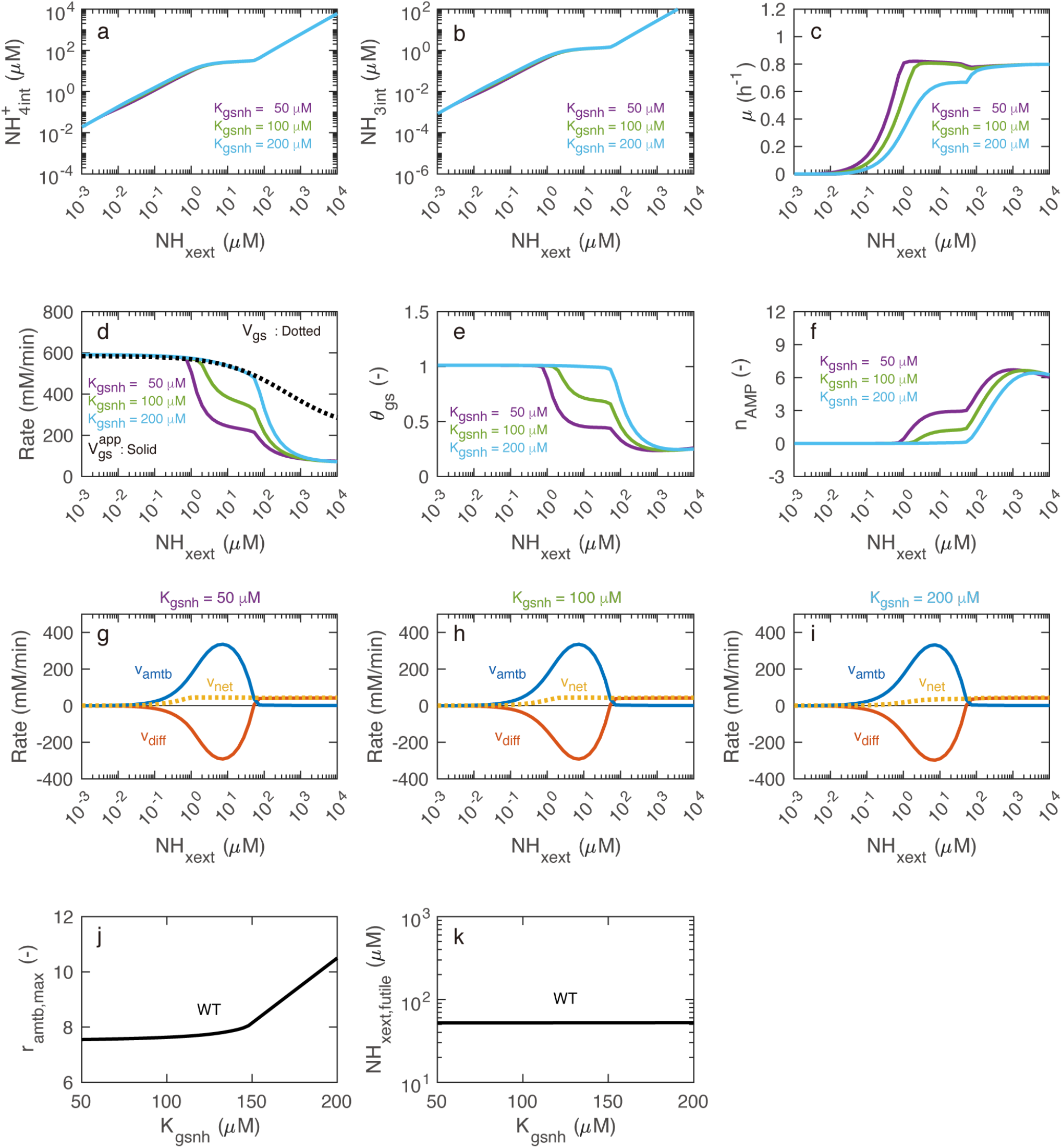
Effects of changes in Michaelis constant of GS for NH_4_^+^ on steady-state behaviors of the wild-type and ΔAmtB strains. (a) Intracellular NH_4_^+^ concentration. (b) Intracellular NH_3_ concentration. (c) Specific growth rate. (d) The apparent (*V*_*gs*_^*app*^) and true (*V*_*gs*_) maximum rates of the GS reaction. (e) Factor indicating the activity of GS, determined by the adenylylation state of GS. (f) Average number of adenylylated subunits in the GS dodecamer. (g–i) AmtB-mediated NH_4_^+^ transport (*v*_*amtb*_), unmediated NH_3_ diffusion (*v*_*diff*_), and net flux (*v*_*net*_) at Michaelis constant of 50, 100, and 200 μM. Positive flux values indicate the movement of NH_x_ from the extracellular environment into the cytoplasm. (j) The maximum values of *r*_*amtb*_ when the external NH_x_ was changed (*r*_*amtb,max*_). (k) The highest external NH_x_ concentration at which futile cycling occurs (*NH*_*xext,futile*_). In panels (a-b), all lines mostly coincide.

**Fig. S11.**
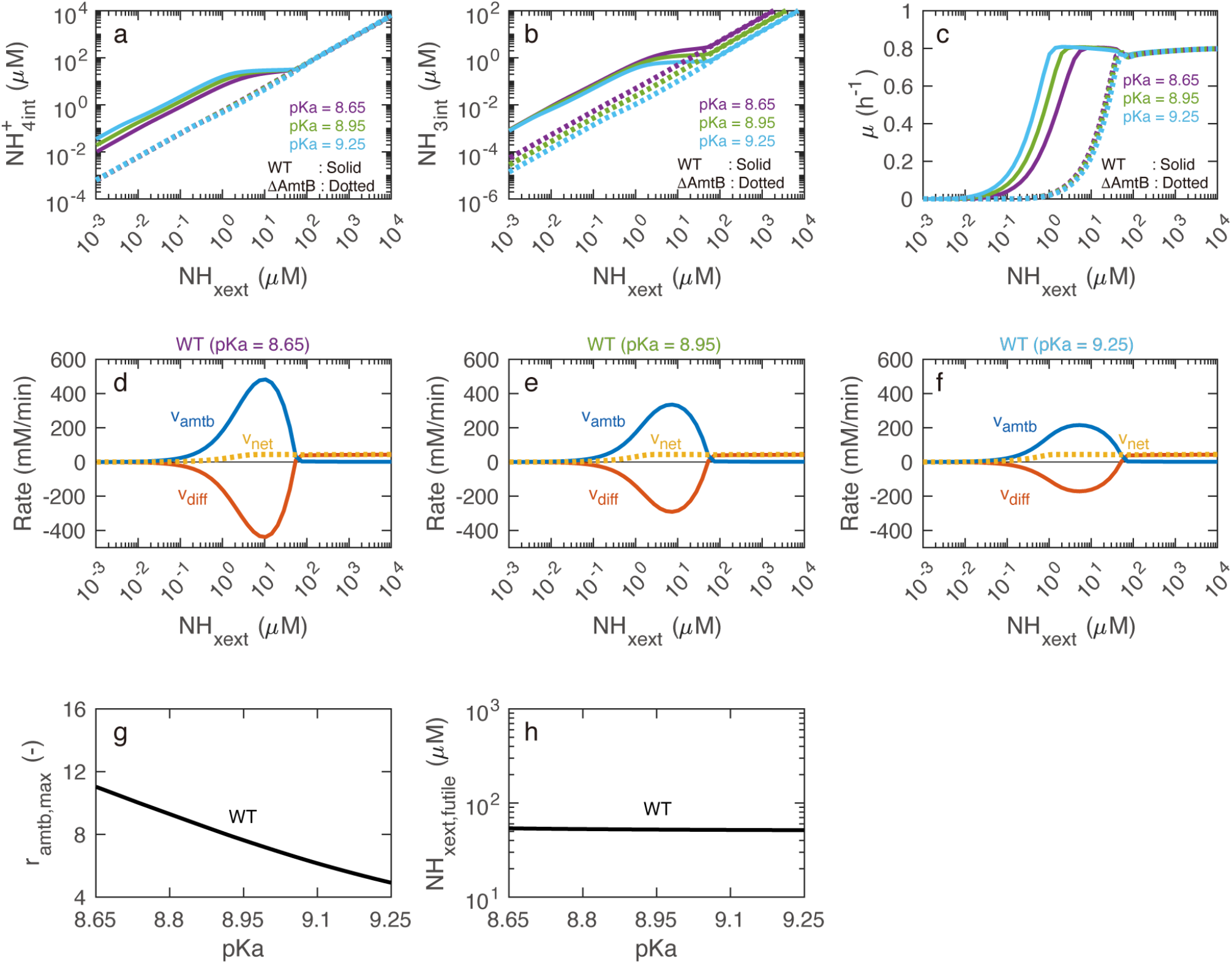
Effects of changes in pKa on steady-state behaviors of the wild-type and ΔAmtB strains. (a) Intracellular NH_4_^+^ concentration. (b) Intracellular NH_3_ concentration. (c) Specific growth rate. (d–f) AmtB-mediated NH_4_^+^ transport (*v*_*amtb*_), unmediated NH_3_ diffusion (*v*_*diff*_), and net flux (*v*_*net*_) at pKa of 8.65, 8.95, and 9.25. Positive flux values indicate the movement of NH_x_ from the extracellular environment into the cytoplasm. (g) The maximum values of *r*_*amtb*_ when the external NH_x_ was changed (*r*_*amtb,max*_). (h) The highest external NH_x_ concentration at which futile cycling occurs (*NH*_*xext,futile*_). In panels (a) and (c), dotted lines mostly coincide.

**Table S1.**
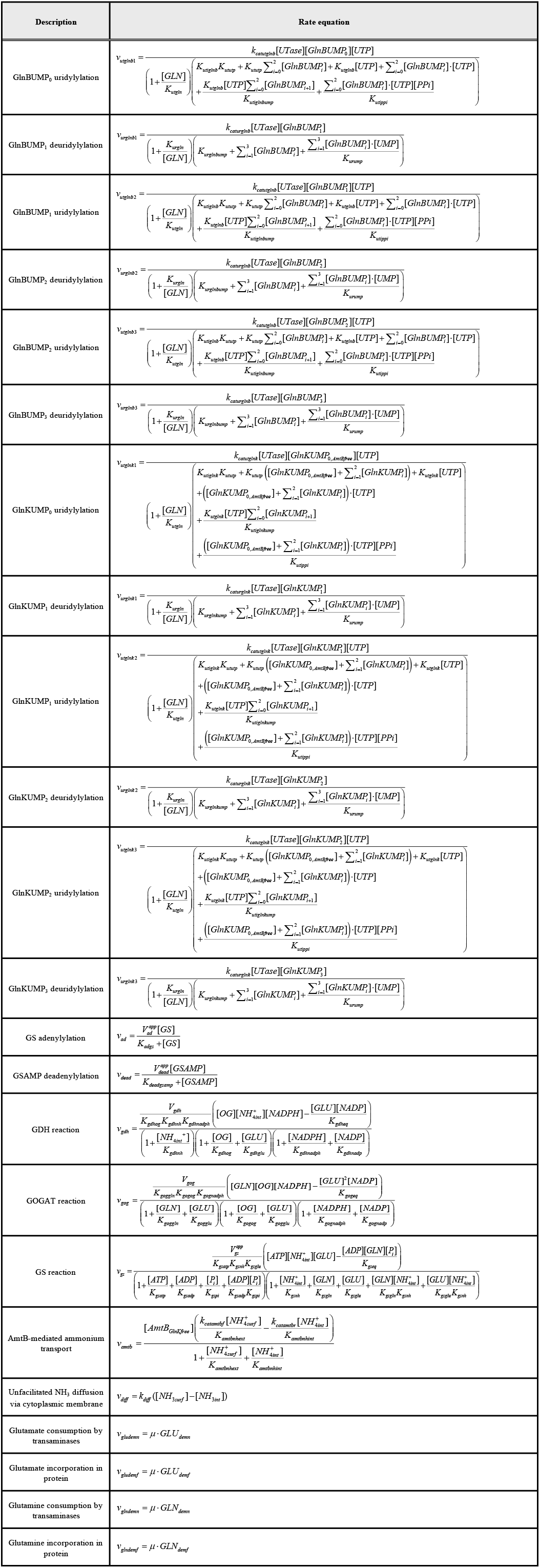
Rate equations of the models (*n* = 23)

**Table S2.** Description of the model variables. The variables (*n* = 13) are computed using the initial values and balance equations. The square brackets in the column “Variable” denote concentrations. The initial values for [GlnKUMP_0_] and [GS] are equal to [GlnK_total_] and [GS_total_], respectively, which are experiment-dependent parameters.

| Variable | Description | Initial value | Balance equation |
| --- | --- | --- | --- |
| $[GlnBUMP_0]$ | Trimer of the GlnB protein without UMP | 0.00065 mM [1] | $\frac{d[GlnBUMP_0]}{dt} = -v_{utglnb1} + v_{urglnb1}$ |
| $[GlnBUMP_1]$ | GlnB with one UMP group | 0 mM | $\frac{d[GlnBUMP_1]}{dt} = v_{utglnb1} - v_{urglnb1} - v_{utglnb2} + v_{urglnb2}$ |
| $[GlnBUMP_2]$ | GlnB with two UMP groups | 0 mM | $\frac{d[GlnBUMP_2]}{dt} = v_{utglnb2} - v_{urglnb2} - v_{utglnb3} + v_{urglnb3}$ |
| $[GlnBUMP_3]$ | GlnB with three UMP groups | 0 mM | $\frac{d[GlnBUMP_3]}{dt} = v_{utglnb3} - v_{urglnb3}$ |
| $[GlnKUMP_0]$ | Trimer of the GlnK protein without UMP | See legend | $\frac{d[GlnKUMP_0]}{dt} = -v_{utglnk1} + v_{urglnk1}$ |
| $[GlnKUMP_1]$ | GlnK with one UMP group | 0 mM | $\frac{d[GlnKUMP_1]}{dt} = v_{utglnk1} - v_{urglnk1} - v_{utglnk2} + v_{urglnk2}$ |
| $[GlnKUMP_2]$ | GlnK with two UMP groups | 0 mM | $\frac{d[GlnKUMP_2]}{dt} = v_{utglnk2} - v_{urglnk2} - v_{utglnk3} + v_{urglnk3}$ |
| $[GlnKUMP_3]$ | GlnK with three UMP groups | 0 mM | $\frac{d[GlnKUMP_3]}{dt} = v_{utglnk3} - v_{urglnk3}$ |
| $[GS]$ | Dodecamer of glutamine synthetase | See legend | $\frac{d[GS]}{dt} = -v_{ad} + v_{dead}$ |
| $[GSAMP]$ | Adenylylated GS | 0 mM | $\frac{d[GSAMP]}{dt} = v_{ad} - v_{dead}$ |
| $[GLU]$ | Glutamate | 80 mM [2] | $\frac{d[GLU]}{dt} = v_{gdh} + 2v_{gog} - v_{gs} - v_{gludemn} - v_{gludemf} + v_{glndemn}$ |
| $[GLN]$ | Glutamine | 2 mM [2] | $\frac{d[GLN]}{dt} = -v_{gog} + v_{gs} - v_{glndemn} - v_{glndemf}$ |
| $[NH_{x,int}]$ | Total of internal $NH_4^+$ and $NH_3$ | 0.02 mM [3] | $\frac{d[NH_{x,int}]}{dt} = v_{amtb} + v_{diff} - v_{gdh} - v_{gs}$ |

**Table S3.**
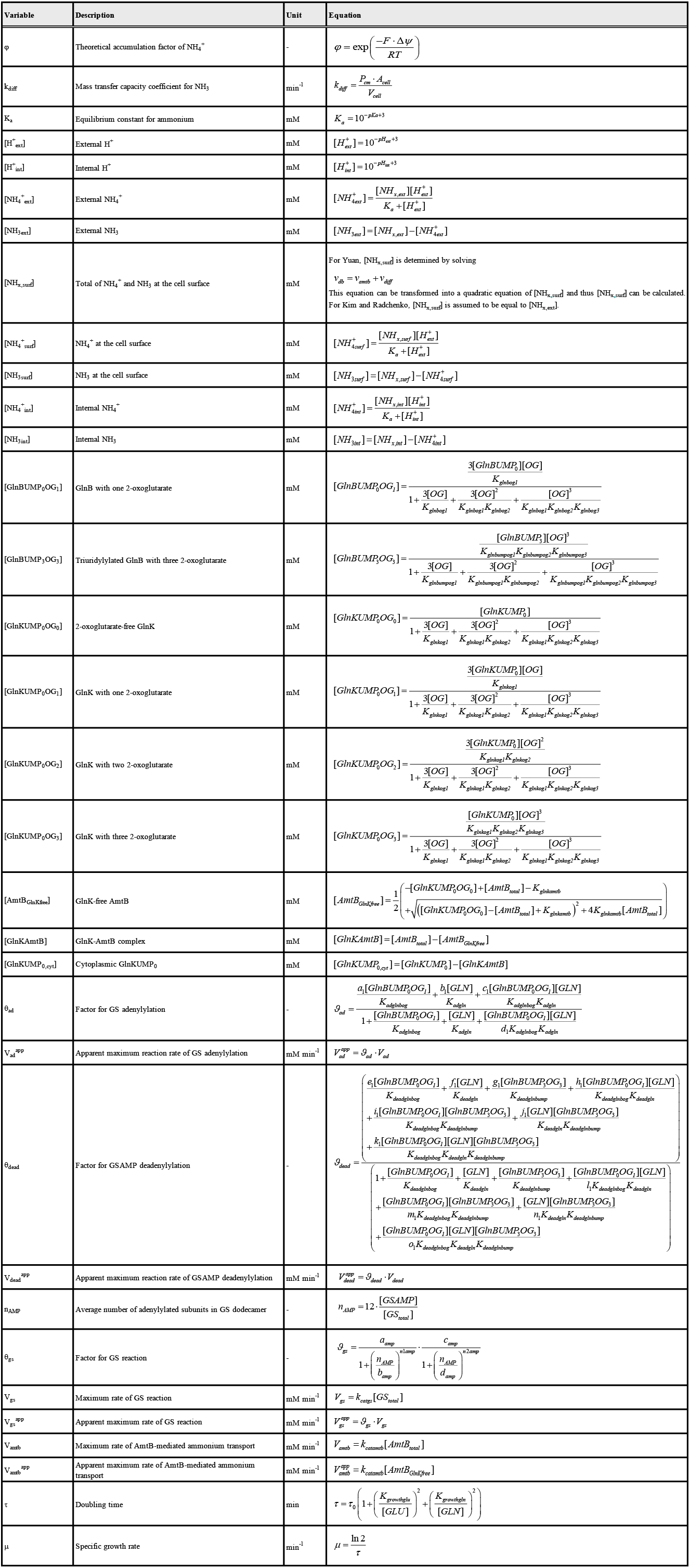
Description of the model auxiliary variables. The auxiliary variables (*n* = 33) are computed using the equations below.

**Table S4.**
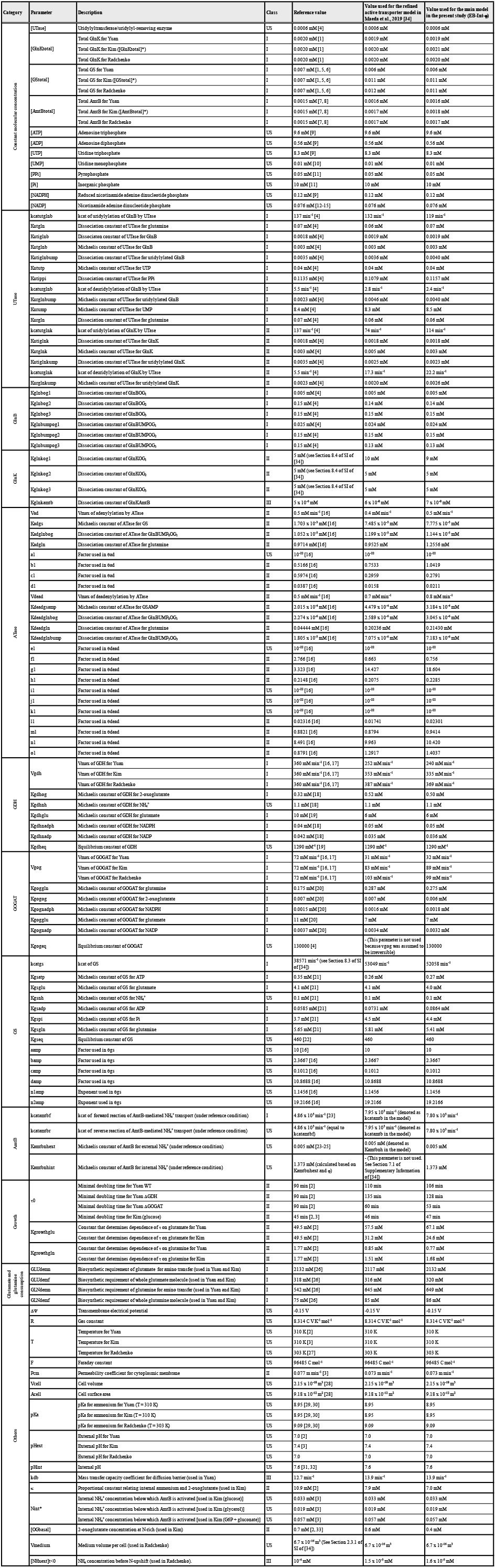
Description of the model parameters. The “Class” column shows parameter class (see the main text). We used class I (*n* = 52), class II (*n* = 39), and class III (*n* = 3) parameters. “US” in the class column indicates unsearched parameters in parameter estimation (*n* = 50). The parameter set shown in “Value used for the main model in the present study (EB-Int-*ρ*)” is the same as the parameter set highlighted in red in Table S6. Please note that, in this study, we mainly used parameters denoted as “Kim” and “glucose” for condition-dependent parameters. We used the parameters denoted as “Yuan” and “Radchenko” for parameter estimation and model validation (Figs. S2-S4). “Reference condition” for AmtB-related parameters indicates Δφ = -150 mV, T = 310 K, and thus *ρ* = 275.

**Table S5.**
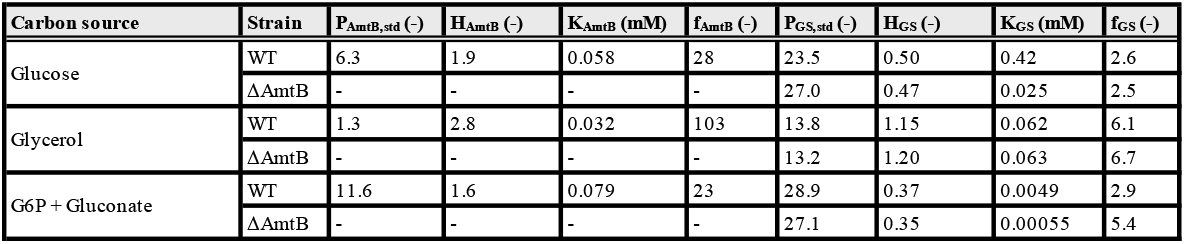
Parameters for calculation of AmtB, GlnK, and GS concentrations. For ΔAmtB, [AmtB_total_] = 0 and [GlnK_total_] = 0, and thus P_AmtB,std_, H_AmtB_, K_AmtB_, and f_AmtB_ are not provided for. Please note that, in this study, we mainly used parameters denoted as “Glucose” for condition-dependent parameters. We used the parameters denoted as “Glycerol” and “G6P+Gluconate” for model validation (Fig. S3). This table is identical to Table S8 in our previous study [34].

**Table S6.**
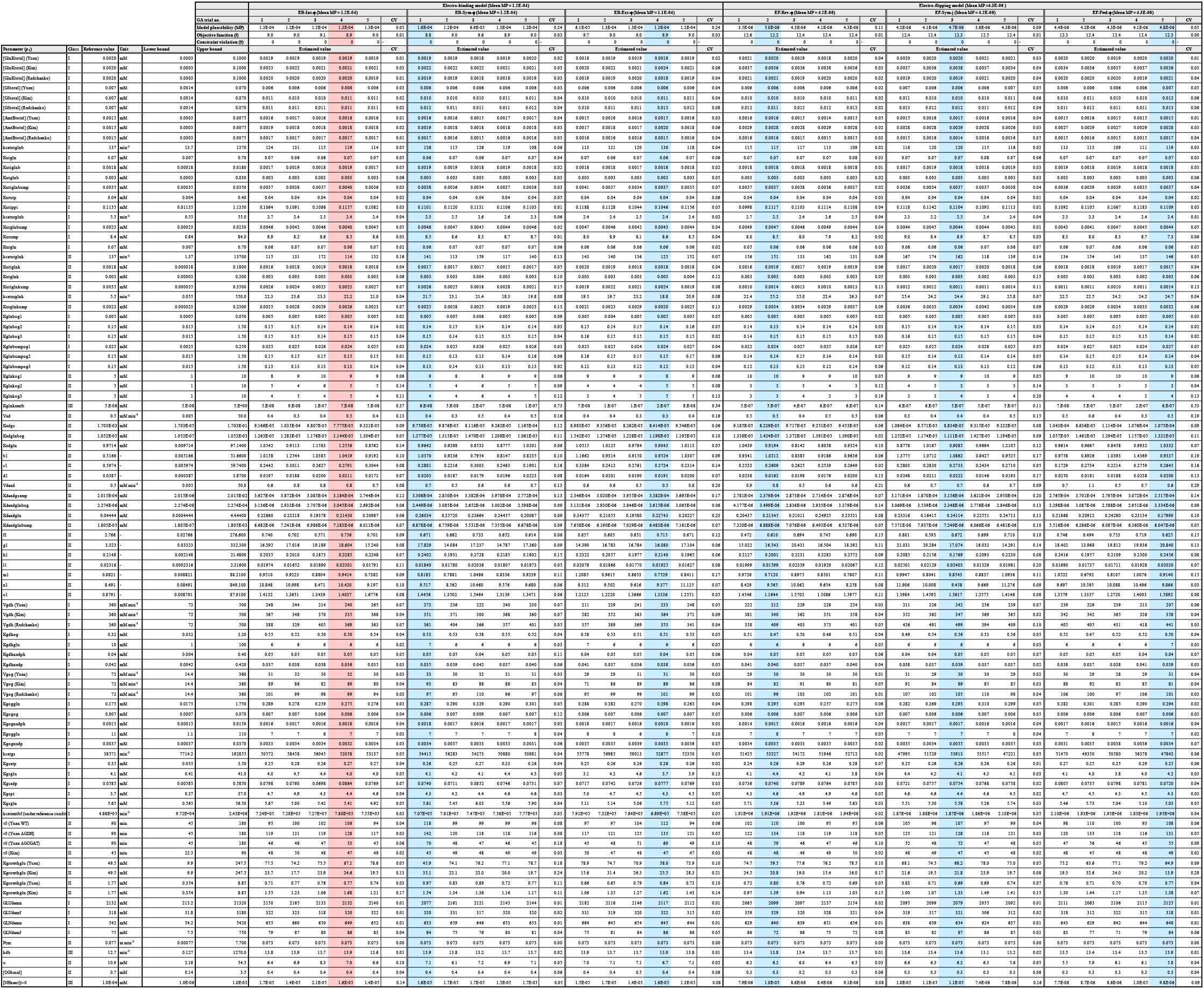
List of estimated parameter sets. Class I (*n* = 52), class II (*n* = 39), and class III (*n* = 3) search parameters are shown here. We performed five independent trials of genetic algorithms (GAs) for parameter estimation of each model variant. Since GAs are stochastic algorithms, estimated parameter values are different for each trial. The best parameter set for each model variant is highlighted in red or blue, and those are used for Figs 9 and S7. The best parameter set for the EB-Int-*φ* model is highlighted in red, which is used for analyzes in the present study, unless otherwise stated. All the 15 parameter sets (3 model variants x 5 trials) for each of the electro-binding and electro-flipping models are used to draw Fig. S5. “CV” columns show the coefficient of variation. “Reference condition” for *k*_*catamtbf*_ indicates Δψ = -150 mV, T = 310 K, and thus *φ* = 275.

**Table S7.**
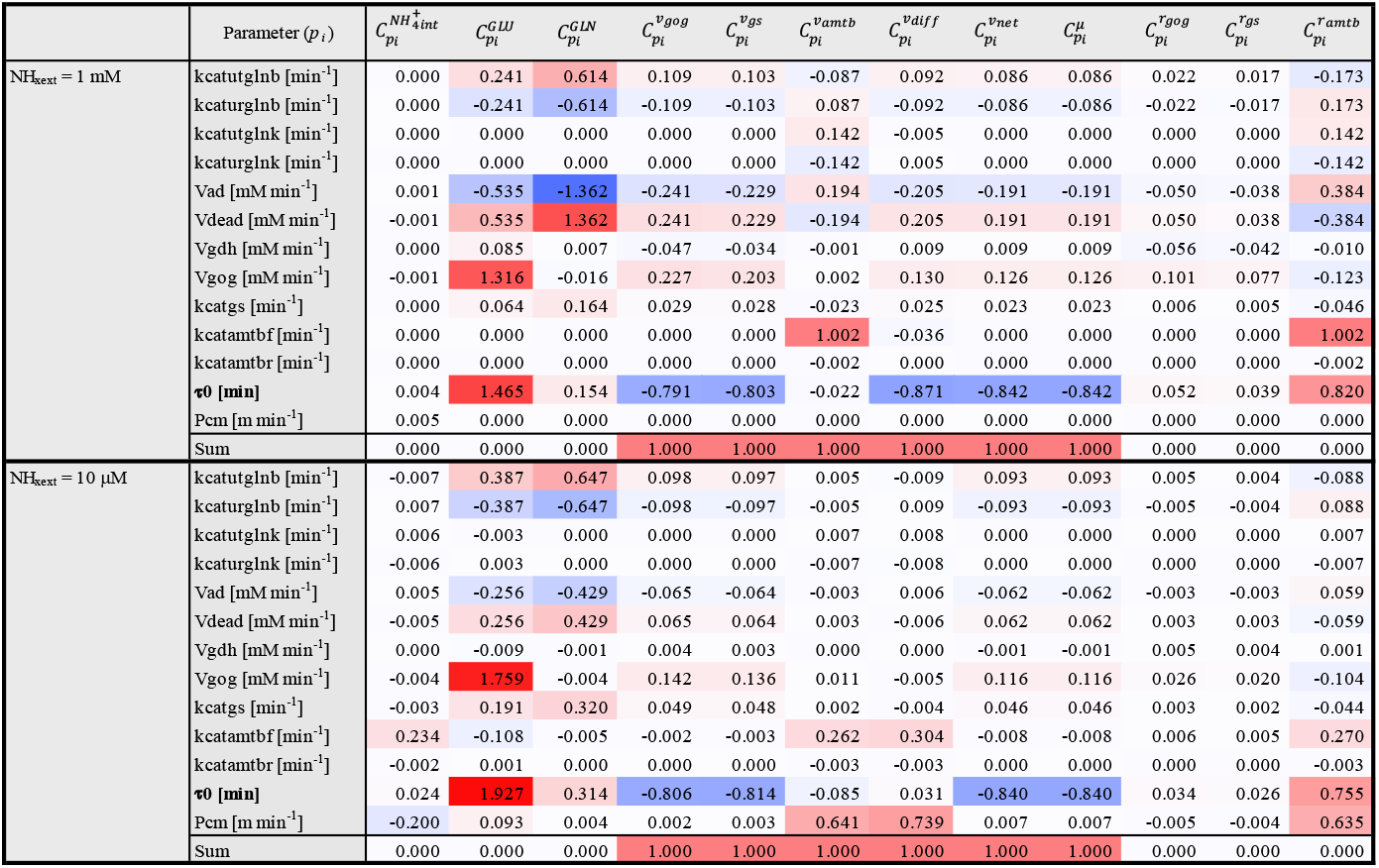
Control coefficients of important model variables. We calculated control coefficients with the EB-Int-*φ* model (the main model) with the red-highlighted parameter set in Table S6. The calculation was performed at the steady state in Kim’s experiment (WT, NH_xext_ = 1 mM and 10 μM) We numerically calculated control coefficients by slightly perturbing parameters (Δ*p*_*i*_ /*p*_*i*_ = 10^-6^). Positive and negative values are highlighted in red and blue, respectively, and color strength depends on the magnitude of control coefficients. Calculation of the row labeled “Sum”: control coefficients for parameters with dimensions of inverse time are summed directly, whereas **control coefficients for τ**_**0**_ **is summed with a negative sign because it has a dimension of time**. For model variables without time dimensions, the sum is zero, whereas for variables with dimensions of inverse time, the sum is one. This indicates that the summation law is satisfied.

## References

1. van Heeswijk, W. C., Westerhoff, H. V. & Boogerd, F. C. Nitrogen assimilation in Escherichia coli: putting molecular data into a systems perspective. Microbiol Mol Biol Rev. 77, 628–695 (2013).

2. Kleiner, D. The transport of NH3 and NH4+ across biological membranes. Biochimica et biophysica acta. 639, 41–52 (1981).

3. Kleiner, D. Bacterial Ammonium Transport. FEMS Microbiology Letters. 32, 87–100 (1985).

4. Soupene, E., He, L., Yan, D. & Kustu, S. Ammonia acquisition in enteric bacteria: physiological role of the ammonium/methylammonium transport B (AmtB) protein. Proc Natl Acad Sci U S A. 95, 7030–7034 (1998).

5. Winkler, F. K. Amt/MEP/Rh proteins conduct ammonia. Pflugers Arch. 451, 701–707 (2006).

6. Javelle, A., et al. Structural and mechanistic aspects of Amt/Rh proteins. Journal of structural biology. 158, 472–481 (2007).

7. Andrade, S. L. & Einsle, O. The Amt/Mep/Rh family of ammonium transport proteins. Molecular membrane biology. 24, 357–365 (2007).

8. Neuhauser, B., Dynowski, M. & Ludewig, U. Switching substrate specificity of AMT/MEP/Rh proteins. Channels (Austin). 8, 496–502 (2014).

9. Bizior, A., Williamson, G., Harris, T., Hoskisson, P. A. & Javelle, A. Prokaryotic ammonium transporters: what has three decades of research revealed? Microbiology. 169 (2023).

10. Williamson, G., et al. Biological ammonium transporters from the Amt/Mep/Rh superfamily: mechanism, energetics, and technical limitations. Biosci Rep. 44 (2024).

11. Williamson, G., et al. Biological ammonium transporters: evolution and diversification. FEBS J. 291, 3786–3810 (2024).

12. Boogerd, F. C., et al. AmtB-mediated NH3 transport in prokaryotes must be active and as a consequence regulation of transport by GlnK is mandatory to limit futile cycling of NH4(+)/NH3. FEBS Lett. 585, 23–28 (2011).

13. Meek, T. D. & Villafranca, J. J. Kinetic mechanism of Escherichia coli glutamine synthetase. Biochemistry. 19, 5513–5519 (1980).

14. Maeda, K., Westerhoff, H. V., Kurata, H. & Boogerd, F. C. Ranking network mechanisms by how they fit diverse experiments and deciding on E. coli’s ammonium transport and assimilation network. NPJ Syst Biol Appl. 5, 14 (2019).

15. Mirandela, G. D., Tamburrino, G., Hoskisson, P. A., Zachariae, U. & Javelle, A. The lipid environment determines the activity of the Escherichia coli ammonium transporter AmtB. FASEB J. 33, 1989–1999 (2019).

16. Williamson, G., et al. A two-lane mechanism for selective biological ammonium transport. Elife. 9 (2020).

17. van Heeswijk, W. C., et al. An additional PII in Escherichia coli: a new regulatory protein in the glutamine synthetase cascade. FEMS Microbiol Lett. 132, 153–157 (1995).

18. Thomas, G., Coutts, G. & Merrick, M. The glnKamtB operon. A conserved gene pair in prokaryotes. Trends Genet. 16, 11–14 (2000).

19. Radchenko, M. V., Thornton, J. & Merrick, M. Control of AmtB-GlnK complex formation by intracellular levels of ATP, ADP, and 2-oxoglutarate. J Biol Chem. 285, 31037–31045 (2010).

20. Radchenko, M. V., Thornton, J. & Merrick, M. P(II) signal transduction proteins are ATPases whose activity is regulated by 2-oxoglutarate. Proc Natl Acad Sci U S A. 110, 12948–12953 (2013).

21. Radchenko, M. V., Thornton, J. & Merrick, M. Association and dissociation of the GlnK-AmtB complex in response to cellular nitrogen status can occur in the absence of GlnK post-translational modification. Frontiers in microbiology. 5, 731 (2014).

22. Kim, M., et al. Need-based activation of ammonium uptake in Escherichia coli. Mol Syst Biol. 8, 616 (2012).

23. Yuan, J., et al. Metabolomics-driven quantitative analysis of ammonia assimilation in E. coli. Mol Syst Biol. 5, 302 (2009).

24. Kurata, H., Matoba, N. & Shimizu, N. CADLIVE for constructing a large-scale biochemical network based on a simulation-directed notation and its application to yeast cell cycle. Nucleic Acids Res. 31, 4071–4084 (2003).

25. Kurata, H., Masaki, K., Sumida, Y. & Iwasaki, R. CADLIVE dynamic simulator: direct link of biochemical networks to dynamic models. Genome Res. 15, 590–600 (2005).

26. Kurata, H., et al. Extended CADLIVE: a novel graphical notation for design of biochemical network maps and computational pathway analysis. Nucleic Acids Res. 35, e134 (2007).

27. Inoue, K., et al. CADLIVE optimizer: web-based parameter estimation for dynamic models. Source code for biology and medicine. 7, 9 (2012).

28. Inoue, K., Maeda, K., Miyabe, T., Matsuoka, Y. & Kurata, H. CADLIVE toolbox for MATLAB: automatic dynamic modeling of biochemical networks with comprehensive system analysis. Bioprocess Biosyst Eng. 37, 1925–1927 (2014).

29. Javelle, A., Severi, E., Thornton, J. & Merrick, M. Ammonium sensing in Escherichia coli. Role of the ammonium transporter AmtB and AmtB-GlnK complex formation. J Biol Chem. 279, 8530–8538 (2004).

30. Belliveau, N. M., et al. Fundamental limits on the rate of bacterial growth and their influence on proteomic composition. Cell Syst. 12, 924–944 e922 (2021).

31. Westerhoff, H. V. Summation Laws in Control of Biochemical Systems. Mathematics. 11, 2473 (2023).

32. Kahn, D. & Westerhoff, H. V. Control theory of regulatory cascades. J Theor Biol. 153, 255–285 (1991).

33. Westerhoff, H. V. Signalling control strength. J Theor Biol. 252, 555–567 (2008).

34. Slonczewski, J. L., Fujisawa, M., Dopson, M. & Krulwich, T. A. Cytoplasmic pH measurement and homeostasis in bacteria and archaea. Adv Microb Physiol. 55, 1-79, 317 (2009).

35. Reyes-Fernandez, E. Z. & Schuldiner, S. Acidification of Cytoplasm in Escherichia coli Provides a Strategy to Cope with Stress and Facilitates Development of Antibiotic Resistance. Scientific reports. 10, 9954 (2020).

36. Felle, H., Porter, J. S., Slayman, C. L. & Kaback, H. R. Quantitative measurements of membrane potential in Escherichia coli. Biochemistry. 19, 3585–3590 (1980).

37. Tran, Q. H. & Unden, G. Changes in the proton potential and the cellular energetics of Escherichia coli during growth by aerobic and anaerobic respiration or by fermentation. Eur J Biochem. 251, 538–543 (1998).

38. Bot, C. T. & Prodan, C. Quantifying the membrane potential during E. coli growth stages. Biophys Chem. 146, 133–137 (2010).

39. Westerhoff, H. V. & van Dam, K. Thermodynamics and control of biological free-energy transduction (Elsevier, 1987).

40. Javelle, A., et al. Substrate binding, deprotonation, and selectivity at the periplasmic entrance of the Escherichia coli ammonia channel AmtB. Proc Natl Acad Sci U S A. 105, 5040–5045 (2008).

41. Ariz, I., et al. Nitrogen isotope signature evidences ammonium deprotonation as a common transport mechanism for the AMT-Mep-Rh protein superfamily. Sci Adv. 4, eaar3599 (2018).

42. Wang, J., et al. Ammonium transport proteins with changes in one of the conserved pore histidines have different performance in ammonia and methylamine conduction. PLoS One. 8, e62745 (2013).

43. Javelle, A., Thomas, G., Marini, A. M., Kramer, R. & Merrick, M. In vivo functional characterization of the Escherichia coli ammonium channel AmtB: evidence for metabolic coupling of AmtB to glutamine synthetase. Biochem J. 390, 215–222 (2005).

44. Jonker, C. M., Snoep, J. L., Treur, J., Westerhoff, H. V. & Wijngaards, W. C. Putting intentions into cell biochemistry: an artificial intelligence perspective. J Theor Biol. 214, 105–134 (2002).

45. Jonker, C. M., Snoep, J. L., Treur, J., Westerhoff, H. V. & Wijngaards, W. C. BDI-modelling of complex intracellular dynamics. J Theor Biol. 251, 1–23 (2008).

46. Westerhoff, H. V., et al. Macromolecular networks and intelligence in microorganisms. Frontiers in microbiology. 5, 379 (2014).

47. Chang, A., et al. BRENDA, the ELIXIR core data resource in 2021: new developments and updates. Nucleic Acids Res. 49, D498–D508 (2021).

48. Orr, J. & Haselkorn, R. Kinetic and inhibition studies of glutamine synthetase from the cyanobacterium Anabaena 7120. J Biol Chem. 256, 13099–13104 (1981).

49. Duchars, M. G. & Attwood, M. M. Purification, localization, properties and regulation of glutamine synthetase from Hyphomicrobium X. Journal of General Microbiology. 137, 1345–1354 (1991).

50. Krulwich, T. A., Sachs, G. & Padan, E. Molecular aspects of bacterial pH sensing and homeostasis. Nat Rev Microbiol. 9, 330–343 (2011).

51. Chisholm, S. W. Prochlorococcus. Curr Biol. 27, R447–R448 (2017).

52. Bolay, P., Muro-Pastor, M. I., Florencio, F. J. & Klahn, S. The Distinctive Regulation of Cyanobacterial Glutamine Synthetase. Life (Basel). 8 (2018).

53. El Alaoui, S., et al. Glutamine synthetase from the marine cyanobacteria Prochlorococcus spp: characterization, phylogeny and response to nutrient limitation. Environ Microbiol. 5, 412–423 (2003).

54. Zubkov, M. V. Faster growth of the major prokaryotic versus eukaryotic CO2 fixers in the oligotrophic ocean. Nature communications. 5, 3776 (2014).

55. ter Kuile, B. H. & Westerhoff, H. V. Transcriptome meets metabolome: hierarchical and metabolic regulation of the glycolytic pathway. FEBS Lett. 500, 169–171 (2001).

56. Maeda, K., Boogerd, F. C. & Kurata, H. libRCGA: a C library for real-coded genetic algorithms for rapid parameter estimation of kinetic models. IPSJ Transactions on Bioinformatics. 11, 31–40 (2018).

57. Maeda, K., Boogerd, F. C. & Kurata, H. RCGAToolbox: A Real-coded Genetic Algorithm Software for Parameter Estimation of Kinetic Models. IPSJ Transactions on Bioinformatics. 14, 30–35 (2021).

58. Bruggeman, F. J., Boogerd, F. C. & Westerhoff, H. V. The multifarious short-term regulation of ammonium assimilation of Escherichia coli: dissection using an in silico replica. FEBS J. 272, 1965–1985 (2005).

59. Cayley, S., Lewis, B. A., Guttman, H. J. & Record, M. T. Characterization of the cytoplasm of Escherichia coli K-12 as a function of external osmolarity. Journal of Molecular Biology. 222, 281–300 (1991).

60. Folsom, J. P. & Carlson, R. P. Physiological, biomass elemental composition and proteomic analyses of Escherichia coli ammonium-limited chemostat growth, and comparison with iron- and glucose-limited chemostat growth. Microbiology. 161, 1659–1670 (2015).

## References

1. Kim M, Zhang Z, Okano H, Yan D, Groisman A, Hwa T. Need-based activation of ammonium uptake in Escherichia coli. Mol Syst Biol. 2012;8:616.

2. Maeda K, Westerhoff HV, Kurata H, Boogerd FC. Ranking network mechanisms by how they fit diverse experiments and deciding on E. coli’s ammonium transport and assimilation network. NPJ Syst Biol Appl. 2019;5(1):14.

3. Yuan J, Doucette CD, Fowler WU, Feng XJ, Piazza M, Rabitz HA, et al. Metabolomics-driven quantitative analysis of ammonia assimilation in E. coli. Mol Syst Biol. 2009;5:302.

4. Yan D, Lenz P, Hwa T. Overcoming fluctuation and leakage problems in the quantification of intracellular 2-oxoglutarate levels in Escherichia coli. Applied and environmental microbiology. 2011;77(19):6763–71.

5. Javelle A, Severi E, Thornton J, Merrick M. Ammonium sensing in Escherichia coli. Role of the ammonium transporter AmtB and AmtB-GlnK complex formation. J Biol Chem. 2004;279(10):8530–8.

6. Javelle A, Lupo D, Li XD, Merrick M, Chami M, Ripoche P, et al. Structural and mechanistic aspects of Amt/Rh proteins. Journal of structural biology. 2007;158(3):472–81.

7. Kleiner D. Bacterial Ammonium Transport. FEMS Microbiology Letters. 1985;32(2):87–100.

8. Radchenko MV, Thornton J, Merrick M. Control of AmtB-GlnK complex formation by intracellular levels of ATP, ADP, and 2-oxoglutarate. J Biol Chem. 2010;285(40):31037–45.

9. Belliveau NM, Chure G, Hueschen CL, Garcia HG, Kondev J, Fisher DS, et al. Fundamental limits on the rate of bacterial growth and their influence on proteomic composition. Cell Syst. 2021;12(9):924–44 e2.

10. van Heeswijk WC, Molenaar D, Hoving S, Westerhoff HV. The pivotal regulator GlnB of Escherichia coli is engaged in subtle and context-dependent control. FEBS J. 2009;276(12):3324–40.

11. Gosztolai A, Schumacher J, Behrends V, Bundy JG, Heydenreich F, Bennett MH, et al. GlnK Facilitates the Dynamic Regulation of Bacterial Nitrogen Assimilation. Biophys J. 2017;112(10):2219–30.

12. Khademi S, O’Connell J, 3rd, Remis J, Robles-Colmenares Y, Miercke LJ, Stroud RM. Mechanism of ammonia transport by Amt/MEP/Rh: structure of AmtB at 1.35 A. Science. 2004;305(5690):1587–94.

13. Zheng L, Kostrewa D, Berneche S, Winkler FK, Li XD. The mechanism of ammonia transport based on the crystal structure of AmtB of Escherichia coli. Proc Natl Acad Sci U S A. 2004;101(49):17090–5.

14. Radzikowski JL, Vedelaar S, Siegel D, Ortega AD, Schmidt A, Heinemann M. Bacterial persistence is an active sigmaS stress response to metabolic flux limitation. Mol Syst Biol. 2016;12(9):882.

15. Volkmer B, Heinemann M. Condition-dependent cell volume and concentration of Escherichia coli to facilitate data conversion for systems biology modeling. PLoS One. 2011;6(7):e23126.

16. Walter A, Gutknecht J. Permeability of small nonelectrolytes through lipid bilayer membranes. The Journal of membrane biology. 1986;90(3):207–17.

17. Meek TD, Villafranca JJ. Kinetic mechanism of Escherichia coli glutamine synthetase. Biochemistry. 1980;19(24):5513–9.

## References

1. van Heeswijk, W. C., Molenaar, D., Hoving, S. & Westerhoff, H. V. (2009) The pivotal regulator GlnB of Escherichia coli is engaged in subtle and context-dependent control, FEBS J. 276, 3324–40.

2. Yuan, J., Doucette, C. D., Fowler, W. U., Feng, X. J., Piazza, M., Rabitz, H. A., Wingreen, N. S. & Rabinowitz, J. D. (2009) Metabolomics-driven quantitative analysis of ammonia assimilation in E. coli, Mol Syst Biol. 5, 302.

3. Kim, M., Zhang, Z., Okano, H., Yan, D., Groisman, A. & Hwa, T. (2012) Need-based activation of ammonium uptake in Escherichia coli, Mol Syst Biol. 8, 616.

4. Jiang, P., Peliska, J. A. & Ninfa, A. J. (1998) Enzymological characterization of the signal-transducing uridylyltransferase/uridylyl-removing enzyme (EC 2.7.7.59) of Escherichia coli and its interaction with the PII protein, Biochemistry. 37, 12782–94.

5. Heeswijk, W. C. (1998) The glutamine synthetase adenylylation cascade: A search for its control and regulation, Vrije Universiteit, Amsterdam, the Netherlands.

6. Senior, P. J. (1975) Regulation of nitrogen metabolism in Escherichia coli and Klebsiella aerogenes: studies with the continuous-culture technique, J Bacteriol. 123, 407–18.

7. Blauwkamp, T. A. & Ninfa, A. J. (2003) Antagonism of PII signalling by the AmtB protein of Escherichia coli, Mol Microbiol. 48, 1017–28.

8. Zheng, L., Kostrewa, D., Berneche, S., Winkler, F. K. & Li, X. D. (2004) The mechanism of ammonia transport based on the crystal structure of AmtB of Escherichia coli, Proc Natl Acad Sci U S A. 101, 17090–5.

9. Bennett, B. D., Kimball, E. H., Gao, M., Osterhout, R., Van Dien, S. J. & Rabinowitz, J. D. (2009) Absolute metabolite concentrations and implied enzyme active site occupancy in Escherichia coli, Nature chemical biology. 5, 593–9.

10. Neidhardt, F. C. & Umbarger, H. E. (1996) Chemical composition of Escherichia coli in Escherichia coli and Salmonella: cellular and molecular biology (Neidhardt, F. C., ed) pp. 13–17, ACM Press, Washington (D. C.).

11. Wanner, B. L. (1996) Phosphorus assimilation and control of the phosphate regulon in Escherichia coli and Salmonella: cellular and molecular biology (Neidhardt, F. C., ed) pp. 1357–1381, ASM Press, Washington (D. C.).

12. Penfound, T. & Foster, J. W. (1996) Biosynthesis and recycling of NAD in Escherichia coli and Salmonella: cellular and molecular biology (Neidhardt, F. C., ed) pp. 721–730, ASM Press, Washington (D. C.).

13. Andersen, K. B. & von Meyenburg, K. (1977) Charges of nicotinamide adenine nucleotides and adenylate energy charge as regulatory parameters of the metabolism in Escherichia coli, J Biol Chem. 252, 4151–6.

14. Fuhrer, T. & Sauer, U. (2009) Different biochemical mechanisms ensure network-wide balancing of reducing equivalents in microbial metabolism, J Bacteriol. 191, 2112–21.

15. Auriol, C., Bestel-Corre, G., Claude, J. B., Soucaille, P. & Meynial-Salles, I. (2011) Stress-induced evolution of Escherichia coli points to original concepts in respiratory cofactor selectivity, Proc Natl Acad Sci U S A. 108, 1278–83.

16. Bruggeman, F. J., Boogerd, F. C. & Westerhoff, H. V. (2005) The multifarious short-term regulation of ammonium assimilation of Escherichia coli: dissection using an in silico replica, FEBS J. 272, 1965–85.

17. Garcia-Contreras, R., Vos, P., Westerhoff, H. V. & Boogerd, F. C. (2012) Why in vivo may not equal in vitro - new effectors revealed by measurement of enzymatic activities under the same in vivo-like assay conditions, FEBS J. 279, 4145–59.

18. Sakamoto, N., Kotre, A. M. & Savageau, M. A. (1975) Glutamate dehydrogenase from Escherichia coli: purification and properties, J Bacteriol. 124, 775–83.

19. Thermodynamics of Enzyme-Catalyzed Reactions [http://xpdb.nist.gov/enzyme_thermodynamics/enzyme_thermodynamics.html]

20. Rendina, A. R. & Orme-Johnson, W. H. (1978) Glutamate synthase: on the kinetic mechanism of the enzyme from Escherichia coli W, Biochemistry. 17, 5388–93.

21. Meek, T. D. & Villafranca, J. J. (1980) Kinetic mechanism of Escherichia coli glutamine synthetase, Biochemistry. 19, 5513–9.

22. Rhee, S. G., Chock, P. B. & Stadtman, E. R. (1985) Glutamine synthetase from Escherichia coli, Methods in enzymology. 113, 213–41.

23. Javelle, A., Lupo, D., Li, X. D., Merrick, M., Chami, M., Ripoche, P. & Winkler, F. K. (2007) Structural and mechanistic aspects of Amt/Rh proteins, Journal of structural biology. 158, 472–81.

24. Kleiner, D. (1985) Bacterial Ammonium Transport, FEMS Microbiology Letters. 32, 87–100.

25. Javelle, A., Severi, E., Thornton, J. & Merrick, M. (2004) Ammonium sensing in Escherichia coli. Role of the ammonium transporter AmtB and AmtB-GlnK complex formation, J Biol Chem. 279, 8530–8.

26. Reitzer, L. (2003) Nitrogen assimilation and global regulation in Escherichia coli, Annu Rev Microbiol. 57, 155–76.

27. Radchenko, M. V., Thornton, J. & Merrick, M. (2014) Association and dissociation of the GlnK-AmtB complex in response to cellular nitrogen status can occur in the absence of GlnK post-translational modification, Frontiers in microbiology. 5, 731.

28. Radzikowski, J. L., Vedelaar, S., Siegel, D., Ortega, A. D., Schmidt, A. & Heinemann, M. (2016) Bacterial persistence is an active sigmaS stress response to metabolic flux limitation, Mol Syst Biol. 12, 882.

29. Martinelle, K. & Haggstrom, L. (1997) On the dissociation constant of ammonium: Effects of using an incorrect pK(a) in calculations of the ammonia concentration in animal cell cultures, Biotechnology Techniques. 11, 549–551.

30. Lang, W., Block, T. M. & Zander, R. (1998) Solubility of NH3 and apparent pK of NH4+ in human plasma, isotonic salt solutions and water at 37 degrees C, Clinica chimica acta; international journal of clinical chemistry. 273, 43–58.

31. Slonczewski, J. L., Rosen, B. P., Alger, J. R. & Macnab, R. M. (1981) pH homeostasis in Escherichia coli: measurement by 31P nuclear magnetic resonance of methylphosphonate and phosphate, Proc Natl Acad Sci U S A. 78, 6271–5.

32. Slonczewski, J. L., Fujisawa, M., Dopson, M. & Krulwich, T. A. (2009) Cytoplasmic pH measurement and homeostasis in bacteria and archaea, Adv Microb Physiol. 55, 1-79, 317.

33. Yan, D., Lenz, P. & Hwa, T. (2011) Overcoming fluctuation and leakage problems in the quantification of intracellular 2-oxoglutarate levels in Escherichia coli, Applied and environmental microbiology. 77, 6763–71.

34. Maeda, K., Westerhoff, H. V., Kurata, H. & Boogerd, F. C. (2019) Ranking network mechanisms by how they fit diverse experiments and deciding on E. coli’s ammonium transport and assimilation network, NPJ Syst Biol Appl. 5, 14.

